# Fungal Communities in Soybean Cyst Nematode-Infested Soil under Long Term Corn and Soybean Monoculture and Crop Rotation

**DOI:** 10.1101/516575

**Authors:** Noah Strom, Weiming Hu, Deepak Haarith, Senyu Chen, Kathryn Bushley

**Author notes:** **Correspondence:** Kathryn Bushley.

## Abstract

Corn (*Zea mays*) and soybean (*Glycine max*) production forms an integral part of economies worldwide, but yields are limited by biotic and abiotic factors associated with short rotations and long-term monocultures. In this study, a long-term rotation study site with corn and soybean planted in annual rotation, five-year rotation, and long-term monoculture was utilized to examine the relationships between crop sequences, soil fungal communities, soybean cyst nematode (SCN, *Heterodera glycines*) densities, soil properties, and crop yields. High throughput sequencing of the ITS1 region of fungal rDNA revealed that soil fungal community structure varied significantly by crop sequence, with fungal communities under five consecutive years of monoculture becoming progressively similar to communities in long-term monoculture plots associated with their respective crop hosts. Total alpha diversity was greater under corn, but patterns of diversity and relative abundance of specific functional groups differed by crop host, with more pathotrophs proliferating under soybean and more saprotrophs and symbiotrophs proliferating under corn. Soil phosphorus (P) varied significantly by crop sequence, with lower levels of P corresponding with relative abundance of Glomerales, Paraglomerales, and Sebacinales and higher levels of P corresponding with relative abundance of Mortierellales. Soil density of the SCN was positively correlated with relative abundance and diversity of nematode-trapping fungi and with relative abundance of many potential nematode egg parasites. These results suggest several possible explanations for the improved yields associated with crop rotation, including decreased pathogen pressure, modification of soil properties, and increased diversity of soil fungal communities. Future research should investigate the potential of nematode-trapping fungi to regulate SCN densities and examine the relationships between soil P and specific arbuscular mycorrhizal and mortierellalean fungi associated with corn and soybean hosts.

## 1. INTRODUCTION

Corn (*Zea mays*) and soybean (*Glycine max*) production forms an integral part of the economies of multiple nations in both the eastern and western hemisphere (Hartman et al., 2011; Meade et al., 2016) and plays a major role in the food security of developing nations (Hartman et al., 2011; Shiferaw et al., 2011). In the United States in 2016, corn and soybean accounted for 53% of total acreage planted to principal crops and 49% of principal crop production (U.S. Department of Agriculture National Agricultural Statistics Service, 2017). However, both crops suffer yield penalties under continuous monoculture. Corn yield is thought to be limited primarily by levels of soil nitrogen (N), whereas the main factor thought to limit soybean yield is disease pressure from plant pathogens, especially the soybean cyst nematode (SCN, *Heterodera glycines*) (Gentry et al., 2013; Seifert et al., 2017). Other soil properties, especially soil phosphorus (P), may be involved in yield declines of both crops (Bender et al., 2015; Xin et al., 2017). Soil fungi, including arbuscular mycorrhizal fungi (AMF), interact with plants, plant pathogens, and soil properties in complex ways that may affect crop yields (Francl and Dropkin, 1985; Johnson et al., 1992; Tylka et al., 1991).

The SCN is responsible for the largest soybean yield declines, averaging 3.5 million metric tons in yield losses in the U.S., annually (Koenning and Wrather, 2010). While crop rotation to corn is the most widely adopted practice to reduce SCN populations, even five years of continuous corn cropping cannot eliminate this pathogen (Porter et al., 2001). With few available sources of genetic resistance to SCN (Liu et al., 2017) and the phasing out of chemical nematicides due to environmental and human toxicity (Warnock et al., 2017), biocontrol of SCN using nematophagous fungi is being investigated as an alternative to manage this pathogen (Chen and Dickson, 2012). Understanding how these fungi respond to common corn-soybean crop sequences may help identify fungal taxa that could serve as biocontrol agents.

Methods for characterizing soil fungal communities have undergone significant advances in recent years. Early studies used denaturing gradient gel electrophoresis (DGGE) to demonstrate seasonal shifts in bacterial and fungal communities in plant rhizospheres and to a lesser extent in bulk soils (Gomes et al., 2003; Smalla et al., 2001; Smit et al., 2001), but this technique is known to underestimate species diversity (Gomes et al., 2003). In the last decade, high throughput amplicon sequencing has facilitated more complete characterization of agricultural soil microbial communities than earlier culture-based and molecular approaches (Lindahl et al., 2013). Many of these studies have demonstrated the effects of continuous monoculture cropping on the relative abundance of plant pathogens (Bai et al., 2015; Liu et al., 2014b; Wu et al., 2016; Xiong et al., 2014), while others have described the dynamics of beneficial microbes, including AMF (Martínez-García et al., 2015; Senés-Guerrero and Schüßler, 2016) and antagonists of plant pathogens (Hamid et al., 2017; Xu et al., 2012), including those with hypothesized roles in SCN-suppressive soils (Hu et al., 2017). High throughput amplicon sequencing has also been used to examine the effects of tillage (Hu et al., 2017; Smith et al., 2016), organic farming practices (Hartmann et al., 2014), fertilizer use (Lin et al., 2012; Ramirez et al., 2010), crop rotation (Bainard et al., 2017), land-use, and soil properties (Lauber et al., 2008) on soil microbial communities.

In addition to their effects on soil microbial communities, cropping sequences are known to affect soil physical and chemical properties that could impact yield. For example, rotation of corn to soybean has been shown to increase bioavailable N, which improves corn yield (Peterson and Varvel, 1989). Likewise, rotation of soybean to corn has been shown to improve soil aggregate stability and water holding capacity and to increase soil levels of organic matter, P, K, Mg, and Ca, properties that are associated with increased soybean yield (Perez-brandan et al., 2014). However, another study found that soil P was higher under continuous soybean monoculture compared to continuous corn monoculture at two sites in Minnesota, including our study site (Johnson et al., 1991). AMF, which are canonically thought to enhance plant growth through the uptake and transport of soil P (Parniske, 2008), undergo community shifts related to corn-soybean crop sequences (Johnson et al., 1991). It has been hypothesized that the AMF communities associated with corn and soybean becomes less mutualistic under continuous monoculture of each host, potentially contributing to yield declines (Johnson et al., 1992).

The population density of nematophagous fungi may also be affected by cropping sequences. For example, the nematode-endoparasitic fungus, *Hirsutella rhossiliensis*, which is thought to have a density-dependent relationship with SCN (Jaffee et al., 1992), was more commonly isolated from SCN juveniles during soybean cropping years with higher SCN density (Chen and Reese, 1999). Likewise, the SCN egg parasites, *Metacordyceps chlamydosporia* and *Purpureocillium lilacinum*, were shown to increase in relative abundance in the soybean rhizosphere and in SCN cysts over continuous soybean monoculture in fields in northeastern China (Hamid et al., 2017; Song et al., 2016a). Nematode-trapping fungi *Dactylellina ellipsospora* (syn. *Monacrosporium ellipsosporum*) and *Arthrobotrys dactyloides* (Syn: *Drechslerella dactyloides*) have been shown to have a density-dependent relationship with the root knot nematode (*Meloidogyne javanica*) under greenhouse conditions (Jaffee et al., 1993). However, little is known about the relationships between common corn-soybean crop sequences and SCN-antagonistic fungi in bulk soil.

In addition to studying shifts in the abundance of fungal taxa, we also sought to address changes in functional guilds of fungi across crop rotation sequences. The concept of an ecological guild dates to Root (1967), who defined a guild as a group of species that consume the same resource in the same way. Recent approaches have broadly classified fungi according to both their trophic mode and ecological guild (Nguyen et al., 2016). In addition to these broad ecological categories, three guilds of nematophagous fungi are recognized: i) near-obligate endoparasites of free-living nematodes, ii) nematode egg parasites, and iii) nematode-trapping fungi (Chen and Dickson, 2012). A fourth group of nematode-antagonistic fungi consisting of species that kill nematodes through the production of antibiotics is often described (Chen and Dickson, 2012), but this group does not precisely meet the definition of a guild, as its description does not specify a feeding mode. To our knowledge, this is the first study to comprehensively investigate the bulk soil fungal community associated with corn and soybean rotations from the standpoint of trophic modes and nematophagous guilds in addition to individual taxa or operational taxonomic units (OTUs).

This study utilized a unique long-term research site in Waseca, MN, at which corn and soybean have been planted under continuous long-term monoculture, annual rotation, and 5-year rotations since 1982. A previous study conducted at the same site showed that populations of SCN increased under continuous soybean monoculture, whereas populations of the plant parasitic nematodes, *Pratylenchus* (lesion nematode) and *Helicotylenchus* (spiral nematode), increased under continuous corn monoculture (Grabau and Chen, 2016a, 2016b). These nematode populations were correlated with yield declines in soybean and corn, respectively. However, it is unknown whether increases in SCN density at this site are correlated with increases in the relative abundance of SCN-antagonistic fungi in the bulk soil, knowledge that could have profound implications for biocontrol. The objectives of this study were to (i) investigate the role of corn/soybean crop rotations and continuous monoculture in shaping bulk soil fungal communities, and (ii) identify specific fungal taxa or functional guilds correlated with corn and soybean yield, soil properties, and SCN density. We hypothesized that (i) bulk soil fungal communities under consecutive years of monoculture would become progressively more similar to those in long-term monoculture plots; (ii) crop rotation would result in greater alpha diversity of fungal communities compared to continuous monoculture; and (iii) nematophagous fungi would increase in diversity and relative abundance over continuous soybean monoculture, corresponding with an increase in density of the SCN.

## 2. MATERIALS AND METHODS

### 2.1 Experimental design

The experimental design used in this study is described in previous studies conducted at the same site (Grabau and Chen, 2016b, 2016a; Hu et al., 2018). Long-term rotation plots at the University of Minnesota Southern Research and Outreach Center in Waseca, MN (44°04’ N, 93°33’ W), where corn and soybean have been planted in continuous cropping sequences since 1982, were used to study the effects of crop sequences on the bulk soil fungal community in 2015 and 2016. The crop sequences in this study were continuous corn and soybean monocultures (Cc and Ss), 5-year rotations (C1-C5 and S1-S5), annual rotations (Ca and Sa), and non-*Bt* corn and SCN-resistant soybean monocultures (Cn and Sr) (**Table S1**). Since 2010, all treatment plots except for Cn and Sr plots were planted with *Bt* corn (Dekalb 50-66) and SCN-susceptible soybean (Pioneer 92Y12). The Cn and Sr plots were planted with non-*Bt* corn (Dekalb 50-67) and SCN-resistant soybean (Pioneer 92Y22), respectively, since 2010, but prior to 2010 all soybean plots were planted with SCN-susceptible soybeans (**Table S1**). There were four replicates for each treatment, arranged in the field in a randomized complete block design.

Experimental plots were 4.57 m wide by 7.62 m long and contained 6 rows of plants, each. Plots were managed by conventional tillage practices consisting of fall chisel plowing and field cultivation prior to planting. All crops were resistant to glyphosate (Roundup) herbicide, which was used at a rate of 2.2 kg/ha to prevent weed growth. Corn and soybean plots were sprayed with Endigo insecticide at a rate of 245 g/ha at midseason in 2015 for aphid control. No insecticides were used in 2016. Corn plots were fertilized with nitrogen as urea at a rate of 224.4 kg/ha at midseason in 2015 and 2016. No phosphorus-containing fertilizer was applied in 2015 and 2016, but P-K fertilizer was applied at a rate of 80 and 120 lbs. per acre, respectively, to all plots in 2014.

### 2.2 Sample collection

Bulk soil samples were collected in the spring (at planting), midseason (2-3 months after planting), and fall (at harvest). A total of 20 soil cores were taken at regular intervals from between the two central rows of each plot using a 2.54 cm diameter probe sunk to a depth of 20 cm. Soil samples were stored in a cold room at 4 °C until further processing on the same day. Soil cores from each plot were pooled, pushed by hand through a 5 mm mesh screen to break apart cores into smaller aggregates, and thoroughly mixed by hand. A subsample of 100 g of homogenized soil was used for SCN egg quantification. Soil properties analysis was performed on 100 g of homogenized soil collected in the spring by the Research Analytic Laboratory at the University of Minnesota. The remaining homogenized soil (250 g) was stored in a resealable plastic bag at −80 °C for later processing.

### 2.3 SCN egg density quantification and yield measurement

The density of SCN eggs in the bulk soil was determined following the methods described in Hu, et al. (2017). Briefly, cysts were separated from 100 cm^3^ of bulk soil by elutriation followed by centrifugation in a 63% (w/v) sucrose solution. Cysts were crushed to release eggs (Niblack et al., 1993). Eggs were then collected in water and quantified by examining a subsample of egg suspension with an inverted microscope. This number was used to calculate the total number of eggs in 100 cm^3^ of soil. Yield was measured from 20 feet of the two central rows of each experimental plot using a plot combine.

### 2.4 DNA extraction, amplification, and sequencing

Fifty grams of soil from each 250 g soil sample was homogenized further in a coffee grinder, and DNA was extracted from 0.25 g of this soil using a MoBio PowerSoil^®^ DNA Isolation Kit. Twenty microliters of DNA per sample were submitted to the University of Minnesota Genomics Center (Saint Paul, MN, USA) for amplification and sequencing. Additional samples to assess the quality and accuracy of the amplification and sequencing were also included. These included a fungal mock community, a synthetic mock community (Palmer et al., 2018), technical replicates consisting of replicate DNA extractions from the same soil sample, and negative controls in which no soil was added to the extraction kit. For a description of the preparation and analysis of mock communities, see the **Supplementary text**.

A two-step dual-indexed amplification method was used for amplifying fungal internal transcribed spacer 1 (ITS1) regions of fungal rDNA (Gohl et al., 2016). Paired-end sequencing (2 × 250 bp) was carried out using a MiSeq 600 cycle v3 kit on a total of four lanes. 2015 fall and midseason bulk soil samples were sequenced on one lane, and spring 2015 bulk soil samples were sequenced on a separate lane along with technical replicates and a pool of 51 other samples. All 2016 bulk soil PCR products were pooled and sequenced on two lanes, with half of the second lane shared with a pool of 229 other samples. Fungal mock community samples were included in all four lanes. Illumina sequencing on these four lanes resulted in a total 47,902,914 reads from bulk soil samples that passed Illumina quality control. A total of 7 bulk soil samples, which had fewer than 20,000 reads after quality control, were re-submitted for amplification and sequencing, resulting in an additional 744,408 quality reads.

### 2.5 Bioinformatics Processing

Fastq files of reads that passed Illumina quality control were stored and processed at the Minnesota Supercomputing Institute (MSI, Minneapolis, MN, USA). The 2016 bulk soil fastq files from identical samples in separate lanes were merged using the fastq-cat.pl script in the Gopher-biotools package (Garbe, 2015). Preprocessing and merging of reads, OTU clustering, filtering, and curating were performed using the AMPtk v1.1.0 pipeline with default parameters, except where noted (Palmer et al., 2018). The trim length for illumina reads was set to 250 bp for pre-processing. Processed and merged reads were clustered in AMPtk using the “dada2” option, which first clusters sequences with DADA2 v1.6 (Callahan et al., 2016) and then clusters the output iSeqs of DADA2 into OTUs using USearch v9.2.64 (Edgar, 2013). The OTU table was filtered using AMPtk’s default index bleed filtering rate of 0.05% and curated using LULU v1.1.0, which merges “split” OTUs, resulting in better alpha diversity estimates (Frøslev et al., 2017).

Based on the recovery of taxa from our fungal mock community, a synthesis of four different methods and a custom R script (available at https://github.com/stro0070/OTU-taxonomy-assignment) was used for taxonomy assignment (**Supplementary text**). This new approach to assigning taxonomy to OTUs resulted in a greater percentage of recovered taxa from our fungal mock community, as well as greater accuracy and precision of taxonomy assignments compared to any single method. The OUT table with taxonomy was filtered to include only fungal OTUs and members of the synthetic mock community. The AMPtk pipeline resulted in 12,068 OTUs, of which 11,701 remained after filtering to remove non-fungal OTUs and adding back synthetic mock community OTUs. The final OTU table was filtered to remove OTUs with fewer than 10 reads across the entire table, after which 8,587 OTUs remained, of which 8,491 were present in bulk soil samples. After filtering, sequence depth for bulk soil samples ranged from 16,435 to 205,157 reads with an average sequence depth of 63,947. The rarefaction curves for most samples began to level off by around 30,000 reads, indicating the sampling depth was adequate to capture most of the fungal diversity (**Figure S1**).

Fungal taxonomy is currently undergoing substantial revision. As of the date of this writing, some phylum names in the UNITE database are outdated, and subphyla names are not included (UNITE Community, 2017). We decided to retain the UNITE database taxonomy assignments without modification due to the complex nature of the effort that would be required to update them. Thus, we report, for example, on the abundance of Glomeromycota rather than Glomeromycotina and on Mortierellomycota instead of Mortierellomycotina (Spatafora et al., 2016). Results of taxonomy assignments, fungal and synthetic mock communities, and technical replicates analysis are described in the **Supplementary text, Tables S2-5, and Figures S2 and S3**.

FUNGuild v1.1.0 (Nguyen et al., 2016) was used to assign ecological guilds and trophic modes to OTUs. Only OTUs that were assigned a trophic mode with a confidence ranking of “Probable” or higher were used for statistical analyses of trophic modes. The nematophagous fungal guilds, “endoparasites,” “egg parasites,” and “trapping fungi,” were defined by performing literature searches for SCN-parasitic fungi (keywords: “soybean cyst nematode”, “biological control,” “fungi,” “nematophagous”, “trapping”, “predator,” “endoparasite”, “egg parasite”) (Hu et al., 2018). Taxa belonging to each nematophagous guild are listed in **Table S6**.

### 2.6 Statistical analyses

All statistical analyses were performed in RStudio v 3.5.0 (R Core Team, 2018; RStudio Team, 2016). All plots except for distance-based redundancy analysis were created in “ggplot2” (Wickham, 2009). In all analyses in which ANOVA was used, Levene’s test (Levene, 1960) was used to ascertain equal variance, and all *P*-values associated with multiple hypothesis testing were corrected by the false discovery rate (FDR) procedure (Benjamini and Hochberg, 1995).

#### 2.6.1 Yield and SCN density

Significant differences in yield were detected using ANOVA followed by Tukey’s HSD test (Tukey, 1949). Due to unequal variance that could not be corrected through transformation, the Kruskal-Wallis test (Kruskal and Wallis, 1952) was used to compare SCN density by crop sequence and by season for individual crop hosts. Post-hoc comparisons were performed using the “kruskal” function (Mendiburu, 2017). Correlations between cube root-transformed SCN density and susceptible soybean yield were tested by fitting linear models (Yield ~ SCN Density) with the “lm” function in R (R Core Team, 2018) followed by ANOVA.

#### 2.6.2 Soil properties

The mean and standard error for soil properties associated with each crop sequence were calculated using the “aggregate” function in the “stats” package in R (R Core Team, 2018). Significant effects of crop sequence on individual soil properties were tested using ANOVA on the model Y ~ Block + CropSequence. A square root transformation was used for Fe in 2015 and 2016 and for NO_3_ in 2016 to meet the ANOVA assumption of equal variance. Individual soil properties were tested for correlations with fungal community dissimilarity in spring 2015 and 2016 using the “adonis” function with 9999 permutations in the R package, “vegan” (Oksanen et al., 2016). Vectors for each soil property were generated using the “envit” function. Correlations between individual soil properties and yield were tested using the “lm” function followed by ANOVA.

#### 2.6.3 Differential abundance

Differential abundance analyses were performed at the phylum and species level, for trophic modes, and for nematophagous guilds. Due to the high false discovery rate associated with using proportions for differential abundance testing (Weiss et al., 2017), read counts were instead transformed to centered log ratios (CLRs) (Aitchison, 1982; Martín-Fernández et al., 2003), which are associated with a much lower false discovery rate (Gloor et al., 2017). This is the same approach used in the R package “ALDEx2,” (Gloor, 2018) and has been used in other metabarcoding studies (Lee et al., 2014; Mcdonald et al., 2018). CLRs of taxa, trophic modes, and nematophagous guilds with equal variance according to Levene’s test were compared across crop sequences using ANOVA on the linear model, Y ~ Block + CropSequence for each sampling timepoint (e.g. Spring 2015) and across seasons using ANOVA on the linear model Y ~ Block + CropSequence + Season + CropSequence:Season for each sampling year. For CLRs that failed Levene’s test, the Kruskal-Wallis test (Kruskal and Wallis, 1952) was used, instead. Post-hoc comparisons were made using Tukey’s HSD test (Tukey, 1949) or the “kruskal” function with FDR correction in the R package, “agricolae” (Mendiburu, 2017).

Spearman correlation tests were used to test for correlations between CLRs of taxa, trophic modes, and nematophagous guilds and soybean and corn monoculture year, SCN density, and yield in each SeasonYear (e.g. Fall 2016). Data from Sr and Cn plots were removed prior to testing for correlations between taxa/guilds and yield, because these cultivars were a different genotype than plants in the majority of treatments. In addition to removing data from Sr and Cn plots, data from Sa and Ca plots were removed prior to testing for correlations between taxa and monoculture year of either host, because annual rotation plots could not be assigned a numeric monoculture year.

#### 2.6.4 Fungal community diversity

Beta diversity analyses were performed on the OTU table subset by each SeasonYear (e.g. Fall 2016). Read counts were converted to relative abundances and square root transformed (Hellinger transformation) (Legendre and Gallagher, 2001), and a Bray-Curtis dissimilarity matrix was constructed using these transformed data (Bray and Curtis, 1957). Non metric multidimensional scaling (NMDS) (Kruskal, 1964) was performed using the “metaMDS” function in vegan (Oksanen et al., 2016). The *R*^*2*^ and *P*-values for the effect of crop sequence on fungal communities were calculated using the “adonis” function in vegan with 9999 permutations on the formula, Y ~ Crop Sequence. Additionally, the adonis function was used to parse the effects of block, season, and crop sequence on fungal community dissimilarity data subset by year, using the formula Y ~ Block + Season + CropSequence + Season:CropSequence. Distance-based redundancy analysis was performed using the “capscale” function in vegan (Legendre and Anderson, 1999; Oksanen et al., 2016) using an OTU table collapsed by order with the QIIME2 “collapse” plugin (Caporaso et al., 2010). Read counts were Hellinger transformed prior to this analysis. The analysis was constrained by soil properties, spring SCN density, and yield and scaled by “species”.

Alpha diversity metrics were calculated using the “estimate_richness” function in the “Phyloseq” package (McMurdie and Holmes, 2013). Effects of crop sequence and season on alpha diversity were tested for individual years using ANOVA on the same model as for beta diversity. Pairwise comparisons across crop sequences and seasons were made using Tukey’s HSD test (Tukey, 1949). Spearman correlation tests were used to test for correlations between the diversity of nematophagous guilds and trophic modes and SCN density, yield, and monoculture year of each crop host. Rarefaction curves were generated using the “rarecurve” function in vegan (Oksanen et al., 2016).

## 3. DATA ACCESSION

The raw sequences from bulk soil samples were deposited into the NCBI database (Accession number: PRJNA484933).

## 4. RESULTS

### 4.1 Soybean cyst nematode density and crop yields

Corn yields were significantly higher in years following soybean (C1 and Ca), especially in the first year of corn after five years of soybean (C1) (Figure 1A). However, long-term corn monoculture (Cc) yield was not significantly lower than corn yield from earlier years of corn monoculture (C1-C5) (Figure 1A). By contrast, soybean did not have significantly higher yields in the first year following five years of corn (S1) or in annual rotation with corn (Sa) compared to other soybean crop sequences (Figure 1B). In 2015, SCN-resistant soybean (Sr) had significantly greater yields than SCN-susceptible soybean (Ss), which had significantly lower yields than SCN-susceptible soybean under shorter-term monocultures (S3 and S5) (Figure 1B).

**Figure 1.**
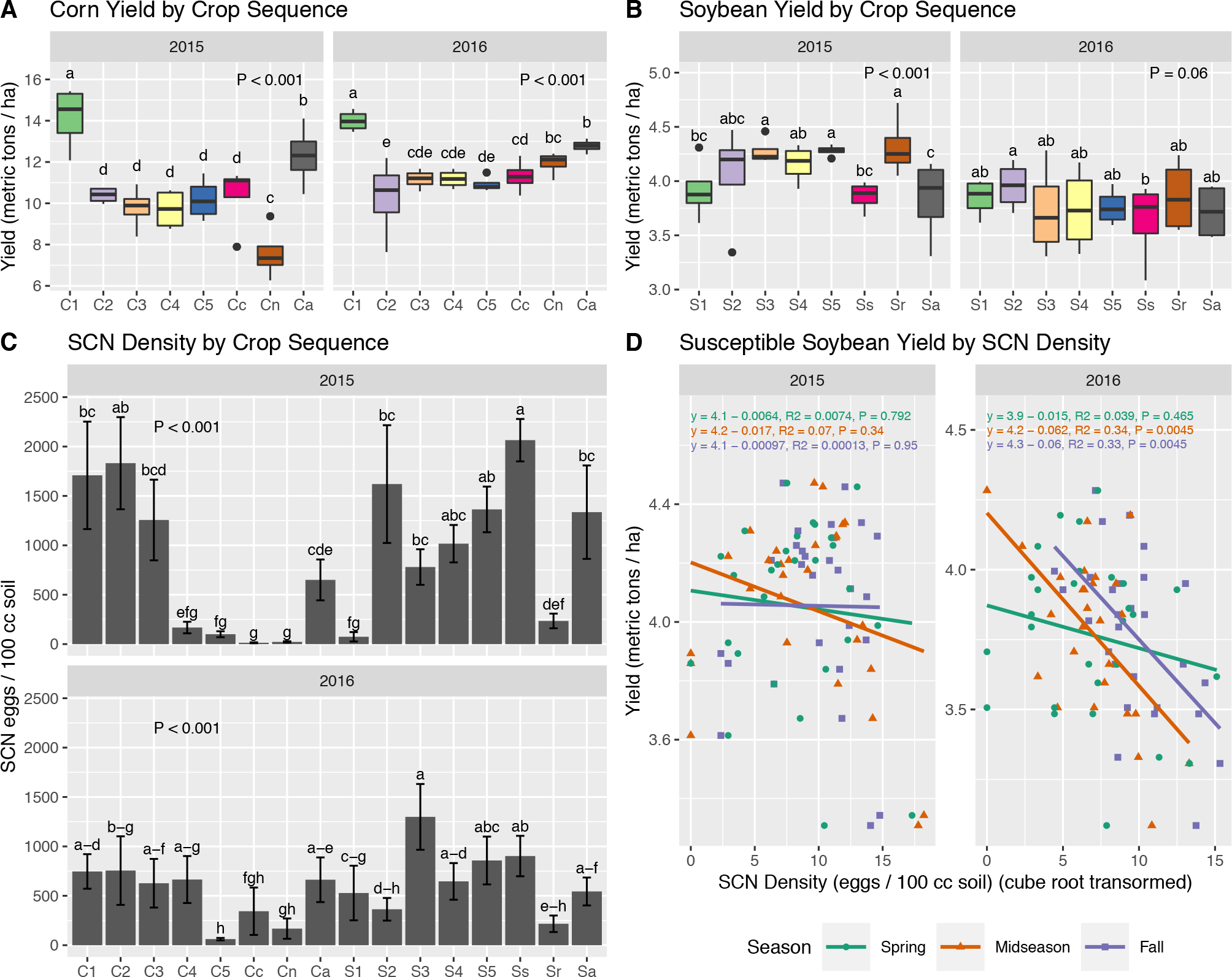
Corn (**A**) and soybean (**B**) yields were compared using ANOVA on the linear model, Yield ~ Block + Crop Sequence. Reported *P*-values are associated with the Crop Sequence term in this model. Different lowercase letters denote significant differences in mean yield as determined by Tukey’s honestly significant difference (HSD) test. SCN density (**C**) was compared across crop sequences using the Kruskal-Wallis test on data pooled from all seasons (spring, midseason, and fall) within a sampling year, and significance categories were calculated using the “kruskal” function in R at a significance level of α= 0.05 with false discovery rate (FDR)-correction. Error bars display one standard error above and below the mean. (**D**) Relationships between soybean yield and SCN density are described by linear models. *P*-values associated with these models are FDR-corrected. Crop sequences include 5 consecutive years of corn after the 5th year of soybean (C1-C5); 5 consecutive years of soybean after the 5th year of corn (S1-S5); soybean in annual rotation with corn (Sa); corn in annual rotation with soybean (Ca); long-term *Bt* corn (Cc) and SCN-susceptible soybean (Ss) monocultures; non-*Bt* Corn (Cn) and SCN-resistant soybean (Sr) monocultures.

SCN density differed significantly by crop sequence in 2015 and 2016 according to the Kruskal-Wallis test (*P* < 0.0001) (Figure 1C). In general, SCN egg density decreased with increasing years of corn monoculture and increased with increasing years of soybean monoculture (Figure 1C). SCN density was also significantly lower in SCN-resistant soybean plots compared to other soybean plots (Figure 1C). Significant (*P* < 0.01) negative correlations between 2016 soybean yield and 2016 midseason and fall SCN density were observed (Figure 1D). When SCN density data were subset by crop host, a significant effect of season was observed for SCN-susceptible soybean in 2016 (*P* < 0.0001), with higher SCN density in fall compared to spring or midseason (**Figure S4**).

### 4.2 Fungal community comparisons across crop sequences and seasons

Adonis, a non-parametric multivariate ANOVA used to identify sources of variation between communities (Anderson, 2001; Oksanen, 2015), revealed significant effects of crop sequence and season on fungal community dissimilarity, with crop sequence explaining the highest proportion of dissimilarity in both 2015 and 2016 (Table 1). NMDS plots from each sampling timepoint displayed a pattern or trend in which long-term continuous monoculture communities showed the clearest separation by crop host, with long-term soybean monoculture (Ss and Sr) communities on opposite sides of the NMDS plots from long-term corn monoculture (Cr and Cn) communities (Figure 2A).

**Figure 2.**
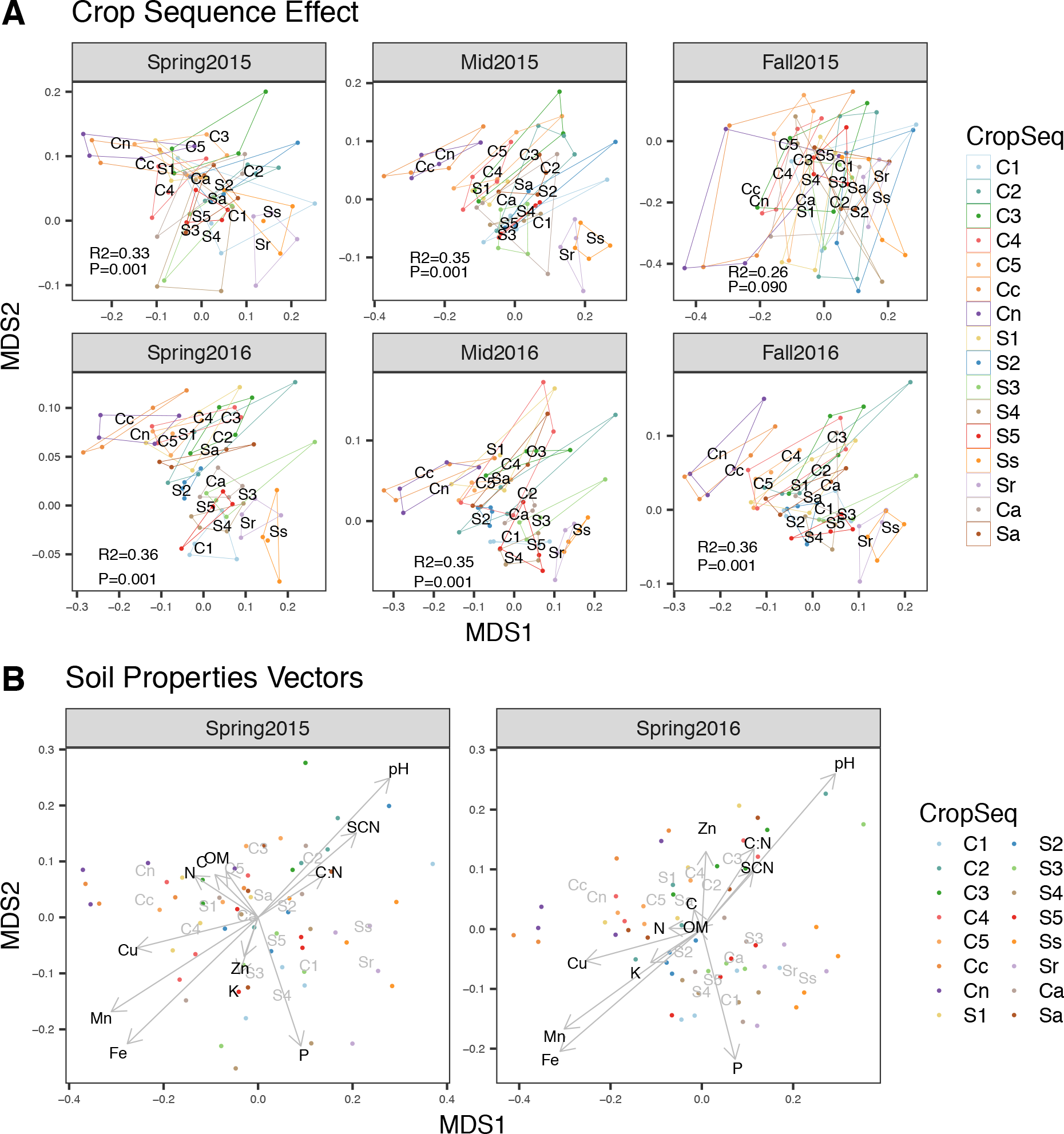
(**A**) Non-metric multidimensional scaling (NMDS) plots showing the effect of crop sequence on fungal communities in each SeasonYear. *R*^*2*^ and *P*-values were derived from adonis on the formula, Bray-Curtis Distance ~ Crop Sequence. (**B**) NMDS plots of fungal communities with environmental data vectors. Vectors indicate the weight and direction of soil properties and soybean cyst nematode (SCN) density in relation to fungal communities. Crop sequences include 5 consecutive years of corn after the 5th year of soybean (C1-C5); 5 consecutive years of soybean after the 5th year of corn (S1-S5); soybean in annual rotation with corn (Sa); corn in annual rotation with soybean (Ca); long-term *Bt* corn (Cc) and SCN-susceptible soybean (Ss) monocultures; non-*Bt* Corn (Cn) and SCN-resistant soybean (Sr) monocultures.

Between these extremes, communities generally became more similar to the long-term monoculture communities with increasing years of monoculture of each crop. For example, communities from later years of corn or soybean monoculture were more similar to their respective long-term monoculture communities, with communities from S3, S4, and S5 mapping close to Ss and Sr and communities from C3, C4, and C5 mapping close to those from Cc and Cn. Communities associated with first year crop sequences (S1 and C1), however, were more similar to fifth year crop sequence communities associated with the alternate crop (C5 and S5), especially in spring. These communities from first year crop sequences became increasingly more similar to their respective host communities during the growing season from spring to fall (Figure 2A).

**Table 1.** Adonis results. *R*^*2*^ and *P*-values were calculated based on the formula, Community Distance ~ Block + Season + Crop Sequence + Season:Crop Sequence. Bray-Curtis distance was calculated using Hellinger-transformed OTU counts. Significant effects are reported at \*\**P* < 0.01.

|  | 2015 |  | 2016 |  |
| --- | --- | --- | --- | --- |
| | $R^2$ | $P$-value | $R^2$ | $P$-value |
| Season | 0.11 | 0.001** | 0.057 | 0.001** |
| CropSeq | 0.17 | 0.001** | 0.23 | 0.001** |
| Season:CropSeq | 0.11 | 0.998 | 0.10 | 1 |

### 4.3 Soil properties

Of the soil properties tested, only P, Mn, and Cu were found to differ significantly by crop sequence (**Table S7**). Phosphorus decreased under continuous corn monoculture and increased under continuous soybean monoculture in both 2015 and 2016, while Mn and Cu showed the reciprocal pattern. Although all measured soil properties except for Zn had significant relationships with fungal community dissimilarity in at least one sampling year (**Table S7 and** Figure 2B), only pH, Fe, and Mn had *R*^*2*^ values greater than 0.1 according to univariate adonis (**Table S7**). The vector for P on the NMDS plot showed an association with soybean crop sequences Figure 2B), and P levels under continuous soybean (Ss and Sr) were over twice those observed under continuous corn (Cc) (**Table S7**). Correlations between individual soil properties and yields of corn and soybean typically had opposite signs for the two crops (**Table S8**). While significant relationships between soil properties and corn yield were inconsistent between 2015 and 2016, soybean yields were consistently significantly positively correlated with organic matter (OM), manganese (Mn), copper (Cu), and total organic carbon (TOC), and significantly negatively correlated with pH (**Table S8**).

### 4.4 Alpha diversity

Chao1 estimates of fungal OTUs varied significantly by crop sequence in 2015 and by season in both years (Table 2). Although there were no pairwise significant differences in Chao1 between crop sequences according to Tukey’s HSD test, Chao1 was significantly greater under corn monoculture when compared between crop hosts (**Table S9**) and tended to increase under continuous corn monoculture and decrease under continuous soybean monoculture (Figure 3A, **Table S10**). Numbers of observed OTUs assigned to two different trophic modes by FUNGuild (Nguyen et al., 2016) significantly differed by crop sequence and season (Table 2) and displayed consistent patterns in relation to crop host, with saprotrophs and symbiotrophs having greater diversity under corn compared to soybean, especially under long-term monoculture (Figure 3B, **S9**). Symbiotroph diversity also showed significant negative correlations with SCN density and corn yield at one sampling timepoint, each (**Table S10**). Richness of OTUs assigned to nematode-trapping fungi and egg parasite guilds varied significantly by crop sequence in both years and by season in 2015 (Table 2). Fifth-year and long-term soybean monoculture plots (S5 and Ss) and first-year corn monoculture plots (C1) typically had the most trapping fungi and egg parasite OTUs, whereas long-term corn monoculture plots (Cn and Cc) typically had the fewest (Figure 3C). The diversity of nematode-trapping and egg parasitic guilds was significantly correlated with SCN egg density in spring of both years and negatively correlated with corn monoculture year in multiple timepoints, each (**Table S10**).

**Table 2.** Alpha diversity metrics across seasons and crop sequences. Chao1 is total alpha diversity of the bulk soil fungal communities. Diversity metrics are also reported for individual nematophagous guilds and for three trophic modes. Only OTUs assigned a trophic mode with a confidence ranking of “Probable” or “Highly Probable” were used for this analysis. Significant *P*-values associated with ANOVA on the model Diversity ~ Block + Season + Crop Sequence + Season:Crop Sequence are reported at \**P* < 0.05, \*\**P* < 0.01, and \*\*\**P* < 0.001. Different lowercase letters indicate significant differences in means according to Tukey’s honestly significant difference (HSD) test.

| 2015 |  |  |  |  |  |  |  |
| --- | --- | --- | --- | --- | --- | --- | --- |
| ANOVA | Chao1 | Saprotrophs | Pathotrophs | Symbiotrophs | Trapping fungi | Egg parasites | Endoparasites |
| Season | <0.001*** | <0.001*** | <0.001*** | <0.001*** | <0.001*** | 0.02* | 0.67 |
| CropSeq | 0.024* | 0.0010** | 0.38 | <0.001*** | 0.0059** | <0.001*** | 0.56 |
| Season:CropSeq | 0.95 | 0.67 | 0.87 | 0.16 | 0.0051** | 0.94 | 0.70 |
| Season |  |  |  |  |  |  |  |
| Spring | 824a | 98a | 27a | 35b | 3.95a | 5.38b | 0.55a |
| Mid | 842a | 94a | 24a | 62a | 3.58a | 5.69ab | 0.5a |
| Fall | 618b | 72b | 21b | 36b | 2.39b | 6.09a | 0.44a |
| 2016 |  |  |  |  |  |  |  |
| ANOVA | Chao1 | Saprotrophs | Pathotrophs | Symbiotrophs | Trapping fungi | Egg parasites | Endoparasites |
| Season | <0.001*** | <0.001*** | <0.001*** | <0.001*** | 0.058 | 0.31 | 0.24 |
| CropSeq | 0.12 | 0.044* | 0.19 | <0.001*** | <0.001*** | <0.001*** | 0.023* |
| Season:CropSeq | 0.98 | 0.98 | 0.66 | 0.85 | 0.99 | 0.98 | 0.34 |
| Season |  |  |  |  |  |  |  |
| Spring | 969b | 103b | 26b | 53b | 3.69a | 5.53a | 0.52a |
| Mid | 1018b | 106b | 28b | 67a | 3.55a | 5.69a | 0.36a |
| Fall | 1161a | 117a | 35a | 68a | 4.28a | 5.86a | 0.5a |

**Figure 3.**
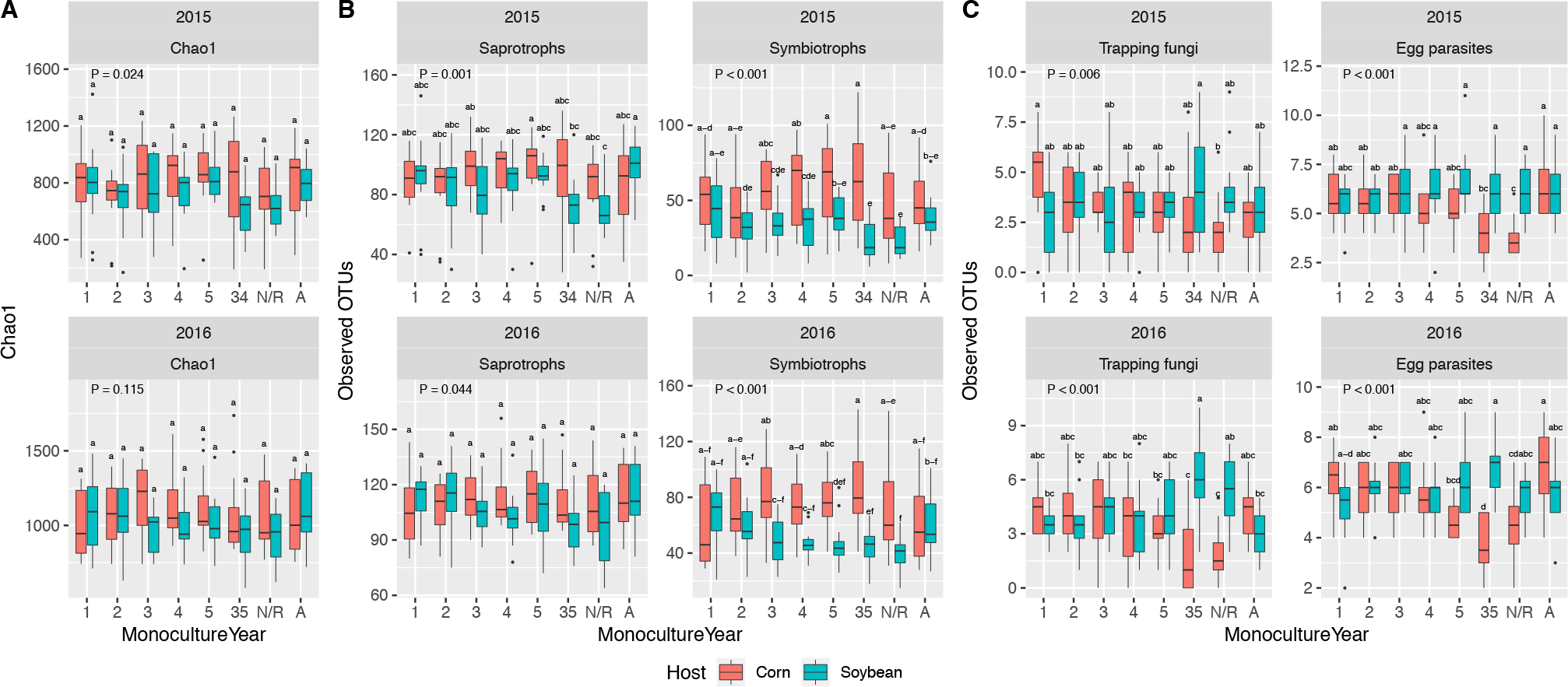
Alpha diversity of fungal community, trophic modes, and nematophagous guilds by crop host and monoculture year. (**A**) Total fungal community alpha diversity, estimated by Chao1. (**B**) Observed OTUs classified as saprotrophs and symbiotrophs according to FUNGuild. (**C**) Observed OTUs classified as nematode-trapping fungi or nematode egg parasites. The x-axis denotes monoculture year of corn and soybean: **1-5** = years of 1-5 of continuous soybean and corn monoculture; **34,35** = long-term corn and soybean monocultures of 34 and 35 years; **N/R** = continuous non-*Bt* Corn and SCN-resistant soybean monocultures since 2010. **A** = corn-soybean annual rotation plots. Data from spring, midseason, and fall within each year were pooled prior to analysis. *P*-values are associated with the Crop Sequence term according to ANOVA on the linear model, Diversity ~ Block + Season + Crop Sequence + Season:Crop Sequence. Different letters denote significant differences in means according to Tukey’s honestly significant difference (HSD) test.

### 4.5 Fungal community profiles

Averaging across all sampling timepoints and crop sequences, a majority of reads belonged to OTUs assigned to phylum Ascomycota (56%), with Basidiomycota (14%), Mortierellomycota (13%), and Chytridomycota (2%) having the next most abundant read counts (Figure 4A). A significant portion of reads (12%) were assigned to kingdom Fungi but not to any fungal phylum. Fewer than 1% of reads, each, were assigned to Blastocladiomycota, Calcarisporiellomycota, Entomophthoromycota, Entorrhizomycota, Glomeromycota, Kickxellomycota, Monoblepharomycota, Mucoromycota, Neocallimastigomycota, Olpidiomycota, Rozellomycota, and Zoopagomycota.

**Figure 4.**
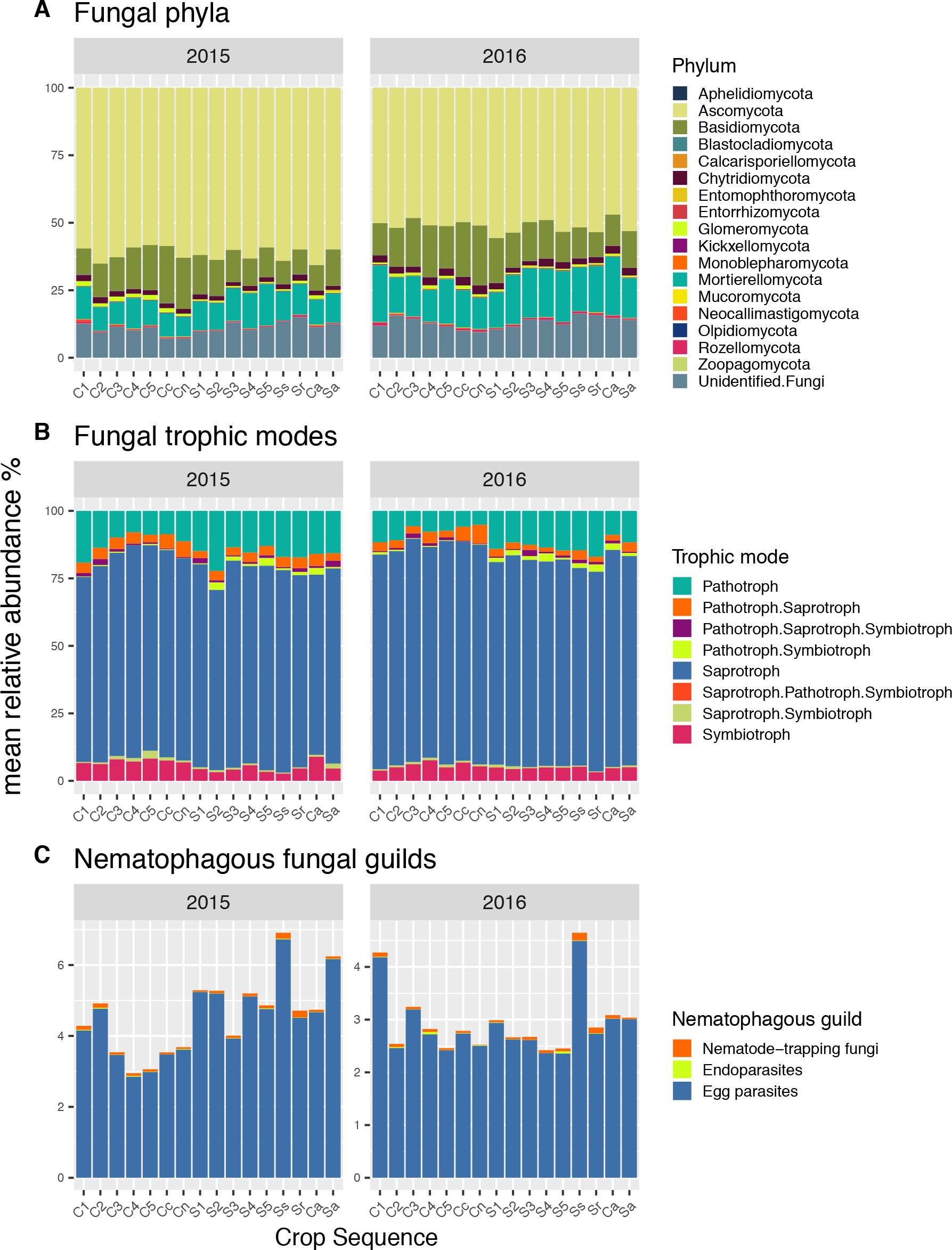
Fungal community profiles by crop sequence for each year. Data from each season (spring, midseason, and fall) were pooled prior to plotting. Each bar represents the average proportion of (**A**) phyla, (**B**) trophic modes, or (**C**) nematophagous guilds associated with each crop sequence in a given year. Only OTUs that were assigned a trophic mode with a confidence ranking of “Probable” or higher were included in the trophic mode profile plots. Crop sequences include 5 consecutive years of corn after the 5th year of soybean (C1-C5); 5 consecutive years of soybean after the 5th year of corn (S1-S5); soybean in annual rotation with corn (Sa); corn in annual rotation with soybean (Ca); long-term *Bt* corn (Cc) and SCN-susceptible soybean (Ss) monocultures; non-*Bt* Corn (Cn) and SCN-resistant soybean (Sr) monocultures.

Seventy-nine percent of sequence reads were clustered into OTUs that could not be assigned a functional guild or trophic mode at a confidence ranking of “Probable” or higher by FUNGuild (Nguyen et al., 2016). Of sequence reads belonging to OTUs that were assigned a “Probable” or “Highly Probable” trophic mode, 76% were classified as Saprotrophs, 12% as Pathotrophs, 5% as Symbiotrophs, 3% as Pathotroph-Saprotrophs, and 1% or less each as Pathotroph-Symbiotrophs, Saprotroph-Symbiotrophs, Pathotroph-Saprotroph-Symbiotrophs, and Saprotroph-Pathotroph-Symbiotrophs. (Figure 4B). A list of taxa assigned to each trophic mode is given in **Table S11**. A majority of reads belonging to OTUs assigned to ecological guilds with “Probable” or “Highly Probable” assignments were classified as Undefined Saprotrophs (66%), followed by Plant Pathogens (10%), Undefined Saprotroph Wood Saprotrophs (3%), and Arbuscular Mycorrhizal fungi (3%), with each other category making up 2% or less of the total. (**Figure S5**). A list of taxa assigned to each guild is given in (**Table S12**).

In terms of nematophagous fungal guilds, 3.7% of total reads belonged to OTUs classified as nematode egg parasites, while a much smaller proportion belonged to OTUs classified as nematode-trapping fungi (0.07%) and nematode endoparasites (0.01%) (Figure 4C). Four species belonging to the nematode endoparasite guild, nine species belonging to the nematode egg parasite guild, and six genera containing twenty-one species of nematode-trapping fungi were detected in our sequencing data (Figure 5 **and Table S13**). Additional BLAST searches of OTUs identified as “*Clonostachys*” and “*Purpureocillium*” against the UNITE database (UNITE Community, 2017) revealed that known egg-parasitic taxa, *Clonostachys rosea* and *P. lilacinum*, respectively, were their most likely identities, confirming that these OTUs belonged in the egg parasite guild.

**Figure 5.**
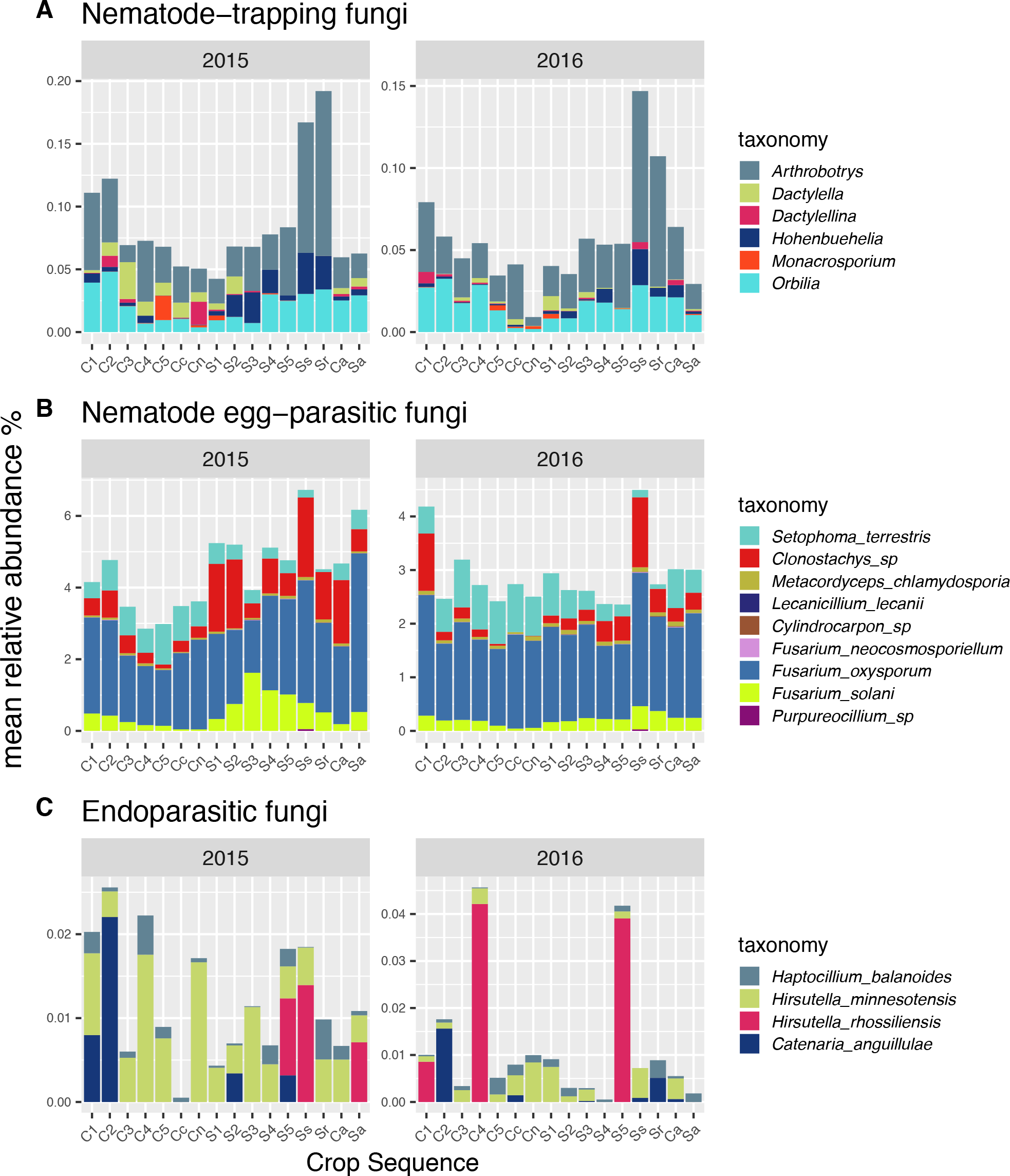
Nematophagous fungal guild profiles by crop sequence in each year. Data from each season (spring, midseason, and fall) were pooled prior to creating profile plots. Each bar represents the average proportion of (**A**) nematode-trapping fungal taxa, (**B**) nematode egg-parasitic taxa, or (**C**) nematode endoparasitic taxa associated with each crop sequence in a given year. Crop sequences include 5 consecutive years of corn after the 5th year of soybean (C1-C5); 5 consecutive years of soybean after the 5th year of corn (S1-S5); soybean in annual rotation with corn (Sa); corn in annual rotation with soybean (Ca); long-term *Bt* corn (Cc) and SCN-susceptible soybean (Ss) monocultures; non-*Bt* Corn (Cn) and SCN-resistant soybean (Sr) monocultures.

### 4.6 Differential abundance by crop sequence

While it is impossible to ascertain true shifts in absolute abundance of taxa through metabarcoding, CLRs (centered log ratios) can be used to identify shifts in relative abundance with an acceptably low false discovery rate (Gloor et al., 2017). Using the CLR transformation, two main patterns were observed regarding crop sequence effects on the relative abundance of fungal taxa, trophic modes, and nematophagous guilds: 1) an increase in relative abundance over years of soybean monoculture and a decline over years of corn (soybean-associated fungi), and 2) an increase in relative abundance over years of corn monoculture and a decline over years of soybean (corn-associated fungi) (Figures 6, 7, **S6-8, Table S14**). CLRs associated with annual rotation plots were often intermediate to those from long-term monoculture plots associated with either crop host.

**Figure 6.**
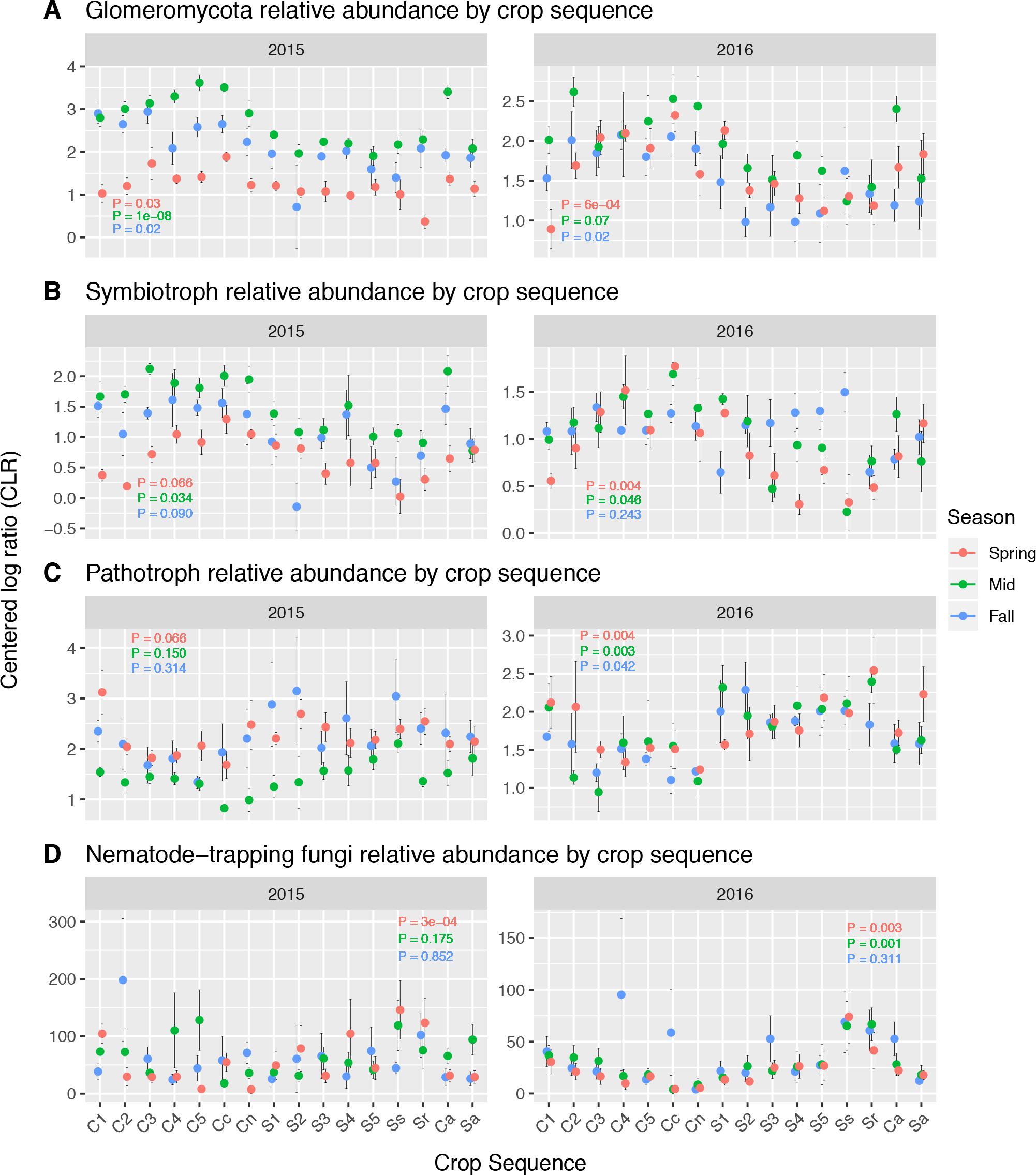
Differential abundance of (**A**) Glomeromycota, (**B**) symbiotrophs, (**C**) pathotrophs, and (**D**) nematode-trapping fungi across crop sequences. The phylum, trophic modes, and nematophagous guild shown all had centered log ratios (CLRs) that differed significantly by crop sequence according to ANOVA or the Kruskal-Wallis test (*P* < 0.05 after false discovery rate (FDR) correction) in multiple sampling timepoints. FDR-corrected *P*-values associated with differential abundance tests are shown on each plot. For consistency, data from all timepoints are shown, even if differences were non-significant. Error bars show one standard error above and below the mean. Crop sequences include 5 consecutive years of corn after the 5th year of soybean (C1-C5); 5 consecutive years of soybean after the 5th year of corn (S1-S5); soybean in annual rotation with corn (Sa); corn in annual rotation with soybean (Ca); long-term *Bt* corn (Cc) and SCN-susceptible soybean (Ss) monocultures; non-*Bt* Corn (Cn) and SCN-resistant soybean (Sr) monocultures.

**Figure 7.**
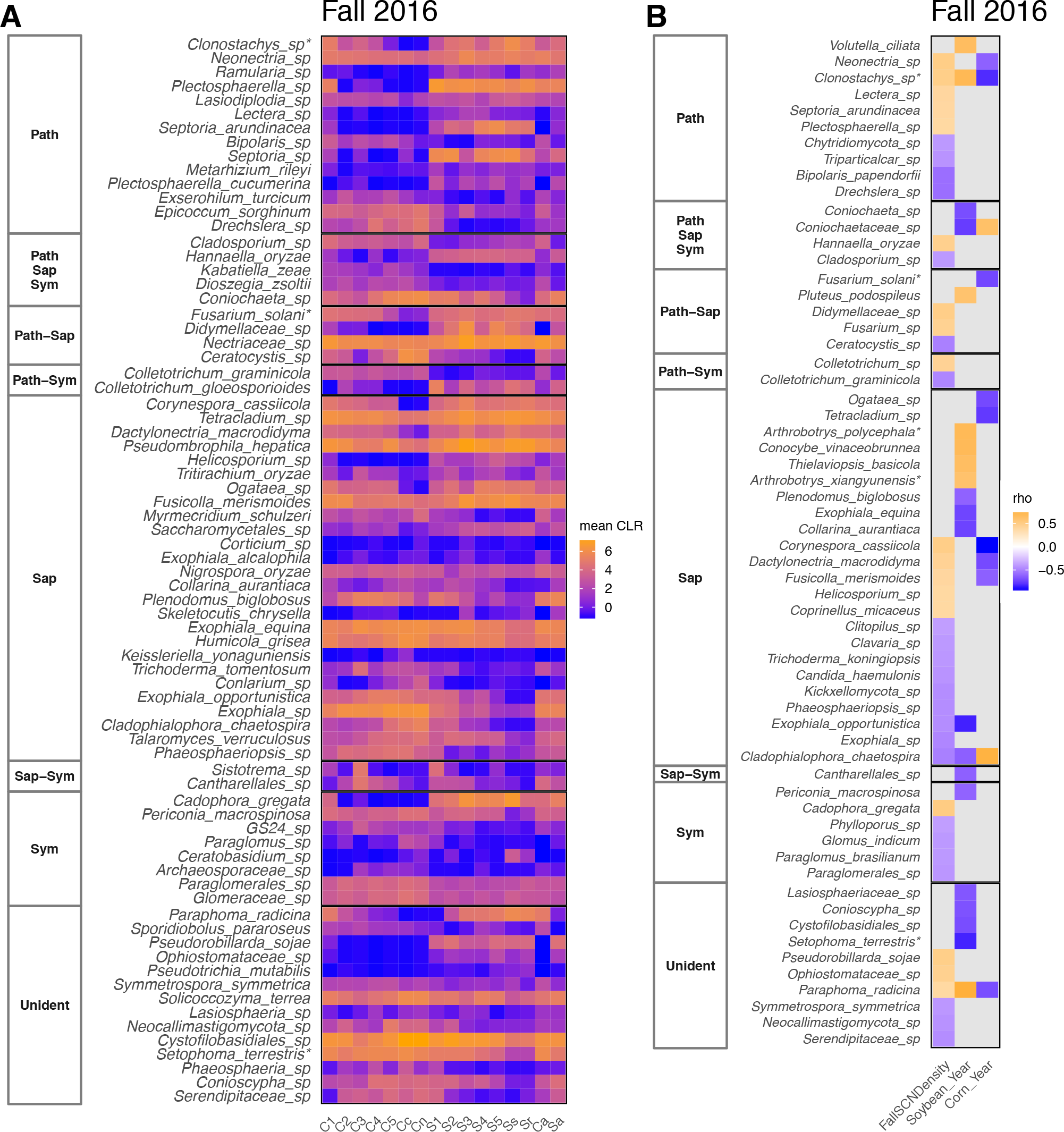
Differential abundance of fungal taxa by crop sequence in fall 2016. (**A**) Centered log ratios (CLRs) of fungal taxa that differed significantly by crop sequence according to ANOVA or the Kruskal-Wallis test (*P* < 0.05 after false discovery rate (FDR) correction). (**B**) Taxa that significantly correlated with SCN density or monoculture year of corn or soybean, according to Spearman’s rank correlation test. Trophic modes assigned to each taxon are given in the left-hand columns. **Path** = pathotroph; **Sap** = saprotroph; **Sym** = symbiotroph; **Unident** = unidentified trophic mode. Within each trophic mode category, taxa are ordered from top to bottom by strength of correlation with soybean monoculture year. Asterisks denote taxa belonging to nematophagous guilds, as defined in this study. Crop sequences include 5 consecutive years of corn after the 5th year of soybean (C1-C5); 5 consecutive years of soybean after the 5th year of corn (S1-S5); soybean in annual rotation with corn (Sa); corn in annual rotation with soybean (Ca); long-term *Bt* corn (Cc) and SCN-susceptible soybean (Ss) monocultures; non-*Bt* Corn (Cn) and SCN-resistant soybean (Sr) monocultures.

Phyla, trophic modes, and nematophagous guilds were identified that showed soybean-associated or corn-associated patterns. The most consistent pattern observed at the phylum level was for Glomeromycota, which was more abundant under corn monoculture in all but one sampling timepoint (Figure 6A). In terms of trophic modes, symbiotrophs showed a corn-associated pattern, whereas pathotrophs showed a soybean-associated pattern (Figure 6B,C). Among nematophagous guilds, nematode-trapping fungi displayed the clearest pattern, with the highest relative abundance associated with later years of soybean monoculture, including both susceptible (Ss) and SCN-resistant (Sr) soybean plots, and with early years of corn monoculture (Figure 6D). These patterns were also supported by significant spearman rank correlation tests showing significant positive correlations of relative abundance of these phyla, trophic modes, or nematophagous guilds with increasing years of soybean or corn monoculture at one or more sampling timepoints.

At the genus and species level, many fungi showed soybean-associated patterns of relative abundance (Figures 7, **S7, and S8**). Although Glomeromycota generally showed higher relative abundance under corn (Figure 6A), one AMF species, *Rhizophagus irregularis*, showed a soybean-associated pattern in midseason of both years (**Figure S7, S8**). Soybean-associated taxa also included several potential soybean pathogens, such as *Septoria arundinacea*, *Fusicolla merismoides* (Syn. *Fusarium merismoides*), and *Dactylonectria macrodidyma* (Malapi-Wight et al., 2015) (Figures 7, **S7, and S8**). Several fungi that have been shown to be capable of colonizing both soybean roots and SCN cysts also exhibited a soybean-associated pattern. These included *Paraphoma radicina* (Stiles and Glawe, 1989) and the soybean pathogens *Corynespora cassiicola* (Carris et al., 1986), *Cadophora gregata* (Syn. *Phialophora gregata*) (Carris et al., 1986), and *Fusarium solani* (Stiles and Glawe, 1989). Increasing years of soybean monoculture saw greater relative abundance for non-pathogenic soybean endophytes, such as *Plectosphaerella* sp. (Impullitti and Malvick, 2013), and *Pseudorobillarda sojae* (Uecker and Kulik, 1986). One species of Tremellomycete yeast, *Hannaella oryzae*, which commonly inhabits plant phyllospheres (Landell et al., 2014), showed a soybean-associated pattern at four sampling timepoints (Figures 7, **S7, and S8**).

Many nematophagous fungi showed soybean-associated patterns of relative abundance (Figures 7, **S7, and S8**). These included the nematode-trapping fungi *Orbilia auricolor*, *Arthrobotrys scaphoides*, *Athrobotrys polycephala*, and *Arthrobotrys xiangyunensis*, of which the latter two were also significantly positively correlated with SCN density at one and two sampling points, respectively (**Figure S8**). Among potential egg-parasites, *Clonostachys* sp. showed a strong soybean-associated pattern and was also significantly correlated with SCN density at three sampling timepoints (Figures 7B, **S8**). Several species of *Mortierella*, including *Mortierella elongata*, *Mortierella alpina*, and *Mortierella polygonia*, some isolates of which have been shown to inhibit SCN egg hatch (Juba et al., 2004), showed soybean-associated patterns and significant correlations with SCN density (**Figures S7, S8**). *Didymella americana* (syn. *Peyronellaea americana*), which has been isolated from nematode cysts (Aveskamp et al., 2010), was positively correlated with soybean monoculture year in midseason 2016, while an OTU less precisely identified as *Didymellaceae* sp. showed a soybean-associated pattern and significant correlations with SCN density at multiple timepoints (**Figure S8**). One basidiomycete species, *Coprinellus micaceus* (syn. *Coprinus micaceus*) was positively correlated with SCN density in fall 2016 and with soybean monoculture year in spring 2016 (Figures 7B **and S8**). Some members of *Coprinellus* are known to produce nematicidal compounds either as a defense against herbivory or, for nematophagous species, as part of a feeding strategy (Degenkolb and Vilcinskas, 2016; Plaza et al., 2016)

A contrasting group of fungi showed a corn-associated pattern (Figures 7, **S7, and S8**). Among symbiotrophic taxa, these included AMF identified as *Paraglomus brasilianum* and *Glomus indicum*, as well as the dark septate endophyte *Periconia macrospinosa* (Doran et al., 1984). A corn-associated pattern was also observed for OTUs identified as members of the family, Serendipitaceae, which is in the order Sebacinales (Figures 7, **S7, and S8**). Increasing years of corn monoculture saw greater relative abundance of several saprotrophic taxa, including *Helicosporium pallidum*, *Cyphellophora* sp. (Tedersoo et al., 2014), *Cladophialophora chaetospira*, *Exophiala equina*, *Talaromyces verruculosus*, *Coniochaeta ligniaria* (Jiménez et al., 2017), *Collarina aurantiaca*, *Myrmecridium phragmitis*, *Humicola grisea*, *Myrmecridium schulzeri* (Tedersoo et al., 2014), and *Apodus* sp. (Tedersoo et al., 2014). The plant pathogens *Cladosporium*, *Plenodomus biglobosus* (Marin-Felix et al., 2017), *Parastagonospora avenae*, *Parastagonospora nodorum*, *Phaeosphaeriopsis* sp., *Setophoma terrestris*, *Drechslera* sp., *Ceratocystis* sp., and *Phaeosphaeria* sp. all showed corn-associated patterns, as did one Tremellomycete yeast, a member of Cystofilobasidiales (Fell et al., 1999).

No significant correlations were found between the relative abundance of specific taxa and soybean yield at any sampling timepoint, but a few taxa showed significant correlations with corn yield in spring and midseason 2016 (**Figure S8**). In general, taxa that correlated positively with corn yield had greater relative abundance under continuous soybean monoculture. One of these taxa was classified as *Glomus aggregatum*, an AMF associated with soybean monoculture in midseason 2016, and another was identified as *Mortierella* sp. (**Figure S7 and S8**).

### 4.7 Differential abundance by season

Changes in relative abundance by season were generally more significant than by crop sequence, most likely due to the fact that data from multiple crop sequences were combined for these analyses, resulting in a larger sample size. The number of phylum-level discoveries ranged from 6 fungal phyla in 2016 soybean plots to 13 fungal phyla in 2016 corn plots (**Figure S9**). The number of species-level discoveries ranged from 77 taxa in 2016 soybean plots to 337 taxa in 2016 corn plots (**Table S15**). Unlike differential abundance by crop sequence, in which only two main patterns were evident, taxa, trophic modes, and nematophagous guilds exhibited a variety of seasonal shifts and showed similar seasonal patterns regardless of crop host (**Figure S9-S11**).

There was some consistency between shifts in relative abundance of taxa by crop sequence and shifts observed across seasons. For example, several species of corn-associated AMF, as well as the plant pathogen, *S. terrestris*, which increased in relative abundance over continuous corn monoculture, also increased over the growing season under corn, having greater relative abundance in midseason or fall compared to spring (**Table S15**). Soybean-associated taxa, including many potentially nematophagous fungi, like *Plectosphaerella cucumerina*, *Clonostachys* sp., *D. americana*, *Fusarium oxysporum*, *F. solani*, *M. chlamydosporia*, and several species of *Mortierella* (Hu et al., 2018), had greater relative abundance at either midseason or fall under soybean in 2015, and several others, including *A. scaphoides* and *Coprinopsis candidolanata*, decreased over the corn growing season in 2016 (**Table S15**). The soybean-associated AMF, *R. irregularis*, increased from spring to midseason under soybean in 2015 (**Table S15**). Similarly, the soybean endophyte, *P. sojae*, and the soybean pathogen, *C. gregata*, increased in relative abundance throughout the soybean growing season in 2015 and 2016 (**Table S15**). Other soybean-associated plant pathogens, like *S. arundinacea*, which consistently decreased under continuous corn monoculture, also decreased throughout a single corn growing season in 2016 (**Table S15**).

### 4.8 Correspondence between fungal taxa, yield, SCN density, and soil properties

Distance-based redundancy analysis showed that a combination of crop yield, SCN density, and soil properties explained, on average, 57% and 52% of variation between fungal communities at the ordinal level in corn plots and soybean plots, respectively. Corn yield and soil P varied with fungal communities in the same direction in 2015 and 2016 and corresponded with the relative abundance of Mortierellales, an order containing phosphate-solubilizing fungi (Li et al., 2018) (Figure 8A,B). A spearman correlation test revealed that the relative abundance of this order was positively correlated with soil P in 2015 (Table 3). While not immediately apparent on the redundancy analysis plots, the relative abundance of Sebacinales, an order containing ectomycorrhizal and endophytic species (Weiß et al., 2011, 2016), and two glomeromycete orders, Glomerales and Paraglomerales, were significantly negatively correlated with soil P (Table 3). SCN density and yield corresponded with soybean-associated fungal communities in opposite directions in 2016 (Figure 8C,D). Hypocreales, an order containing many nematophagous and entomopathogenic fungi (Castillo Lopez et al., 2014; Ghayedi and Abdollahi, 2013; Manzanilla-Lopez, Rosa H; Esteves, Ivania; Finetti-Sialer, Mariella; Hirsch, Penny; Ward, Elaine, Devonshire, Jean; and Hidalgo-Diaz, 2013; Stirling, 2014), corresponded with SCN density in both corn and soybean distance-based redundancy analysis plots (Figure 8).

**Figure 8.**
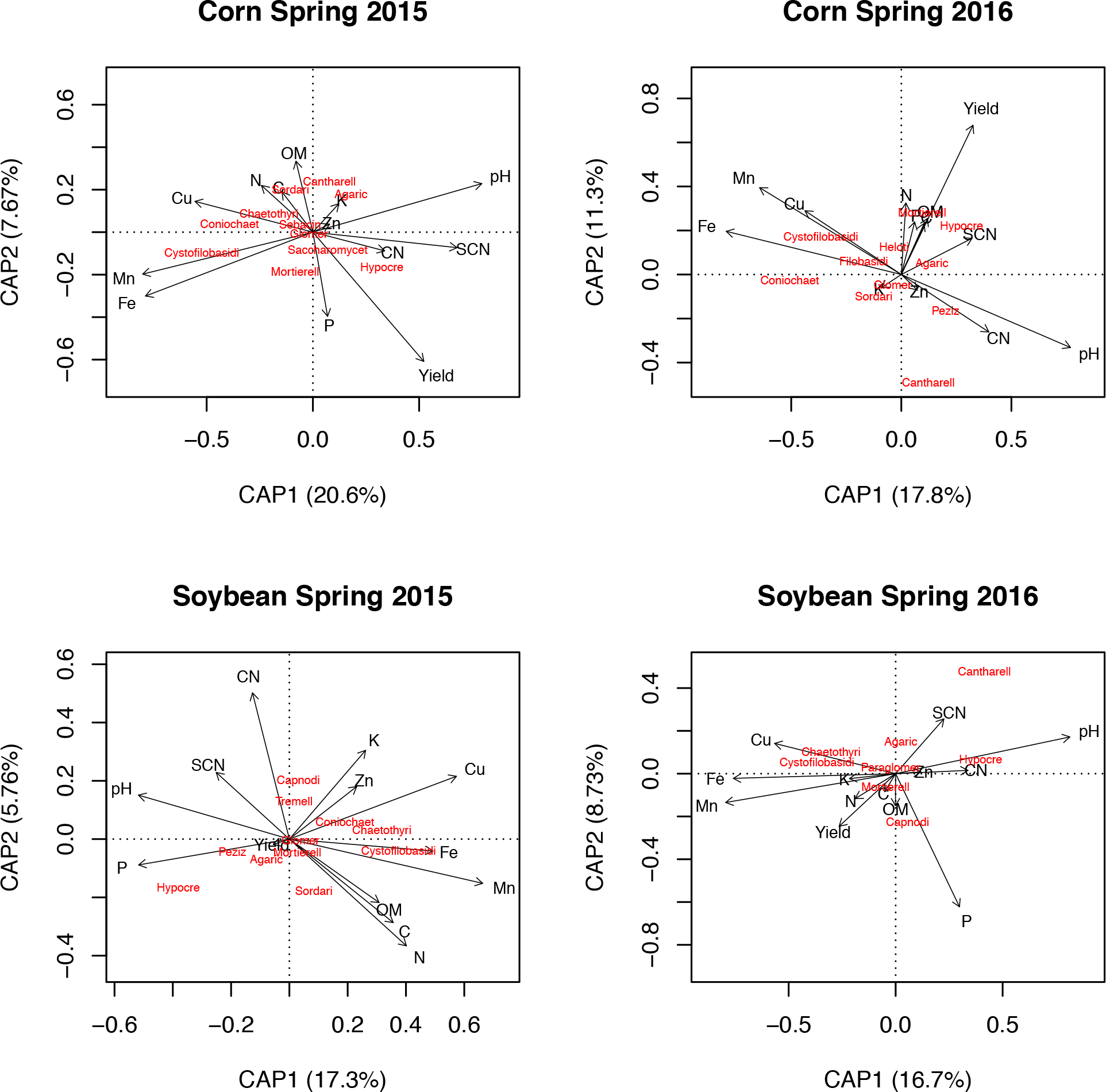
Distance-based redundancy analysis (dbRDA) of fungal communities for (**A**) corn plots in 2015, (**B**) corn plots in 2016, (**C**) soybean plots in 2015, and (**D**) soybean plots in 2016. Relative position of fungal orders in the biplots is based on Bray-Curtis dissimilarity of square root transformed relative abundance at the ordinal level. Vectors indicate the weight and direction of soil properties, soybean cyst nematode (SCN) density, and crop yields associated with fungal communities. The dbRDA axes describe the percentage of the variation explained by each axis while being constrained to account for differences in soil properties, SCN density, and yield. Order names were shortened by removing “ales” from the end of each name. Orders that mapped near the plot center or that overlapped, including Glomerales, Paraglomerales, and Sebacinales on some plots, were omitted to aid readability. C = carbon; Cu = copper; Fe = iron; K = potassium; Mn = manganese; N = nitrogen; OM = organic matter; P = phosphorus; CN = carbon:nitrogen; SCN = soybean cyst nematode.

**Table 3.**
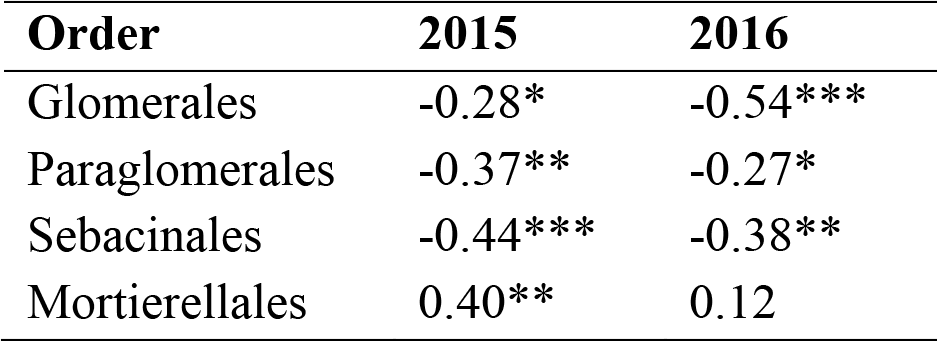
Spearman correlations (*rho*) between centered log ratios (CLRs) of four fungal orders and soil phosphorus in spring of 2015 and 2016. Significant relationships are reported at FDR-corrected *P*-values of \**P* < 0.05, \*\**P* < 0.01, and \*\*\**P* < 0.001.

## 5. DISCUSSION

Utilization of a long-term research site at which crops have been planted in continuous rotations and monoculture since 1982 allowed us to gain unique insights into the effects of crop sequences on soil fungi. Shifts in the composition of fungal communities correlated with years of continuous monoculture of soybean or corn, providing further evidence for the role of plant hosts in shaping soil microbial communities (Berg et al., 2005; Kuske et al., 2002; Peiffer et al., 2013). Overall soil fungal alpha diversity was higher under corn monoculture, but the diversity of specific functional guilds showed contrasting patterns, with saprotrophs and symbiotrophs increasing in diversity under continuous corn and nematode-trapping and egg parasitic fungi increasing in diversity under continuous soybean. Several potential predators and opportunistic pathogens of SCN were identified whose relative abundance was positively correlated with SCN density and with years of continuous soybean monoculture. These predominantly included members of nematode-trapping and egg-parasitic guilds but also included fungi that may immobilize or kill nematodes through the production of nematicidal compounds. Soil chemical properties, most notably P, shifted over continuous monoculture of both crop hosts and corresponded with shifts in fungal taxa, including several fungal orders involved in P solubilization and acquisition (Mortierellales, Glomerales, Paraglomerales, Sebacinales). Changes in soil properties, SCN density, and the relative abundance of specific fungal groups, like AMF and plant-pathogenic fungi, suggest multiple causes of yield declines under continuous monoculture of soybean and corn.

### 5.1 Yield and soybean cyst nematode density

The results of corn yield (Figure 1A) were similar to those of a metanalysis of hundreds of thousands of fields in the United States (Seifert et al., 2017) and to a previous study at our site in which a yield penalty was observed during the second year of corn monoculture, with insignificant yield penalties with additional years of corn (Grabau and Chen, 2016b). Decreased nitrogen availability has been identified as the main limiting factor in corn yields under continuous corn cropping (Gentry et al., 2013). However, in our study total N actually increased under continuous corn monoculture (**Table S7**), most likely due to N fertilization with urea in the corn plots. The most significant change in any soil property was seen for P, which decreased under corn monoculture and increased under soybean monoculture, a result similar to one observed in a previous study at the same site (Johnson et al., 1991). While a significant correlation between soil P and corn yield was not detected (**Table S8**), corn yield and soil P corresponded with fungal communities in the same direction (Figure 8A,B), raising the possibility that corn yield was in fact limited, at least in part, by available P. One notable difference between our results and those of Grabau and Chen (2016b) was that non-*Bt* corn yields were lower in 2015 and higher in 2016 compared to long-term continuous *Bt*-corn (Figure 1A). One possible explanation for this observation is that pressure from plant parasitic nematodes or insect pests was higher in 2015, allowing *Bt* corn to out-perform non-*Bt* corn (Grabau and Chen, 2016b). However, this hypothesis cannot be tested, because measurements of plant-parasitic nematodes or insect pests other than SCN were not taken for this study.

Soybean yields in our study (Figure 1B) were somewhat less consistent with previous reports. In the same metanalysis of corn and soybean fields in the United States, soybean yields declined each year for up to five years of consecutive monoculture, without the yield penalty leveling off as for corn (Seifert et al., 2017). A similar pattern was observed in a previous study at our site, with the lowest soybean yields observed for long-term SCN-susceptible soybean monoculture (Ss) (Grabau and Chen, 2016a). By contrast, our results showed relatively stable yields in years 1 through 5 of continuous soybean monoculture (Figure 1B). Our findings were consistent, however, with Grabau and Chen’s (2016a) study in that SCN-resistant soybean (Sr) had higher yields than SCN-susceptible soybean in long-term monoculture, especially in 2015 (Figure 1B), the year in which SCN density was greater (Figure 1C). Interestingly, this is the same year in which *Bt* corn outperformed non-*Bt* corn, which could suggest that weather conditions or some other factor associated with this growing season encouraged reproduction of plant parasitic nematodes or insect pests affecting both soybean and corn. SCN has been proposed to be the single most limiting factor for soybean yield (Seifert et al., 2017) and has previously been shown to be negatively correlated with soybean yields at this site (Grabau and Chen, 2016b). Our results showing significant negative correlations between SCN egg density and susceptible soybean yields (Figure 1D) corroborate these findings.

Overall, the pattern of SCN density observed in this experiment (Figure 1C) supports the practice of multi-year rotations with corn to manage SCN populations. SCN levels in soybean annual rotation plots were comparable to those in long-term soybean monoculture and only declined significantly after four or five years of corn monoculture in 2015 and 2016, respectively (Figure 1C), a more extreme result than an earlier report at the same site that showed significant declines in SCN density within the first year of corn (Porter et al., 2001). SCN may become more problematic with a warming climate that favors longer growing seasons, resulting in more generations of this polycyclic pathogen per year (Boland et al., 2004; Chen, 2011). The practice of annual rotation, which resulted in a significant improvement in corn yields (Figure 1A), may no longer be optimal for improving yields of soybean due to its failure to regulate SCN. Alternatively, it is possible for long-term soybean monoculture to lead to the build-up of microbes that antagonize SCN, resulting in SCN-suppressive soil (Chen, 2007). While our study showed that this build-up occurs, it cannot provide evidence that soil fungi helped to control SCN densities, due to the nature of the study design. However, a nearby field at the same experimental site with a similar long-term soybean monoculture history showed evidence of biological suppression of SCN (Hu et al., 2017), and it is possible that such suppression is also occurring in the long-term soybean monoculture plots used in our study.

### 5.2 Fungal communities shaped by long-term monoculture

Analysis of fungal community structure revealed that with increased years of monoculture, soil fungal communities become more similar to those of the long-term monoculture plots associated with their respective crop hosts (Figure 2A). This finding is in agreement with other studies showing this same phenomenon for other crops, including coffee (Zhao et al., 2018) vanilla (Xiong et al., 2014), *Pseudostellaria heterophylla* (Wu et al., 2016), and soybean (Bai et al., 2015). These shifts in bulk soil communities may be related to shifts in soil properties that favor the proliferation of certain groups of fungi (Lauber et al., 2008), or they could be due to the “rhizosphere effect” in which certain groups of microbes proliferate in the soils associated with plant roots (Hiltner, 1904; Kristin and Miranda, 2013; Lapsansky et al., 2016). Our research suggests that many factors, including soil properties, plant hosts, and even plant-parasitic nematodes like the SCN help shape the fungal community over continuous monoculture. These results add to the growing body of evidence that long-term monoculture has a dramatic impact on soil microbial communities.

### 5.3 Alpha diversity affected by crop sequence

Plant species richness is a predictor of soil fungal richness (Maltz et al., 2008; Tedersoo et al., 2014), and a greater variety of root exudates can support a more diverse fungal community (Broeckling et al., 2008). We expected, therefore, that annual rotation (Ca and Sa) would result in increased soil fungal alpha diversity. However, total alpha diversity was more strongly associated with corn monoculture than with crop rotation (Figure 3A **and Table S9**). This pattern was driven, in part, by the diversity of symbiotrophic and saprotrophic fungi (Figure 3B). Our result showing increased diversity of symbiotrophs under corn monoculture is consistent with an earlier study at the same site documenting higher diversity of AMF under corn monoculture compared to soybean (Johnson et al., 1991). This result may imply that corn has fewer selective barriers to AMF colonization than soybean. Alternatively, a more diverse AMF community may be the result of greater below-ground biomass in corn (Johnson et al., 1991). A larger root system may support more AMF, overall, resulting in greater detection of diverse AMF in our metabarcoding analysis. The increased diversity of saprotrophs could also be related to biomass, as a greater amount of crop residue, which promotes the proliferation of diverse saprotrophic fungi (Ma et al., 2013), remains after corn harvest compared to soybean harvest (Doran et al., 1984). Our data showing both greater diversity of saprotrophic fungi (Figure 3B **and Table S9**) and greater amounts of organic matter (**Table S7**) under corn monoculture support this hypothesis.

While overall alpha diversity was higher under corn monoculture (**Table S9**), nematode-trapping and egg-parasitic nematophagous guilds showed increased alpha-diversity under soybean monoculture, correlating with higher egg densities of SCN (Figure 3C, **Tables S9 and S10**). This finding is consistent with other ecological studies that show an increase in bacterial and fungal diversity in response to increased availability of limited resources (Cline et al., 2018; She et al., 2018). Further research is needed to understand whether these fungi contribute to controlling SCN populations or if their merely act as opportunistic SCN parasites without substantially reducing SCN density.

Our findings regarding total alpha diversity (Figure 3A **and Table S9**) suggest another potential mechanism through which crop rotation benefits soybean. General suppression of soil-borne plant pathogens relies on a diverse fungal community that limits the establishment of pathogenic microbes in the soil (Garbeva et al., 2011). It is possible that corn monoculture helps create such a community by increasing the taxonomic and functional diversity of soil fungi, a pattern that may also contribute to soil fertility (Mäder et al., 2002). It is also possible that increased AMF diversity under corn monoculture contributes, in part, to higher soybean yields following years of corn (Johnson et al., 1991, 1992).

Seasonal variation, including an increase in bulk soil fungal alpha diversity (Chao1) from midseason to fall in 2016 (Table 2), was consistent with the pattern observed for alpha diversity of fungal communities in SCN cysts in 2015 and 2016 at the same study site (Hu et al., 2018). It is possible that these related patterns reflect an ecological relationship between bulk soil and SCN cyst fungal communities, given that soil may act as a reservoir for microbial taxa (Mendes et al., 2014) from which certain groups of microbes can be selectively filtered by niche environments (Kristin and Miranda, 2013), like SCN cysts. Unexpectedly, the opposite pattern was observed in 2015, in which alpha diversity in the fall was significantly lower than midseason (Table 2). It is possible that the lower alpha diversity in fall 2015 was due to abnormal weather or environmental conditions that introduced a sampling bias. The fall of 2015 was one of the hottest on record, with temperatures above 80o F, no precipitation the week prior to sampling, and significantly higher solar radiation compared to fall 2016 (Regents of the University of Minnesota, 2018). It is possible that some fungi or fungal DNA that would otherwise have been present in the soil was destroyed by excessive heat, dryness, and solar radiation at this sampling timepoint.

### 5.4 Arbuscular mycorrhizal fungi

Earlier studies at our experimental site found that unique communities of AMF proliferated under continuous corn and soybean monoculture and that the proliferation of specific AMF taxa was correlated with yield declines in their respective crop hosts (Johnson et al., 1991, 1992). Johnson (1992) hypothesized that one reason for these yield declines was that AMF that proliferated under monoculture were less mutualistic than AMF sustained by crop rotation. While it is probable that our metabarcoding approach failed to capture the full diversity of AMF (Kohout et al., 2014; Stockinger et al., 2010), our results nonetheless supported the idea that corn and soybean host unique AMF communities (Figures 7, **S7, and S8**). In further support of Johnson’s (1992) hypothesis is the observation that OTUs identified as Glomeraceae sp., which proliferated under corn, were negatively correlated with corn yield at one timepoint, whereas OTUs identified as *Glomus aggregatum*, which showed a soybean-associated pattern, were positively correlated with corn yield (**Figure S8**). These findings support the idea that soybean-associated AMF are more mutualistic on corn than are corn-associated AMF.

However, it may be possible for AMF to be correlated with corn yield declines without being the cause of these declines. Our results suggest an alternative hypothesis that specific AMF proliferated under corn monoculture as an adaptation to lower soil P (**Table S7**). In this model, both AMF and soil P are correlated with yield declines in corn. However, it is possible that yield declines would be more severe without AMF, which help the plant take up the limited soil P. This model is in agreement with studies showing that corn yields and uptake of P are higher if the preceding crop is corn rather than a non-mycorrhizal crop, such as canola (Miller, 2000). In order to address these contrasting hypotheses, further studies testing the efficiency with which soybean and corn-associated AMF uptake and translocate P and affect the yield both crop hosts are warranted.

### 5.5 Nematophagous fungi

Of the three nematophagous guilds that were examined, nematode-trapping fungi had the strongest relationships with crop sequence and SCN density (Figures 3C, **6D and S6**). A pioneering experiment on fungal-nematode interactions showed that nematode-trapping fungi build up in soil with a large quantity of free-living nematodes and may provide a form of biological control against plant parasitic nematodes under such conditions (Linford, 1937). Subsequent studies failed to corroborate this conclusion (Stirling, 2014). However, a density-dependent host-parasite relationship, which is an important trait for biocontrol agents of plant-parasitic nematodes (Jaffee et al., 1992), has been observed experimentally for several species of trapping fungi, including *D. ellipsospora* (syn. *M. ellipsosporum*), *A. dactyloides* (Syn: *D. dactyloides*) (Jaffee et al., 1993), and *Dactylellina haptotyla* (Jaffee, 2003). Trapping fungi that produce adhesive knobs (e.g. *D. ellipsospora*) or constricting rings (e.g. *A. dactyloides*) have been shown to respond to an increase in nematodes as a food source more than trapping fungi that produced adhesive networks (e.g. *Arthrobotrys oligospora*) (Jaffee, 2003). However, our findings did not corroborate these results. Instead, we observed increased relative abundance of adhesive-network forming fungi, including *Arthrobotrys superba*, *A. polycephala* (Yu et al., 2014), and *A. xiangyunensis* (Liu et al., 2014a) under consecutive years of soybean monoculture (Figures 7, **S7, and S8**). Of the trapping fungi we detected, only *A. polycephala* and *A. xiangyunensis* showed significant positive correlations with SCN egg density (Figures 7B **and S8**).

One of the more interesting findings was that the relative abundance of nematode-trapping fungi in SCN-resistant soybean plots was nearly as great as in long-term susceptible soybean monoculture plots (Figure 6D). One possible explanation for this observation is that the plant host, rather than SCN density, is responsible for the build-up of trapping fungi in the bulk soil (Bordallo et al., 2002). However, another possible explanation is that the abundance of trapping fungi DNA in bulk soils of SCN-resistant soybean plots is due to a soil memory effect (Lapsansky et al., 2016). From 1982 until 2010, these plots were planted with SCN-susceptible soybean and therefore likely had a sustained build-up of SCN cysts and, possibly, predators of SCN, like nematode-trapping fungi. These fungi could remain active as soil saprobes or merely be present as large quantities of spores even after a decline in SCN cyst density.

The nematode egg parasite guild is difficult to define for several reasons. Dozens of species of fungi have been isolated from SCN cysts (Chen and Chen, 2002), but far fewer have been tested for their ability to parasitize eggs *in vitro*. It is, therefore, possible that many of these fungi grow saprotrophically on dead nematode eggs and are not true parasites that infect living eggs. For this study, we chose to define the egg parasite guild as fungal species that have been isolated from nematode cysts and been shown to parasitize nematode eggs in an *in vitro* bioassay. Thus, we have taken a conservative approach, and it is possible that this guild includes far more taxa than we included. In addition to the difficulty in defining the egg parasite guild, it is also difficult to draw conclusions about the reasons for changes in relative abundance or alpha diversity of this guild over various crop sequences (Figure 3C **and Table S14**). Nematode egg parasites often occupy multiple ecological niches. For example, isolates of *F. solani* can act both as plant pathogens (Rupe et al., 1997) and soil saprotrophs (O’Donnell et al., 2008), and some isolates have been shown to parasitize SCN eggs (Chen and Chen, 2003). It is possible that egg parasite diversity increases under soybean monoculture (Figure 3C) because the plant host, rather than the SCN, is hosting these pathogens. Nonetheless, it is intriguing that many fungi that have been isolated from SCN cysts showed positive correlations with SCN density. For example, the well-characterized nematode egg parasite *P. lilacinum* (Song et al., 2016b) was below the limits of detection in many corn monoculture plots but was consistently detected in long-term susceptible soybean monoculture plots (Ss) (Figure 5B). Likewise, potential SCN egg parasites *C. cassiicola*, *P. radicina*, *Clonostachys* sp., and *D. macrodidyma* all showed consistently positive correlations with SCN density across multiple timepoints (Figures 7B **and S8**). Interestingly, two *Mortierella* species, *M. alpina* and *M. elongata*, which were shown to have strong correlations with SCN density when quantified from SCN cysts in our same study system (Hu et al., 2018), also showed significant correlations with SCN density in bulk soil (**Figure S8**). Some *Mortierella* species isolated from SCN cysts have demonstrated a relatively high egg parasitism index, a measure of the percentage of eggs colonized within a cyst (Chen and Chen, 2002). Collectively, these findings suggest that some *Mortierella* species may be parasitic on SCN. However, an alternative hypothesis supported by our results is that the soybean plant promotes the proliferation of *Mortierella* species to improve P acquisition (**Table S9**). Future research testing both of these hypotheses is warranted.

Endoparasites of SCN, such as *H. rhossiliensis* and *Hirsutella minnesotensis*, are commonly isolated from SCN J2s in soybean fields in Minnesota (Liu and Chen, 2000), and the percentage of J2s parasitized by *H. rhossiliensis* has been shown to be affected by crop sequence in a culture-based study at our experimental site (Chen and Reese, 1999). However, our study failed to detect a clear association between this nematophagous guild and crop sequence or SCN density. This may be due to the fact that these fungi make up an extremely small proportion of the entire bulk soil fungal community (Figures 4C **and 5C**), causing their DNA to be below the limits of detection in most bulk soil samples.

### 5.6 Relationships between soil properties, crop sequences, and fungal communities

Even though a majority of soil properties did not differ significantly by crop sequence, the general trend showed that most soil properties increased either under soybean or corn monoculture, with OM, Fe, Mn, Cu, total N, and TOC associated with corn and pH and P associated with soybean (**Table S7**). Many of these changes are likely associated with inputs from fertilizer or from the crops, themselves. Corn, a plant with greater overall biomass than soybean, contributes more organic matter and carbon to the soil (Karlen et al., 1994). While soybean was expected to be associated with higher levels of N due to the effect of nitrogen-fixing rhizobia (Cooper, 2007), the urea fertilizer used in corn plots resulted in greater total N under corn (**Table S7**). Shifts in pH across crop sequences may also be related to urea fertilizer, which makes soil more acidic (Lungu and Dynoodt, 2008).

However, no phosphorus-containing fertilizer was applied to the plots in this study, so a biological cause for differences in P levels is likely. The decline in P under corn monoculture may be related to the associated increase in relative abundance of Glomeromycota and Serendipitaceae sp. (Figures 6, 7, **and S6-S8**). AMF substantially increase the ability of plants to acquire P, but in doing so deplete P from surrounding soil (Zhang et al., 2010). When a portion of the corn plant is removed at harvest, this P is not replaced. Sebacinales, the order containing Serendipitaceae sp., appeared to display the same pattern as the glomeromycete orders (Figure 8 **and Table S9**), a result that may be reflective of a similar ecological function. One well-studied member of Sebacinales, *Serendipita indica* (syn. *Piriformospora indica*), is associated with increased plant levels of N, K, and P (Kumar et al., 2012) and has been shown to actively transport P to corn roots (Yadav et al., 2010). Collectively, these findings suggest that the corn-associated fungal community may help corn to adapt to a low-P environment. Conversely, the correspondence between the soybean-associated order Mortierellales and soil P (Figure 8 **and Table S9**) may be a result of these fungi converting inorganic or organic phosphates into the bioavailable form detected in our assay (Zhang et al., 2011). Higher P levels brought about by the activity of mortierellalean fungi may explain, in part, the improved corn yield following years of soybean. However, it is also possible that the positive correlation between the relative abundance of *Mortierella* sp. and corn yield (**Figure S8**) is related to other mechanisms, such as the impact of *Mortierella* sp. on plant growth-promoting hormones (Li et al., 2018).

## 6. CONCLUSIONS

In this study, we showed that continuous soybean and corn monoculture is associated with significant changes in bulk soil fungal communities, with specific functional groups of fungi proliferating under one or the other crop host. Our hypothesis that fungal communities would become progressively similar to corresponding communities in long-term monoculture plots was supported. However, the hypothesis that crop rotation would result in greater fungal diversity was not supported. Instead, alpha diversity was found to increase in response to corn monoculture rather than to crop rotation. The greatest impact of corn monoculture on soybean yield was arguably its effect on SCN density, which decreased over consecutive years of corn and increased over consecutive years of soybean. Our hypothesis that the diversity and relative abundance of nematophagous fungi would track the population density of SCN was supported for nematode-trapping fungi and for many potential nematode egg parasites, suggesting that members of these guilds hold promise as biocontrol agents against SCN. Relationships between corn monoculture, soil P, and AMF and Sebacinales hint at the adaptive role of these fungi in P acquisition, whereas relationships between soybean monoculture, Mortierellales, and soil P may help explain the increased yields of corn in rotation with soybean. Future research should test nematode-trapping and nematode egg-parasitic isolates for their potential to control SCN populations and address the roles of corn and soybean-associated AMF, sebacinalean, and mortierellalean fungi in P solubilization and uptake. Our findings serve to illustrate the complexity of relationships between crop sequences, soil fungal communities, plant parasitic nematodes, and soil properties and provide fundamental knowledge that can be used to guide future strategies to manage SCN and improve corn and soybean yields.

## AUTHOR CONTRIBUTIONS

NS assisted in collection of soil samples and DNA isolations, processed Illumina metabarcoding data, developed bioinformatics and statistical approaches, analyzed the data, and wrote the manuscript. KEB co-conceived of, supervised and provided guidance on the research, and edited the manuscript. WH helped collect cysts and soil samples, provided training and analytical pipelines for analyzing the data, and edited the manuscript. DH helped collect cysts and soil samples and edited the manuscript. SC co-conceived of and supervised the project, maintained the long-term research study plot site, and provided expertise on SCN biology.

## ABBREVIATIONS

AMF: arbuscular mycorrhizal fungi
CLR: centered log ratio
ITS1: internal transcribed spacer 1
OTU: operational taxonomic unit
SCN: soybean cyst nematode

## ACKNOWLEDGEMENTS

Fungal cultures and tissue for mock communities were generously supplied by Todd Burnes, Peter Kennedy, Senyu Chen, Harold Corby Kistler, and Timothy James. The authors would like to thank Wayne Gottschalk, Cathryn Johnson, Jeff Ballman, and undergraduate workers for their help in collecting soil samples and Bryani Lee for her assistance in processing soil samples and extracting DNA. We also thank Nicholas Dunn at the Minnesota Supercomputing Institute and Jon Palmer for help with implementing AMPtk, and Gabriel Al-Ghalith for help with statistical analysis using centered log ratios. We would also like to acknowledge Daryl Gohl, Allison MacLean, and Corbin Dirkx at the University of Minnesota Genomics Center for sharing their expertise and methods development in amplification and sequencing of fungal DNA for metabarcoding experiments.

## SUPPLEMENTARY MATERIAL

### Supplementary Materials and Methods

Supplementary code, find.longest.taxonomy.R, can be found in the following GitHub repository: https://github.com/stro0070/OTU-taxonomy-assignment

Figure S1 | Rarefaction curves for bulk soil samples.

Figure S2 | Community profiles of fungal mock community.

Figure S3 | Principle coordinates analysis of technical replicates.

Figure S4 | SCN Density by season for each crop host.

Figure S5 | Community profiles of ecological guilds assigned by FUNGuild.

Figure S6 | Spearman correlations between phyla, trophic modes, and nematophagous guilds and SCN density, monoculture year, and yield.

Figure S7 | Differential abundance of taxa by crop sequence.

Figure S8 | Spearman correlations between taxa and SCN density, monoculture year, and yield.

Figure S9 | Phylum differential abundance by season.

Figure S10 | Trophic mode differential abundance by season.

Figure S11. Differential abundance of nematophagous guilds by season

Table S1 | Corn and soybean crop sequences

Table S2 | Mock community 5 composition.

Table S3 | Taxonomy assignments.

Table S4 | Mock community taxa identified by bioinformatics pipeline.

Table S5 | OTUs detected in synthetic mock community sample.

Table S6 | Nematophagous guilds.

Table S7 | Mean values for soil properties within crop sequences

Table S8 | Correlations between soil properties and corn and soybean yields Table S9 | Alpha diversity metrics by crop host

Table S10 | Spearman correlations between alpha diversity and SCN, monoculture year, and yield

Table S11 | Taxonomic composition of trophic modes assigned by FUNGuild.

Table S12 | Taxonomic composition of ecological guilds assigned by FUNGuild.

Table S13 | Nematophagous taxa detected and their assignment to nematophagous guilds.

Table S14 | Differential abundance of fungal phyla, trophic modes, and nematophagous guilds across crop sequences

Table S15 | Differential abundance of taxa by season.

