## Supplementary Materials and Methods for "Fungal Communities in Soybean Cyst Nematode-Infested Soil under Long Term Corn and Soybean Monoculture and Crop Rotation"

**Supplementary text**

Preparation of fungal mock communities

Inclusion of a biological mock community is essential to assess the accuracy of high throughput amplicon sequencing (Nguyen et al., 2015). Our biological (fungal) mock community contained 36 fungal isolates from across several major phyla (Ascomycota, Basidiomycota, Chytridiomycota, and Mucoromycota) and included both closely related and more distantly related taxa to optimize the accuracy and precision of taxonomy assignments (**Table S2**). For fungal mock community samples, DNA from fungal tissue was extracted and the full ITS region was amplified using a REDExtract-N-Amp™ Tissue PCR Kit (Sigma-Aldrich) and primers ITS1f and ITS4. Sanger sequencing was performed at MCLAB (San Francisco, CA, USA) and sequences trimmed and aligned in BioEdit (Hall, 1999). Taxonomy was confirmed by performing BLAST searches against the UNITE and NCBI databases (Altschul et al., 1990; Benson et al., 2013; Kõljalg et al., 2005). Equal quantities of DNA (100 ng) were added for all taxa except for 2 isolates from which only a small amount of DNA was successfully extracted. For these isolates, only 10 ng of DNA was added. Twenty ng of the pooled mock community DNA was submitted along with bulk soil samples from midseason and fall 2015 for amplification and sequencing. For remaining sequencing runs, mock community DNA was amplified only once, and this PCR product was included in sequencing runs for spring 2015 and all 2016 bulk soil samples. A synthetic mock community was also included in the 2016 run to control for barcode bleeding (Palmer et al., 2018).

Taxonomy Assignment

A synthesis of four different methods was used for taxonomy assignment. In the first method, taxonomy was assigned using AMPtk's default "hybrid" option. ITS sequences from isolates of *Hirsutella rhossiliensis*, *Hirsutella minnesotensis*, *Arthrobotrys oligospora*, *and Metacordyceps chlamydosporia* from the fungal mock community, as well as the twelve synthetic mock community sequences (Palmer et al., 2018), were added to the database "on the fly" during this step. The other three methods used the "consensus BLAST" classifier in QIIME2 (Bokulich et al., 2018; Camacho et al., 2009; Caporaso et al., 2010). However, due to the scarcity of sequences for some taxa in the QIIME-formatted UNITE database (UNITE Community, 2017b), the full UNITE + INSD database (release date December 1, 2017) (UNITE Community, 2017a) was used, instead. In order to use this database with QIIME2, a custom mapping file was created to map sequence identifiers to taxonomy assignments in the UNITE + INSD database. Taxonomy was assigned to OTUs using three different sets of parameters in consensus BLAST. All parameter sets required 90% identity and that a minimum of 60% of BLAST hits showed the same taxonomy assignment. However, they differed in number of hits retained for consensus. The first consensus BLAST parameter set saved the top 20 hits (Consensus 20), the second parameter set saved the top 10 hits (Consensus 10), and the last parameter set saved only the best BLAST hit (Consensus 1).

All taxonomy assignments from these different taxonomy assignment methods were saved in a text file (**Table S3**) and run through a custom R script, "find.longest.taxonomy.R," (available at https://github.com/stro0070/OTU-taxonomy-assignment) in order to assign the deepest level of taxonomy to OTUs. These methods were first tested on the fungal mock community and priority was given to taxonomy assignment methods that more accurately and precisely identified known taxa. The priority order was: Consensus 20, AMPtk, Consensus 10, Consensus 1. This hybrid approach to taxonomy assignment was developed because no single taxonomy assignment method identified all OTUs in our fungal mock community and because the best BLAST hit method made several mistakes in taxonomy assignment that were not made by AMPtk or consensus BLAST methods. Using a hierarchical combination of these methods, we were able to identify 100% of fungal mock community members in one mock community PCR product (**Table S4**).

OTU Processing and Quality Control

After filtering, sequence depth for bulk soil samples ranged from 16,435 to 205,157 reads with an average sequence depth of 63,947. Sequence depth for negative controls ranged from 12 to 5491, but the median sample depth for negative controls was 68. One outlier, a water negative control supplied by the sequencing center, had 5,491 reads. The remaining negative controls all had fewer than 1000 reads. The number of OTUs in bulk soil samples ranged from 166 to 1,533, with 837 as the mean number of OTUs per sample. Rarefaction curves for bulk soil samples leveled off around 30,000 reads (**Figure S1**).

Our pipeline was able to capture a majority of mock community fungal taxa, with 100% of taxa being identified in sample "mk5-S3" and 81% and 83% of taxa identified in samples "mk5-S117" and "mk5," respectively (**Table S4**). The number of taxa identified varied depending on which PCR product was used for sequencing. One PCR product was used for mock community "mk5-S3," whereas a different PCR product was used for mock communities, "mk5-S117" and "mk5," which were included in two separate sequencing runs. The latter two mock communities displayed remarkable homogeneity in terms of relative abundance of taxa identified (**Figure S2**). However, a greater percentage of taxa were identified in sample "mk5-S3" (**Table S4**). This result suggests that inter-run variation was low, whereas variation between amplifications was high.

Despite filtering for index bleed in the AMPtk pipeline and removing OTUs with fewer than 10 reads across the entire OTU table, several unexpected taxa, the most abundant being *Phaeocalicium polyporaeum*, remained in the fungal mock community samples (**Table S4**). Applying more stringent filters eliminated an unacceptable number of expected taxa from our fungal mock communities, so more stringent filters were not applied. *P. polyporaeum* was not present in negative controls or in any environment samples, so it is probable that it represents a true contaminant, possibly from herbarium fungal material used in preparation of some samples in the mock community.

Ten out of twelve synthetic mock community sequences were detected in our synthetic mock community sample after index-bleed filtering (**Table S5**) (Palmer et al., 2018). One contaminant, classified as *Amanita*, was also present in extremely low abundance. This OTU was absent from all other samples and may represent a true contaminant in the synthetic mock community sample.

Technical replicates were prepared by amplifying and sequencing DNA extracted from multiple subsamples of soil from the same bulk soil sample. These resulted in communities that were remarkably homogeneous, with 62% of variation between fungal communities being explained by the sample of origin (**Figure S3**).
