## Supplementary figures and images for "Fungal Communities in Soybean Cyst Nematode-Infested Soil under Long Term Corn and Soybean Monoculture and Crop Rotation"

### Supplemental Figure 1

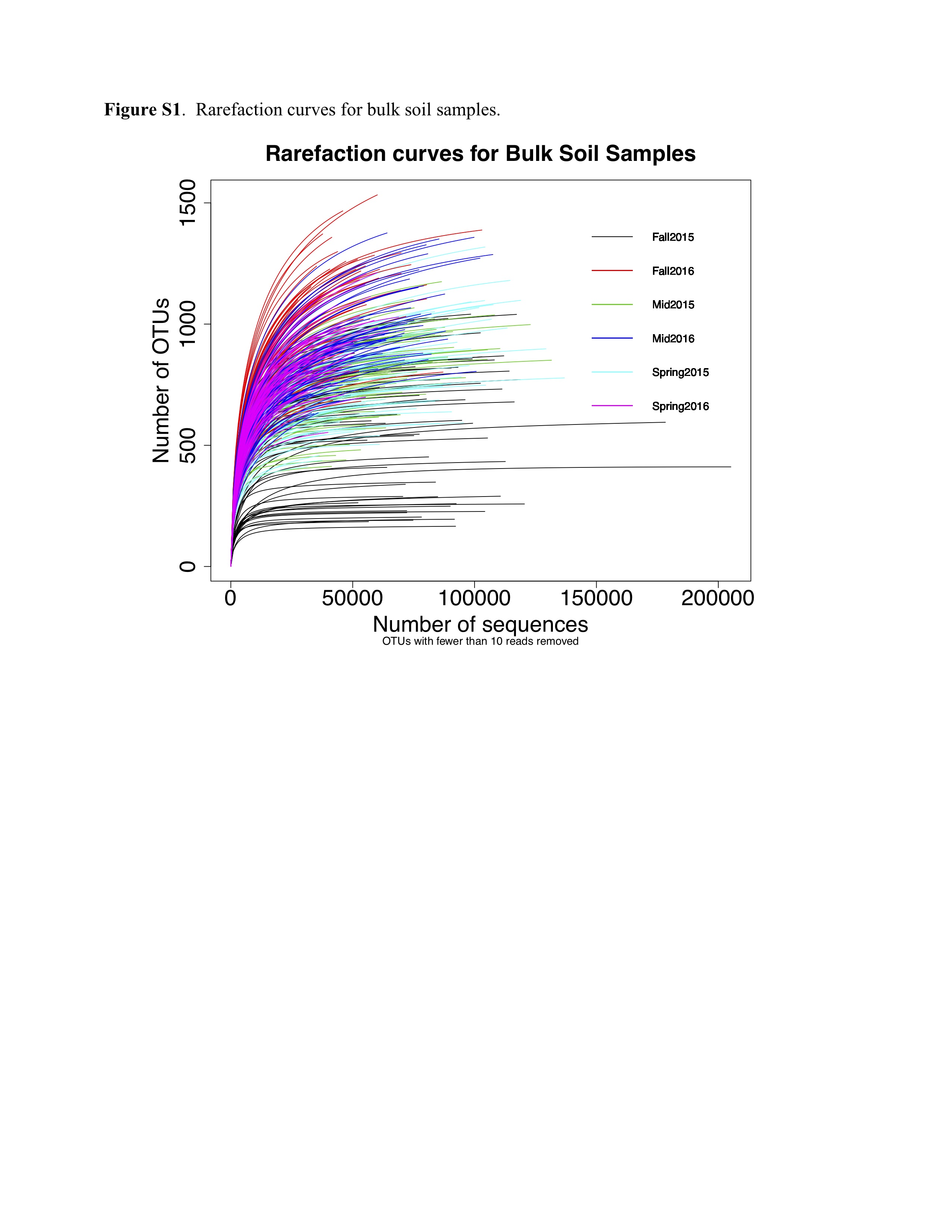

### Supplemental Figure 2

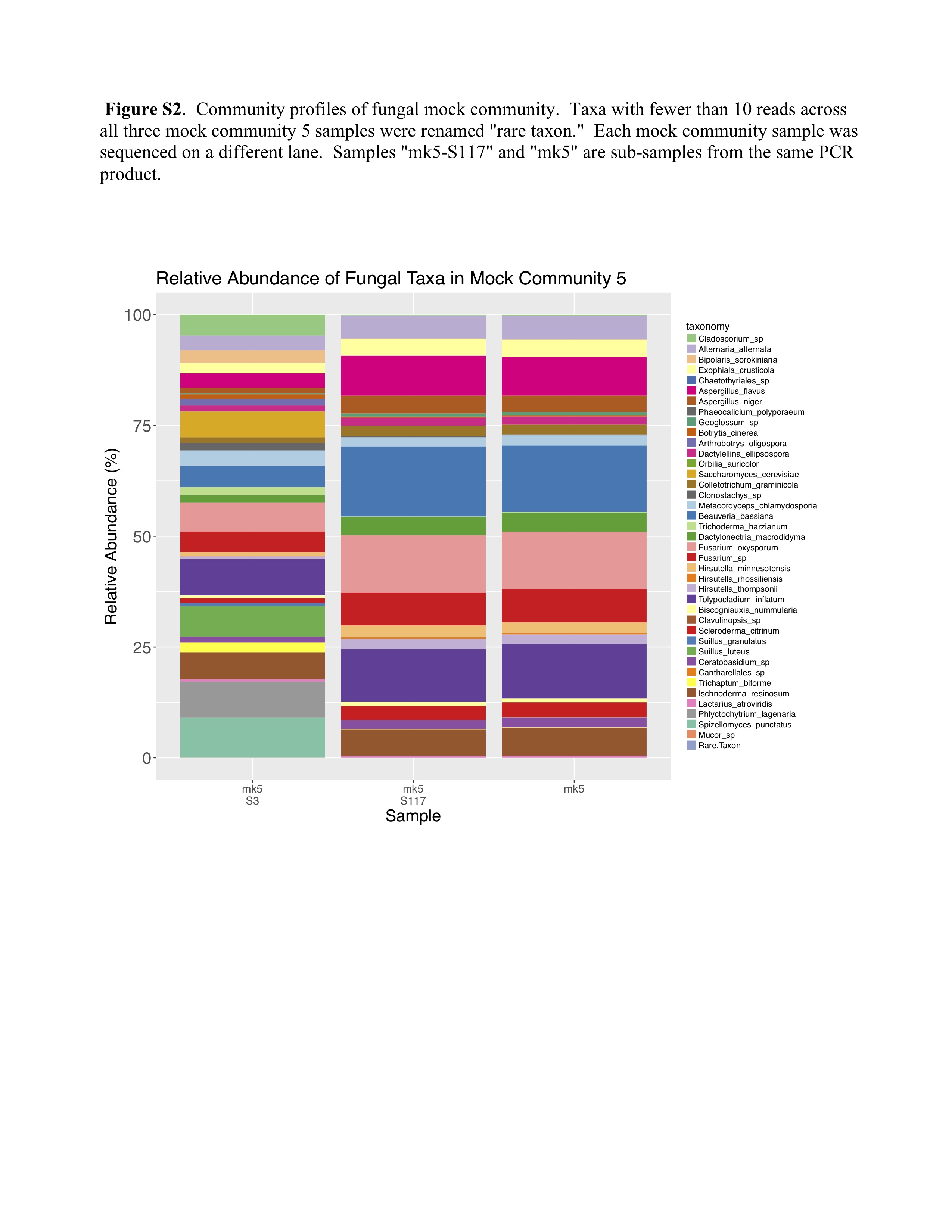

### Supplemental Figure 3

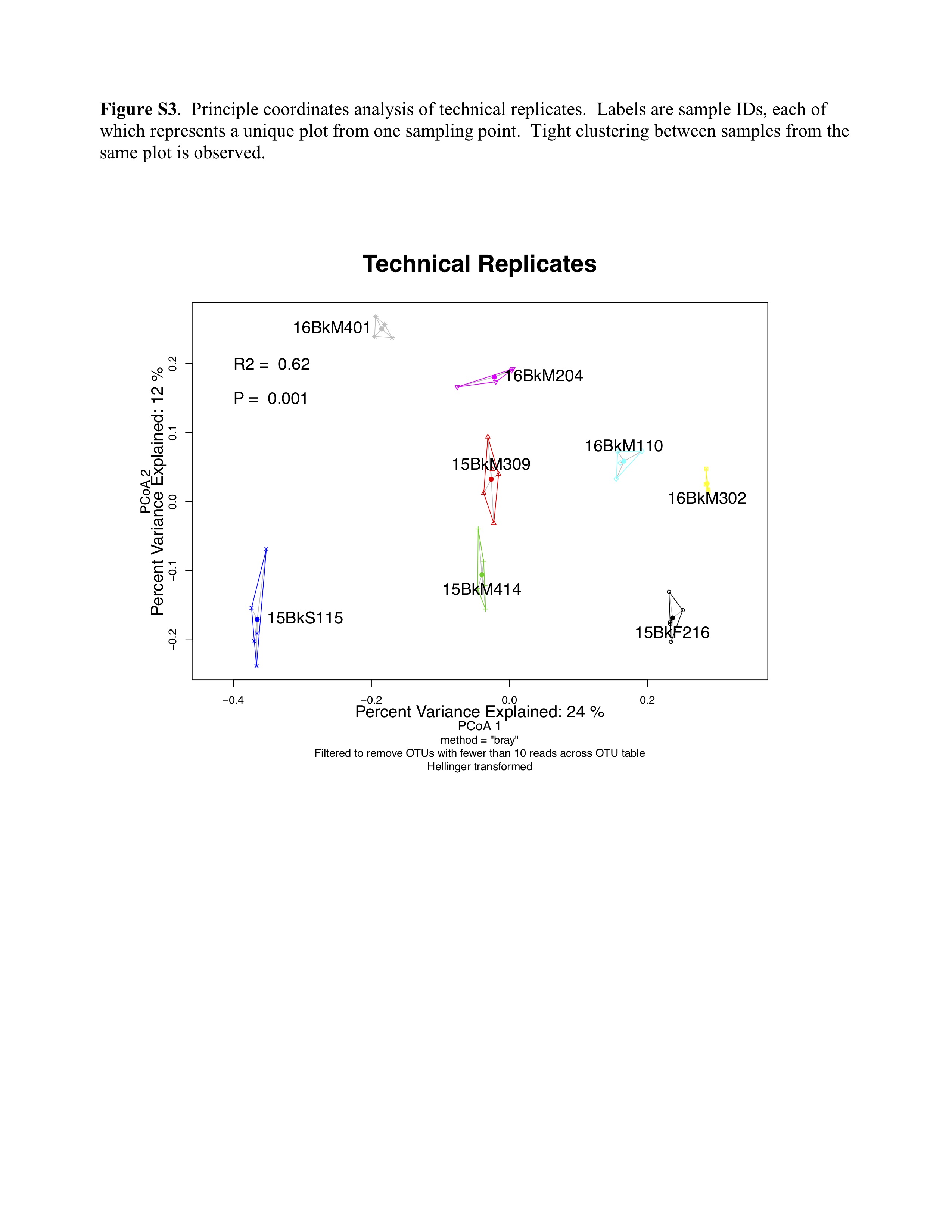

### Supplemental Figure 4

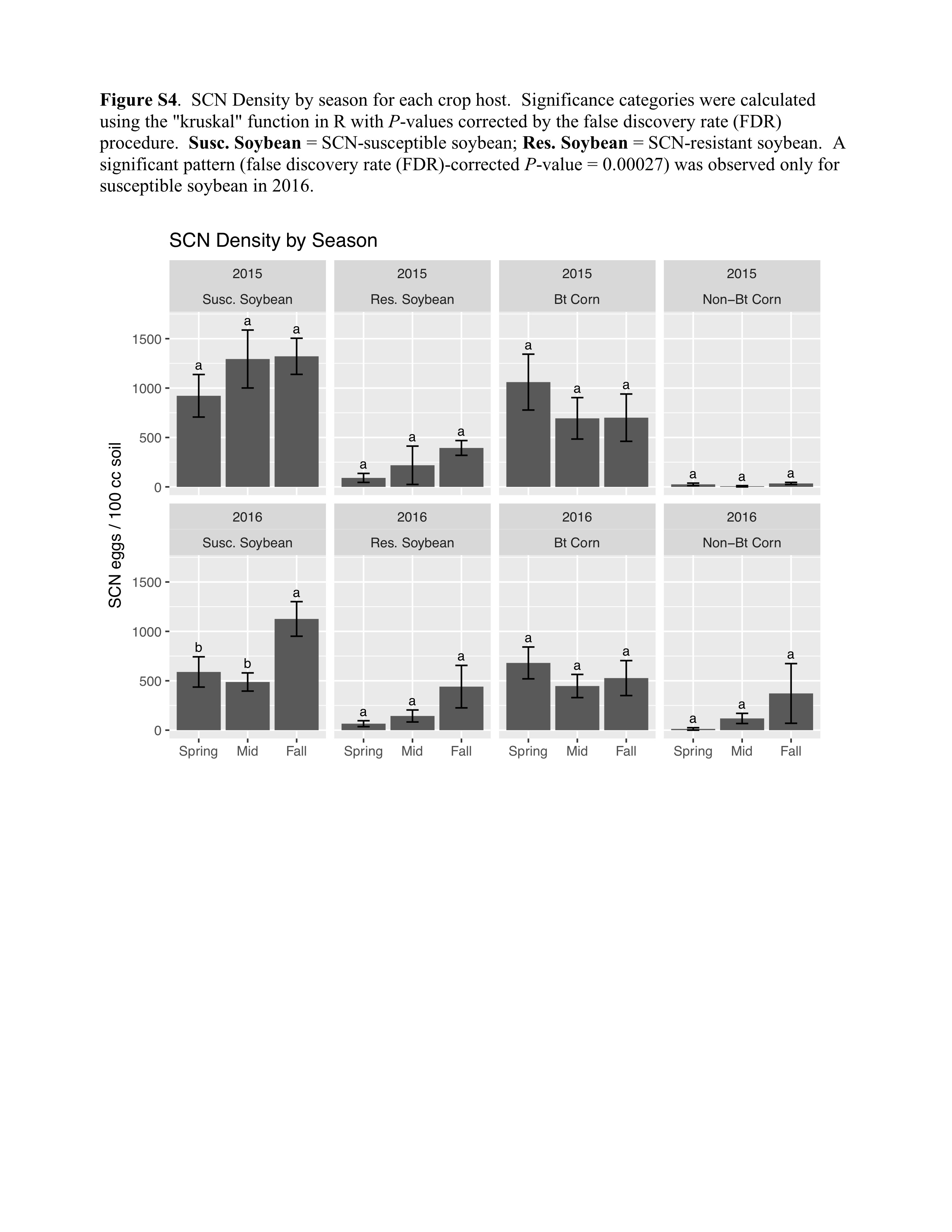

### Supplemental Figure 5

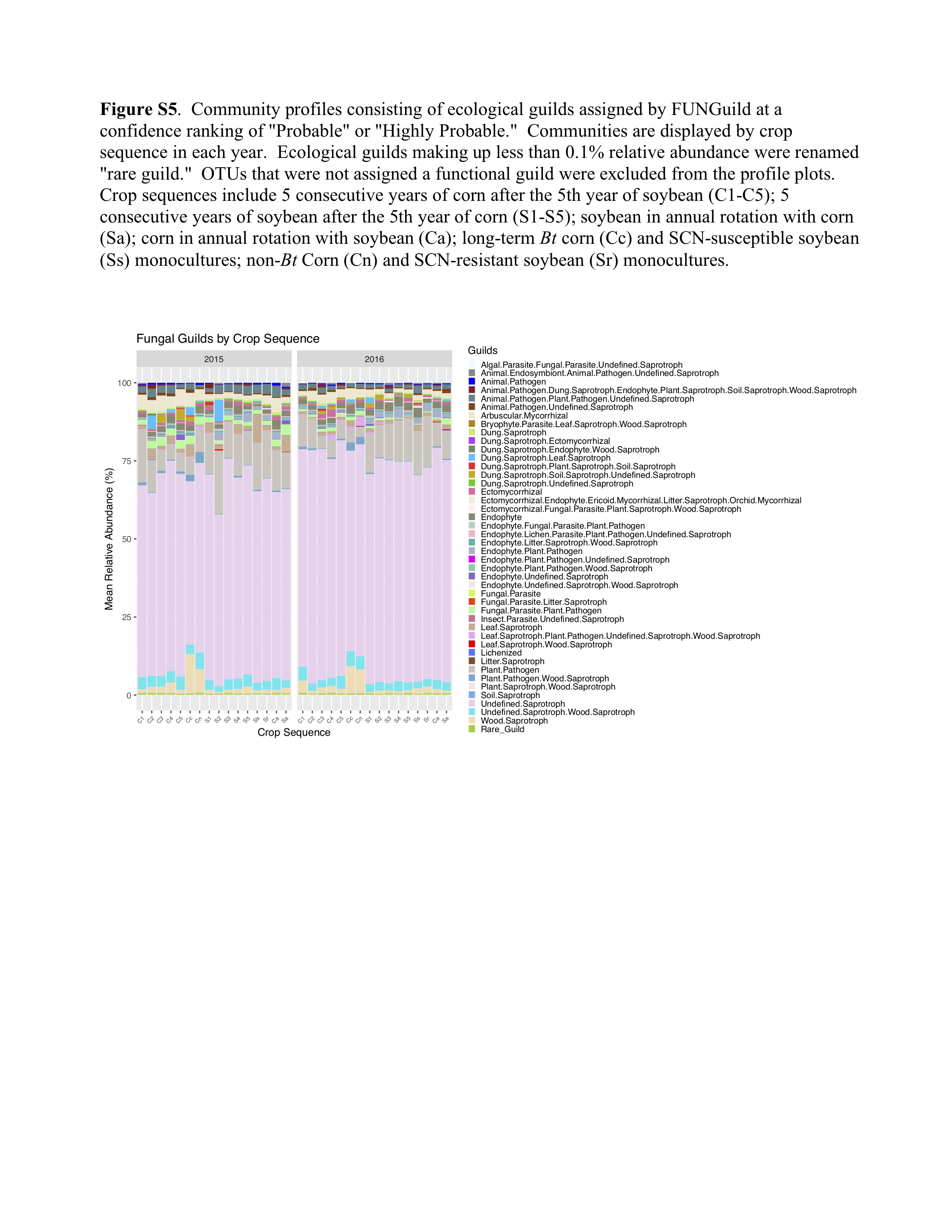

### Supplemental Figure 6

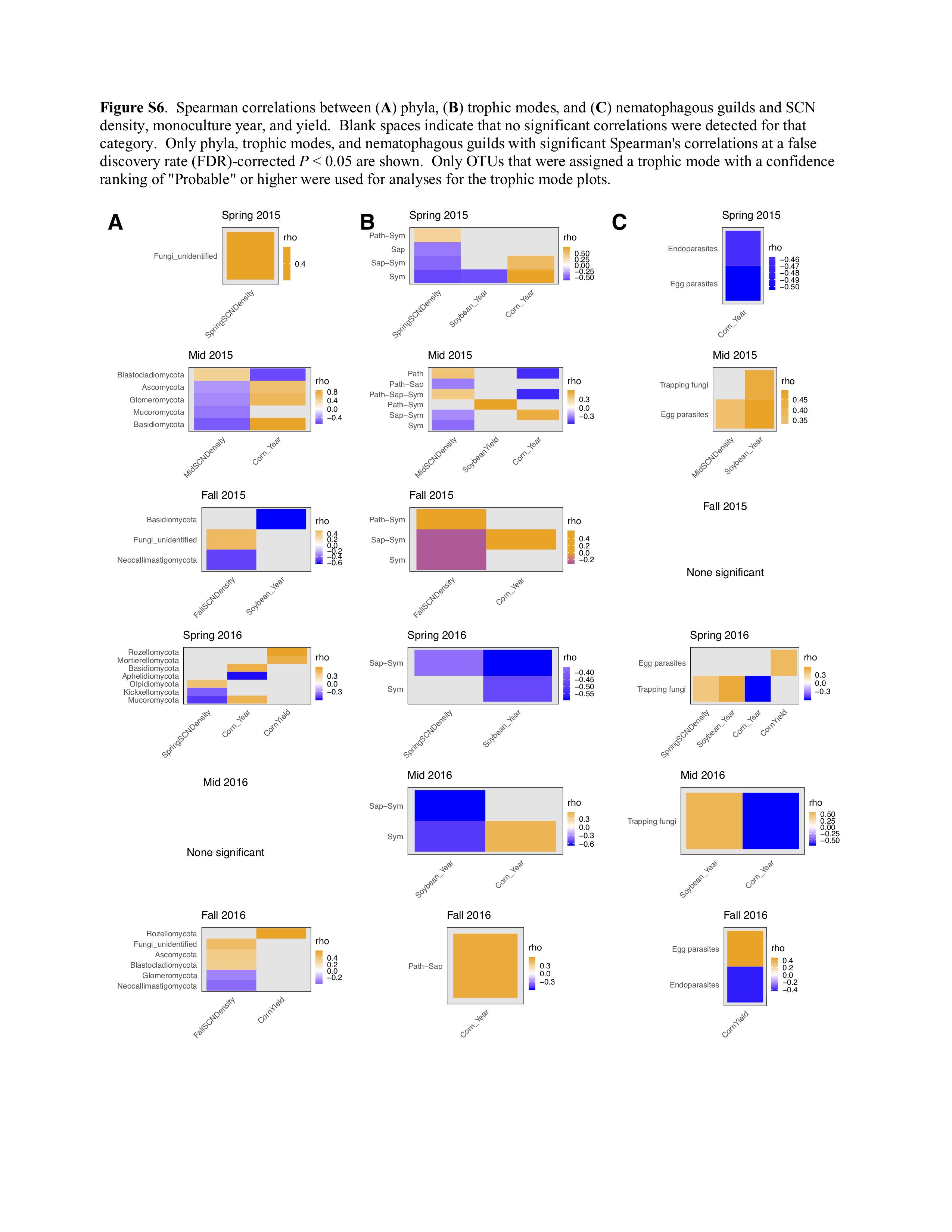

### Supplemental Figure 7

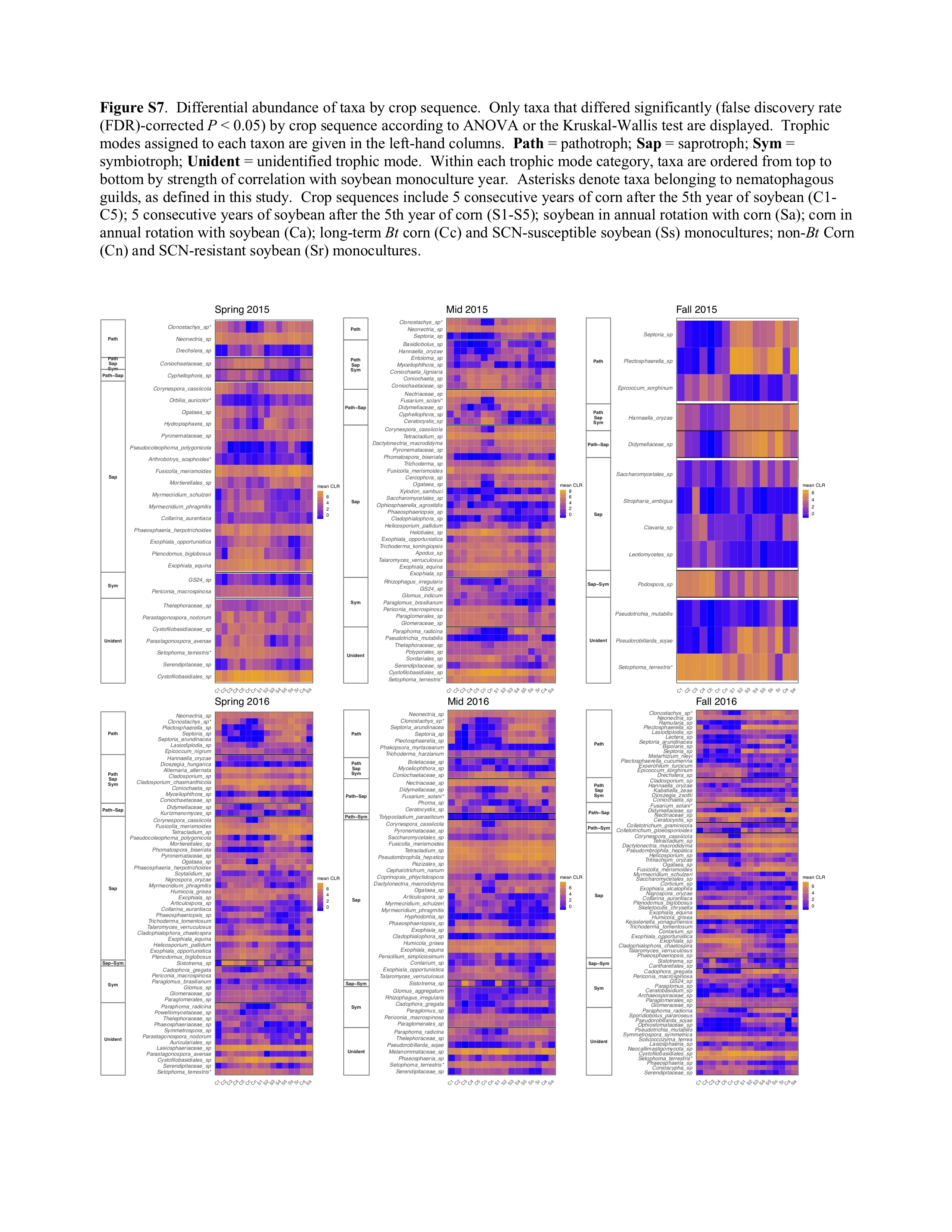

### Supplemental Figure 8

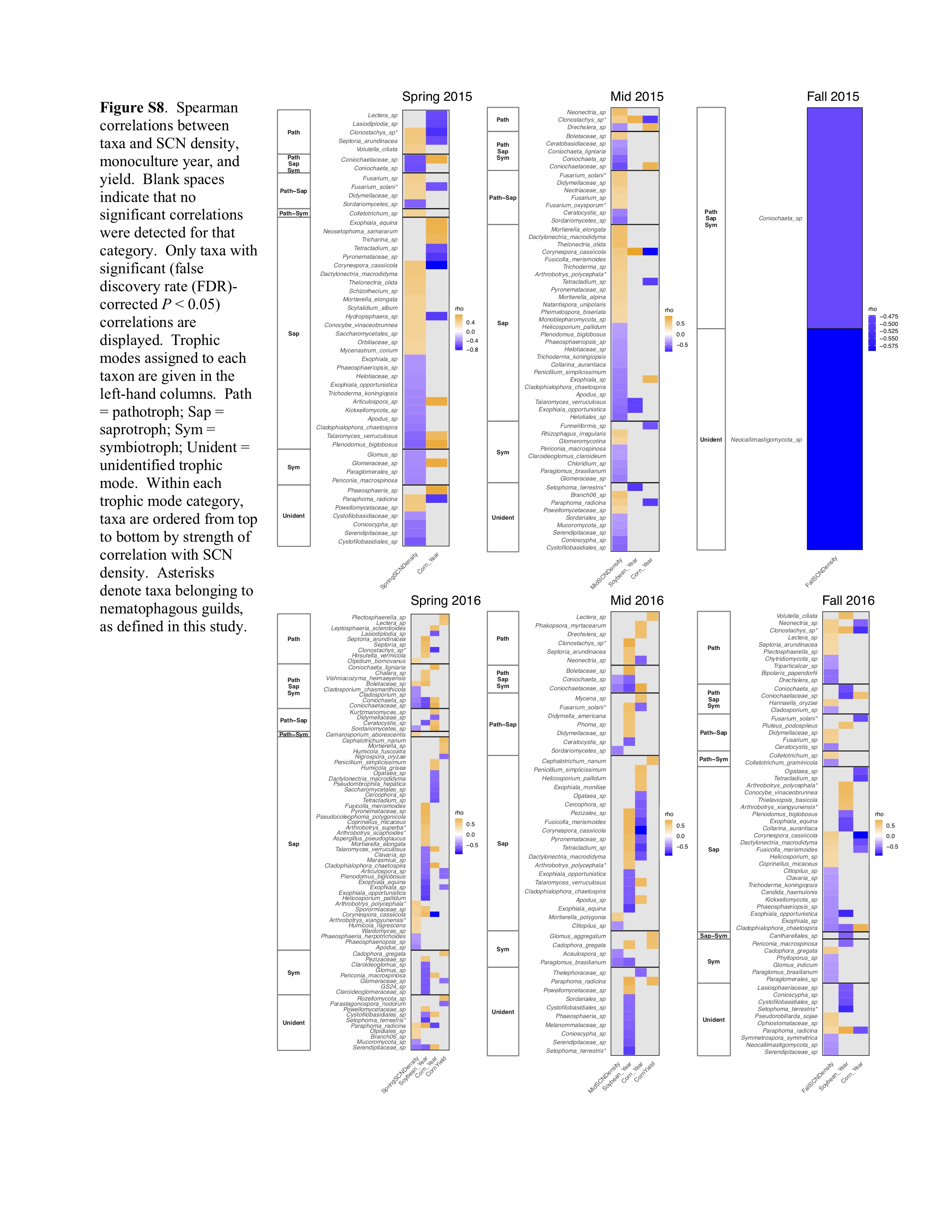

### Supplemental Figure 9

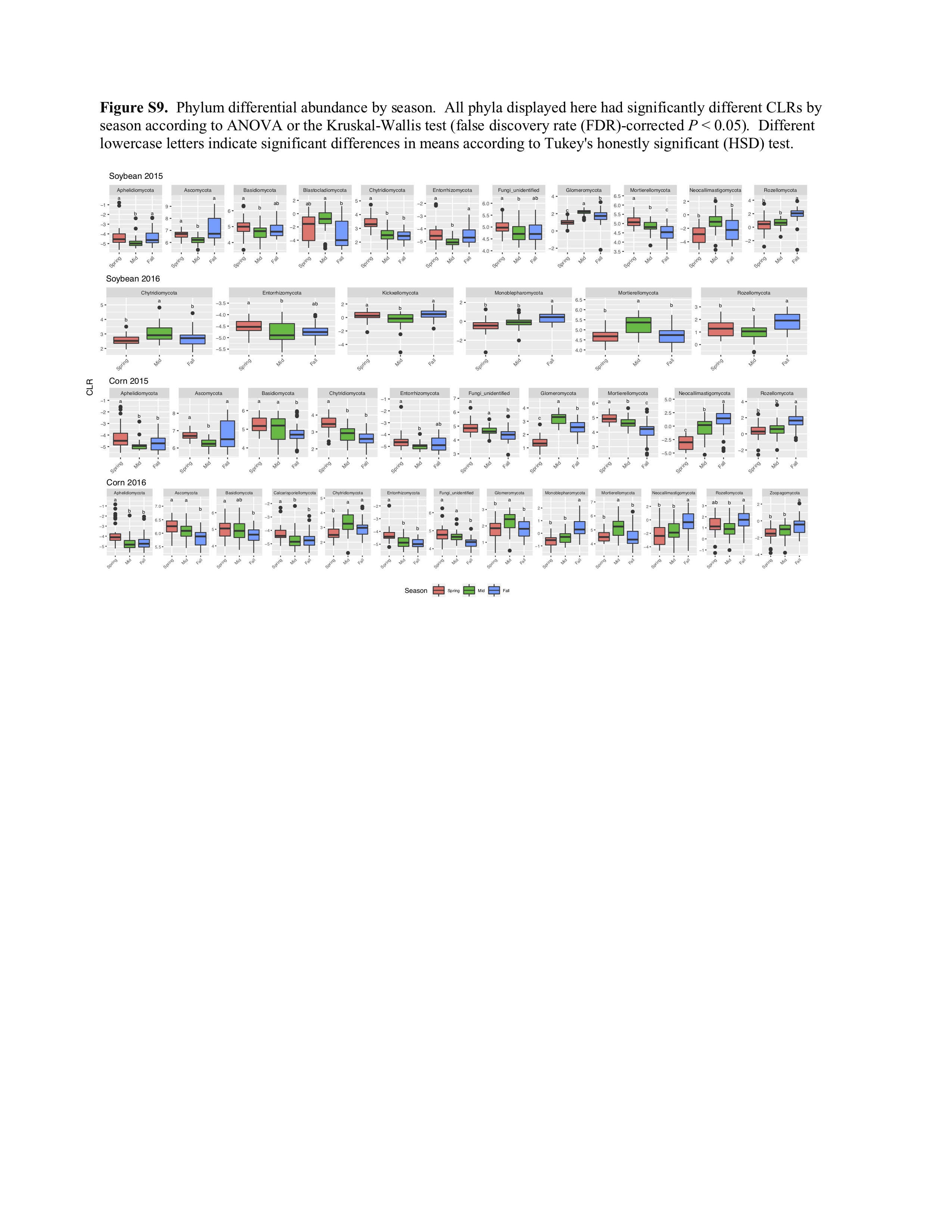

### Supplemental Figure 10

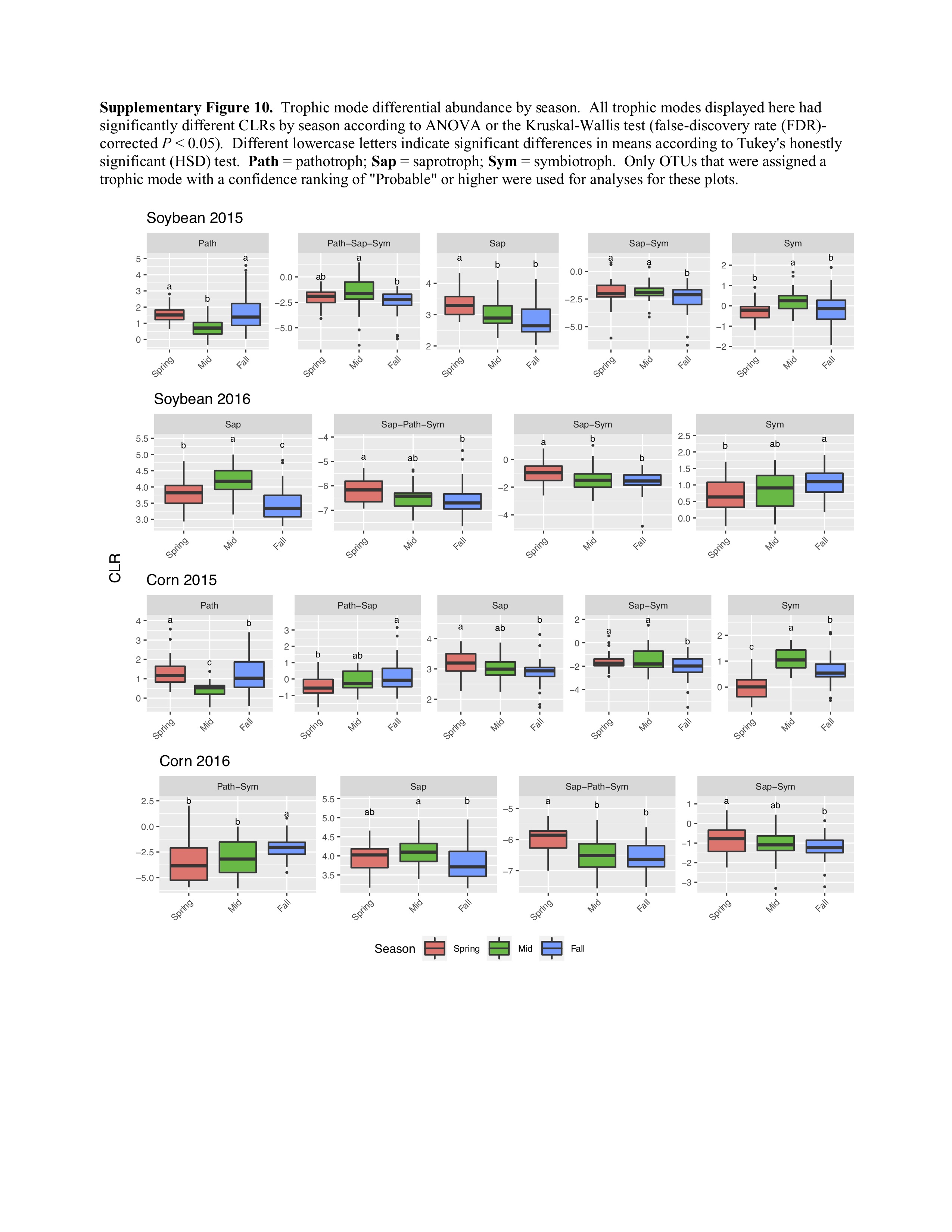

### Supplemental Figure 11

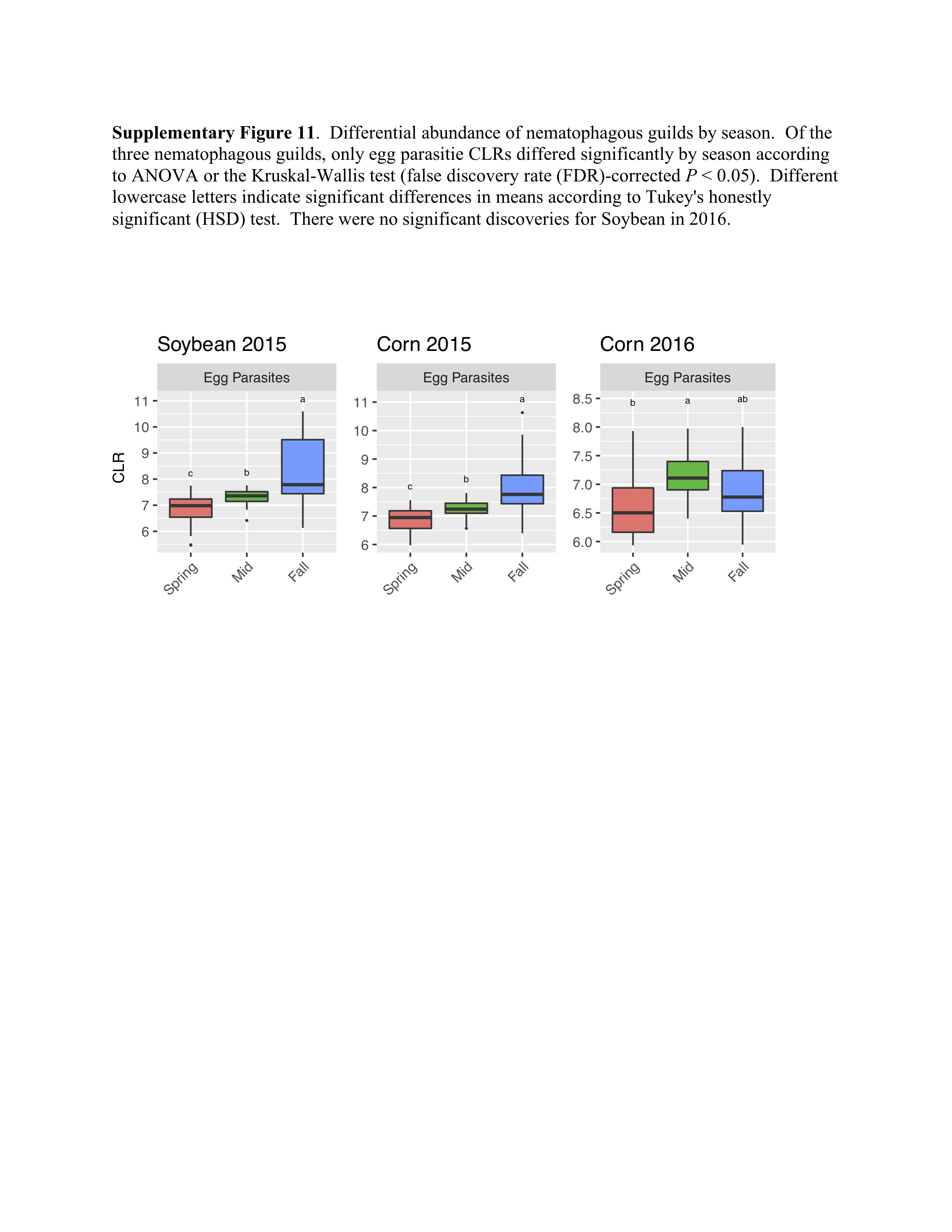
