## Supplemental Code for "Fungal Communities in Soybean Cyst Nematode-Infested Soil under Long Term Corn and Soybean Monoculture and Crop Rotation": find.longest.taxonomy.R

```
##Compare taxonomy assignment from amptk with taxonomy assignment from consensus Blast, and keep the more precise / deepest level of taxonomy, e.g. species-level taxonomy of all classifications.

##Set working directory and import table with taxonomy assigned to OTUs by two or more methods of taxonomy assignment.  Order of columns should be in order of "preferred" taxonomy assignment method.
setwd("~/Dropbox/Noah_USDA_FungiSCN/MetagenomicSequencing/Bulk.Soil.2015.2016/AMPTK/OTU_Tables/3.30.18/")

taxonomy <- read.table("Taxonomy.Assignments.3.30.2018.txt", comment="", header=TRUE, sep="\t", as.is=TRUE, check.names=F) 


#view imported taxonomy dataframe.
head(taxonomy)


##Replace all other taxonomy assignment methods with SynMock taxonomy assignment if Synthetic Mock Community sequences were assigned in AMPTK.  (I added these sequences to the database "on the fly.")
taxonomy[258,'AMPTK'] #View an example of a Synmock assignment

for(i in 1:nrow(taxonomy)){
  if(grepl('|SYN', taxonomy$AMPTK[i], fixed = TRUE)){
    synmock <- strsplit(taxonomy$AMPTK[i], '|', fixed = TRUE)[[1]][3]
    synmock <- strsplit(synmock, ';')[[1]][1]
    taxonomy$Consensus.20[i] <- paste0('k__', synmock, ';p__', synmock, ';c__', synmock, ';o__', synmock, ';f__', synmock, ';g__', synmock, ';s__', synmock)
    taxonomy$Consensus.10[i] <- paste0('k__', synmock, ';p__', synmock, ';c__', synmock, ';o__', synmock, ';f__', synmock, ';g__', synmock, ';s__', synmock)
    taxonomy$Consensus.1[i] <- paste0('k__', synmock, ';p__', synmock, ';c__', synmock, ';o__', synmock, ';f__', synmock, ';g__', synmock, ';s__', synmock)
    taxonomy$AMPTK[i] <- paste0('k__', synmock, ';p__', synmock, ';c__', synmock, ';o__', synmock, ';f__', synmock, ';g__', synmock, ';s__', synmock)
  }

}
taxonomy[258,] #View an example of a Synmock assignment


##Reformat table

##Get rid of Qiime information that I do not need.
taxonomy <- gsub("Unassigned", "", as.matrix(taxonomy))  
head(taxonomy)

taxonomy <- gsub("..__unidentified*", "", as.matrix(taxonomy))
head(taxonomy)

#Eliminate species information if we only know a higher level of taxonomy
taxonomy <- gsub(";s__.*._sp", "", as.matrix(taxonomy)) 
head(taxonomy)


##Reformat AMPTK taxonomy to match Qiime.
head(taxonomy[,'AMPTK'])
taxonomy <- gsub(" ", "_", as.matrix(taxonomy))   #put an underscore in place of a space to match Qiime genus_species format
head(taxonomy[,'AMPTK'])

taxonomy <- gsub(".*;k:", "k__", as.matrix(taxonomy))
head(taxonomy)

taxonomy <- gsub(",p:", ";p__", as.matrix(taxonomy))
head(taxonomy)

taxonomy <- gsub(",c:", ";c__", as.matrix(taxonomy))
head(taxonomy)

taxonomy <- gsub(",o:", ";o__", as.matrix(taxonomy))
head(taxonomy)

taxonomy <- gsub(",f:", ";f__", as.matrix(taxonomy))
head(taxonomy)

taxonomy <- gsub(",g:", ";g__", as.matrix(taxonomy))
head(taxonomy)

taxonomy <- gsub(",s:", ";s__", as.matrix(taxonomy))
head(taxonomy)

##Create a vector to store the most precise taxonomy.
longest.taxonomy <- vector()

##Fill vector with most complete taxonomies, beginning with those that contain species-level taxonomy, biased in order of methods I trust the most  (e.g. Consesus 20, Consesus 10, AMPTK).

labels(taxonomy)  #show taxonomy assignment methods
number.of.methods <- ncol(taxonomy)  #number of taxonomy assignment methods (+ 1 -- The first row is probably "OTU ID").  
OTU_IDs <- taxonomy[,1]  #Save OTU_IDs for later.


##First, solve conflicts between taxonomy assignment methods, trusting assignments in left-most columns over right-most columns.  (I skipped this part for Endophyte taxonomy assignments.)

#Solve conflicts between species.
for(i in 1:nrow(taxonomy)){
  split.string <- list()
  split.string <- strsplit(taxonomy[i,], ';')  
  for(k in 2:(number.of.methods-1)){  #start with k=2 to avoid the OTU_ID column.  End with second to last column.  (compare with column to the right of this one.)
    for(j in k:(number.of.methods-1)){ #k begins with k=j then moves up with j
      if(length(split.string[[k]]) >=7 & length(split.string[[j+1]]) >= 7){ #split strings must be at least length 7 to include species
        if(split.string[[k]][7] != split.string[[j+1]][7]){ #6 refers to the genus. 7 refers to the species
          split.string[[j+1]][7] <- 'N' #replace species names of taxonomy assignment methods to the right with 'N' if there is disagreement with taxonomy assignment methods to the left (left-most assignment methods are assumed to be more accurate.)
          taxonomy[i,j+1] <- paste(split.string[[j+1]], collapse = ';')  #replace in taxonomy vector
        }
      }
    }
    
  }
}


#Solve conflicts between genera.
for(i in 1:nrow(taxonomy)){
  split.string <- list()
  split.string <- strsplit(taxonomy[i,], ';')  
  for(k in 2:(number.of.methods-1)){  #start with k=2 to avoid the OTU_ID column.  End with second to last column.  (compare with column to the right of this one.)
    for(j in k:(number.of.methods-1)){ #k begins with k=j then moves up with j
      if(length(split.string[[k]]) >=6 & length(split.string[[j+1]]) >= 6){ #split strings must be at least length 6 to include genus
        if(split.string[[k]][6] != split.string[[j+1]][6]){ #6 refers to the genus. 7 refers to the species
        split.string[[j+1]][6:7] <- 'N' #replace genus and species of taxonomy assignment methods to the right with 'N' if there is disagreement with taxonomy assignment methods to the left (left-most assignment methods are assumed to be more accurate.)
        taxonomy[i,j+1] <- paste(split.string[[j+1]], collapse = ';')  #replace in taxonomy vector
      }
    }
    }
  
  }
}
  


#Solve conflicts between families.
for(i in 1:nrow(taxonomy)){
  split.string <- list()
  split.string <- strsplit(taxonomy[i,], ';')  
  for(k in 2:(number.of.methods-1)){  #start with k=2 to avoid the OTU_ID column.  End with second to last column.  (compare with column to the right of this one.)
    for(j in k:(number.of.methods-1)){ #k begins with k=j then moves up with j
      if(length(split.string[[k]]) >=5 & length(split.string[[j+1]]) >= 5){ #split strings must be at least length 5 to include family
        if(split.string[[k]][5] != split.string[[j+1]][5]){ #5 refers to the family. 7 refers to the species
          split.string[[j+1]][5:7] <- 'N' #replace family, genus, and species of taxonomy assignment methods to the right with 'N' if there is disagreement with taxonomy assignment methods to the left (left-most assignment methods are assumed to be more accurate.)
          taxonomy[i,j+1] <- paste(split.string[[j+1]], collapse = ';')  #replace in taxonomy vector
        }
      }
    }
    
  }
}


#Solve conflicts between orders.
for(i in 1:nrow(taxonomy)){
  split.string <- list()
  split.string <- strsplit(taxonomy[i,], ';')  
  for(k in 2:(number.of.methods-1)){  #start with k=2 to avoid the OTU_ID column.  End with second to last column.  (compare with column to the right of this one.)
    for(j in k:(number.of.methods-1)){ #k begins with k=j then moves up with j
      if(length(split.string[[k]]) >=4 & length(split.string[[j+1]]) >= 4){ #split strings must be at least length 5 to include family
        if(split.string[[k]][4] != split.string[[j+1]][4]){ #5 refers to the family. 7 refers to the species
          split.string[[j+1]][4:7] <- 'N' #replace order, family, genus, and species of taxonomy assignment methods to the right with 'N' if there is disagreement with taxonomy assignment methods to the left (left-most assignment methods are assumed to be more accurate.)
          taxonomy[i,j+1] <- paste(split.string[[j+1]], collapse = ';')  #replace in taxonomy vector
        }
      }
    }
    
  }
}


#Solve conflicts between classes.
for(i in 1:nrow(taxonomy)){
  split.string <- list()
  split.string <- strsplit(taxonomy[i,], ';')  
  for(k in 2:(number.of.methods-1)){  #start with k=2 to avoid the OTU_ID column.  End with second to last column.  (compare with column to the right of this one.)
    for(j in k:(number.of.methods-1)){ #k begins with k=j then moves up with j
      if(length(split.string[[k]]) >=3 & length(split.string[[j+1]]) >= 3){ #split strings must be at least length 5 to include family
        if(split.string[[k]][3] != split.string[[j+1]][3]){ #3 refers to the class. 
          split.string[[j+1]][3:7] <- 'N' #replace class, order, family, genus, and species of taxonomy assignment methods to the right with 'N' if there is disagreement with taxonomy assignment methods to the left (left-most assignment methods are assumed to be more accurate.)
          taxonomy[i,j+1] <- paste(split.string[[j+1]], collapse = ';')  #replace in taxonomy vector
        }
      }
    }
    
  }
}


#Solve conflicts between phyla.
for(i in 1:nrow(taxonomy)){
  split.string <- list()
  split.string <- strsplit(taxonomy[i,], ';')  
  for(k in 2:(number.of.methods-1)){  #start with k=2 to avoid the OTU_ID column.  End with second to last column.  (compare with column to the right of this one.)
    for(j in k:(number.of.methods-1)){ #k begins with k=j then moves up with j
      if(length(split.string[[k]]) >=2 & length(split.string[[j+1]]) >= 2){ #split strings must be at least length 2 to include phylum
        if(split.string[[k]][2] != split.string[[j+1]][2]){ #2 refers to the phylum. 
          split.string[[j+1]][2:7] <- 'N' #replace phylum, class, order, family, genus, and species of taxonomy assignment methods to the right with 'N' if there is disagreement with taxonomy assignment methods to the left (left-most assignment methods are assumed to be more accurate.)
          taxonomy[i,j+1] <- paste(split.string[[j+1]], collapse = ';')  #replace in taxonomy vector
        }
      }
    }
    
  }
}


#Solve conflicts between kingdoms.
for(i in 1:nrow(taxonomy)){
  split.string <- list()
  split.string <- strsplit(taxonomy[i,], ';')  
  for(k in 2:(number.of.methods-1)){  #start with k=2 to avoid the OTU_ID column.  End with second to last column.  (compare with column to the right of this one.)
    for(j in k:(number.of.methods-1)){ #k begins with k=j then moves up with j
      if(length(split.string[[k]]) >=1 & length(split.string[[j+1]]) >= 1){ #split strings must be at least length 1 to include kingdom
        if(split.string[[k]][1] != split.string[[j+1]][1]){ #2 refers to the phylum. 
          split.string[[j+1]][1:7] <- 'N' #replace phylum, class, order, family, genus, and species of taxonomy assignment methods to the right with 'N' if there is disagreement with taxonomy assignment methods to the left (left-most assignment methods are assumed to be more accurate.)
          taxonomy[i,j+1] <- paste(split.string[[j+1]], collapse = ';')  #replace in taxonomy vector
        }
      }
    }
    
  }
}


##Fill vector with most complete taxonomies, beginning with those that contain species-level taxonomy, biased in order of methods I trust the most  (e.g. Consesus 20, Consesus 10, AMPTK).

##Species-level taxonomy

for(j in 2:number.of.methods){
  for(i in 1:nrow(taxonomy)){
    #Retain species taxonomy, if best method has a species.
    if(grepl("s__", taxonomy[i,j])){
      longest.taxonomy[i] <- taxonomy[i,j]
      
      #Change taxonomy from other methods to "N"
 
      if(j == 2){
        taxonomy[i,(j+1):number.of.methods] <- "N"
      }
      else{
      
      if(j < number.of.methods){
      taxonomy[i,(j+1):number.of.methods] <- "N"   #Change taxonomy of methods to right of current method to "N"
      taxonomy[i,2:(j-1)] <- "N"    #Change taxonomy of methods to left of current method to "N"
      }
      else{
        taxonomy[i,2:(j-1)] <- "N"  #Change taxonomy of methods to left of rightmost method to "N"
      }
      }
    }
  }
}

head(longest.taxonomy)
head(taxonomy)


##Genus-level taxonomy

for(j in 2:number.of.methods){
  for(i in 1:nrow(taxonomy)){
    #Retain genus taxonomy, if best method has a genus.
    if(grepl("g__", taxonomy[i,j])){
      longest.taxonomy[i] <- taxonomy[i,j]
      
      #Change taxonomy from other methods to "N"
 
      if(j == 2){
        taxonomy[i,(j+1):number.of.methods] <- "N"
      }
      else{
           
      if(j < number.of.methods){
        taxonomy[i,(j+1):number.of.methods] <- "N"   #Change taxonomy of methods to right of current method to "N"
        taxonomy[i,2:(j-1)] <- "N"    #Change taxonomy of methods to left of current method to "N"
      }
      else{
        taxonomy[i,2:(j-1)] <- "N"  #Change taxonomy of methods to left of rightmost method to "N"
      }
      }
    }
  }
}

head(longest.taxonomy)
head(taxonomy)
tail(longest.taxonomy)
tail(taxonomy)


##Family-level taxonomy

for(j in 2:number.of.methods){
  for(i in 1:nrow(taxonomy)){
    #Retain family taxonomy, if best method has a family.
    if(grepl("f__", taxonomy[i,j])){
      longest.taxonomy[i] <- taxonomy[i,j]
      
      #Change taxonomy from other methods to "N"
      
      if(j == 2){
        taxonomy[i,(j+1):number.of.methods] <- "N"
      }
      else{
      
      if(j < number.of.methods){
        taxonomy[i,(j+1):number.of.methods] <- "N"   #Change taxonomy of methods to right of current method to "N"
        taxonomy[i,2:(j-1)] <- "N"    #Change taxonomy of methods to left of current method to "N"
      }
      else{
        taxonomy[i,2:(j-1)] <- "N"  #Change taxonomy of methods to left of rightmost method to "N"
      }
      }
    }
  }
}

head(longest.taxonomy)
tail(longest.taxonomy)
head(taxonomy)


##Order-level taxonomy

for(j in 2:number.of.methods){
  for(i in 1:nrow(taxonomy)){
    #Retain order taxonomy, if best method has an order.
    if(grepl("o__", taxonomy[i,j])){
      longest.taxonomy[i] <- taxonomy[i,j]
      
      #Change taxonomy from other methods to "N"
      
      if(j == 2){
        taxonomy[i,(j+1):number.of.methods] <- "N"
      }
      else{
      
      if(j < number.of.methods){
        taxonomy[i,(j+1):number.of.methods] <- "N"   #Change taxonomy of methods to right of current method to "N"
        taxonomy[i,2:(j-1)] <- "N"    #Change taxonomy of methods to left of current method to "N"
      }
      else{
        taxonomy[i,2:(j-1)] <- "N"  #Change taxonomy of methods to left of rightmost method to "N"
      }
      }
    }
  }
}

tail(longest.taxonomy)
tail(taxonomy)


##Class-level taxonomy

for(j in 2:number.of.methods){
  for(i in 1:nrow(taxonomy)){
    #Retain class taxonomy, if best method has a class.
    if(grepl("c__", taxonomy[i,j])){
      longest.taxonomy[i] <- taxonomy[i,j]
      
      #Change taxonomy from other methods to "N"
      
      if(j == 2){
        taxonomy[i,(j+1):number.of.methods] <- "N"
      }
      else{
        
      if(j < number.of.methods){
        taxonomy[i,(j+1):number.of.methods] <- "N"   #Change taxonomy of methods to right of current method to "N"
        taxonomy[i,2:(j-1)] <- "N"    #Change taxonomy of methods to left of current method to "N"
      }
      else{
        taxonomy[i,2:(j-1)] <- "N"  #Change taxonomy of methods to left of rightmost method to "N"
      }
      }
    }
  }
}

tail(longest.taxonomy)
longest.taxonomy[59]
longest.taxonomy[61]
tail(taxonomy)


##Phylum-level taxonomy

for(j in 2:number.of.methods){
  for(i in 1:nrow(taxonomy)){
    #Retain phylum taxonomy, if best method has a phylum.
    if(grepl("p__", taxonomy[i,j])){
      longest.taxonomy[i] <- taxonomy[i,j]
      
      #Change taxonomy from other methods to "N"
      
      if(j == 2){
        taxonomy[i,(j+1):number.of.methods] <- "N"
      }
      else{
        
      if(j < number.of.methods){
        taxonomy[i,(j+1):number.of.methods] <- "N"   #Change taxonomy of methods to right of current method to "N"
        taxonomy[i,2:(j-1)] <- "N"    #Change taxonomy of methods to left of current method to "N"
      }
      else{
        taxonomy[i,2:(j-1)] <- "N"  #Change taxonomy of methods to left of rightmost method to "N"
      }
      }
    }
  }
}

tail(longest.taxonomy)
tail(taxonomy)


##Kingdom-level taxonomy

for(j in 2:number.of.methods){
  for(i in 1:nrow(taxonomy)){
    #Retain kingdom taxonomy, if best method has a kingdom.
    if(grepl("k__", taxonomy[i,j])){
      longest.taxonomy[i] <- taxonomy[i,j]
      
      #Change taxonomy from other methods to "N"
      
      if(j == 2){
        taxonomy[i,(j+1):number.of.methods] <- "N"
      }
      else{
        
      if(j < number.of.methods){
        taxonomy[i,(j+1):number.of.methods] <- "N"   #Change taxonomy of methods to right of current method to "N"
        taxonomy[i,2:(j-1)] <- "N"    #Change taxonomy of methods to left of current method to "N"
      }
      else{
        taxonomy[i,2:(j-1)] <- "N"  #Change taxonomy of methods to left of rightmost method to "N"
      }
      }
    }
  }
}

tail(longest.taxonomy)
tail(taxonomy)

##Add column to taxonomy Excel sheet.
taxonomy <- as.data.frame(taxonomy)
taxonomy$longest.taxonomy <- longest.taxonomy
taxonomy$`#OTU ID` <- OTU_IDs
#row.names(taxonomy) <- OTU_IDs
head(taxonomy)

head(taxonomy$longest.taxonomy) #View longest taxonomy vector

##Reformat longest taxonomy vector to work with downstream Qiime applications.  Basically, just add "identified" where an intermediate level of taxonomy is unknown.

#make new column to hold qiime-formatted taxonomy
taxonomy$qiime.formatted.longest.taxonomy <- taxonomy$longest.taxonomy

taxonomy$qiime.formatted.longest.taxonomy <- gsub("Fungi;c__", "Fungi;p__unidentified;c__", as.matrix(taxonomy$qiime.formatted.longest.taxonomy))

taxonomy$qiime.formatted.longest.taxonomy <- gsub("Fungi;o__", "Fungi;p__unidentified;c__unidentified;o__", as.matrix(taxonomy$qiime.formatted.longest.taxonomy))

taxonomy$qiime.formatted.longest.taxonomy <- gsub("mycota;o__", "mycota;c__unidentified;o__", as.matrix(taxonomy$qiime.formatted.longest.taxonomy))

taxonomy$qiime.formatted.longest.taxonomy <- gsub("mycota;f__", "mycota;c__unidentified;o__unidentified;f__", as.matrix(taxonomy$qiime.formatted.longest.taxonomy))

taxonomy$qiime.formatted.longest.taxonomy <- gsub("mycota;g__", "mycota;c__unidentified;o__unidentified;f__unidentified;g__", as.matrix(taxonomy$qiime.formatted.longest.taxonomy))

taxonomy$qiime.formatted.longest.taxonomy <- gsub("mycota;s__", "mycota;c__unidentified;o__unidentified;f__unidentified;g__unidentified;s__", as.matrix(taxonomy$qiime.formatted.longest.taxonomy))

taxonomy$qiime.formatted.longest.taxonomy <- gsub("mycetes;f__", "mycetes;o__unidentified;f__", as.matrix(taxonomy$qiime.formatted.longest.taxonomy))

taxonomy$qiime.formatted.longest.taxonomy <- gsub("mycetes;g__", "mycetes;o__unidentified;f__unidentified;g__", as.matrix(taxonomy$qiime.formatted.longest.taxonomy))

taxonomy$qiime.formatted.longest.taxonomy <- gsub("mycetes;s__", "mycetes;o__unidentified;f__unidentified;g__unidentified;s__", as.matrix(taxonomy$qiime.formatted.longest.taxonomy))

taxonomy$qiime.formatted.longest.taxonomy <- gsub("ales;g__", "ales;f__unidentified;g__", as.matrix(taxonomy$qiime.formatted.longest.taxonomy))

taxonomy$qiime.formatted.longest.taxonomy <- gsub("ales;s__", "ales;f__unidentified;g__unidentified;s__", as.matrix(taxonomy$qiime.formatted.longest.taxonomy))

taxonomy$qiime.formatted.longest.taxonomy <- gsub("eae;s__", "eae;g__unidentified;s__", as.matrix(taxonomy$qiime.formatted.longest.taxonomy))

head(taxonomy)

#Alternatively, try replacing missing levels with "incertae sedis."  (This is more complicated, and may not be worth my time.)


#Fix jumps from class ("mycetes") to family (";f__")
for(i in 1:length(taxonomy$longest.taxonomy)){
  if(grepl('mycetes;f__', taxonomy$longest.taxonomy[i])){
    split.taxon <- strsplit(taxonomy$longest.taxonomy[i], ';')[[1]][3]
    taxon.name <- strsplit(split.taxon, '__')[[1]][2]
    replacement.taxon.name.with.incertae.sedis <- paste0('mycetes;o__', taxon.name, '_ord_Incertae_sedis;f__')
    split.old.name <- strsplit(taxonomy$longest.taxonomy[i], 'mycetes;f__')
    new.name <- paste(split.old.name[[1]][1], replacement.taxon.name.with.incertae.sedis, split.old.name[[1]][2], sep = "", collapse = "")
    new.name
    taxonomy$qiime.formatted.longest.taxonomy[i] <- new.name
  }
}

taxonomy$longest.taxonomy[8659]
taxonomy$qiime.formatted.longest.taxonomy[8659]

#Fix jumps from class ("mycetes") to genus (";g__")
for(i in 1:length(taxonomy$longest.taxonomy)){
  if(grepl('mycetes;g__', taxonomy$longest.taxonomy[i])){
    split.taxon <- strsplit(taxonomy$longest.taxonomy[i], ';')[[1]][3]
    taxon.name <- strsplit(split.taxon, '__')[[1]][2]
    replacement.taxon.name.with.incertae.sedis <- paste0('mycetes;o__', taxon.name, '_ord_Incertae_sedis;f__', taxon.name, '_fam_Incertae_sedis;')
    split.old.name <- strsplit(taxonomy$longest.taxonomy[i], 'mycetes;g__')
    new.name <- paste(split.old.name[[1]][1], replacement.taxon.name.with.incertae.sedis, 'g__', split.old.name[[1]][2], sep = "", collapse = "")
    new.name
    taxonomy$qiime.formatted.longest.taxonomy[i] <- new.name
  }
}

taxonomy$longest.taxonomy[11342]
taxonomy$qiime.formatted.longest.taxonomy[11342]

#Fix jumps from family to genus
for(i in 1:length(taxonomy$longest.taxonomy)){
  if(grepl('ales;g_', taxonomy$longest.taxonomy[i])){
    split.taxon <- strsplit(taxonomy$longest.taxonomy[i], ';')[[1]][4]
    taxon.name <- strsplit(split.taxon, '__')[[1]][2]
    replacement.taxon.name.with.incertae.sedis <- paste0('ales;f__', taxon.name, '_fam_Incertae_sedis;g__')
    split.old.name <- strsplit(taxonomy$longest.taxonomy[i], 'ales;g__')
    new.name <- paste(split.old.name[[1]][1], replacement.taxon.name.with.incertae.sedis, split.old.name[[1]][2], sep = "", collapse = "")
    new.name
    taxonomy$qiime.formatted.longest.taxonomy[i] <- new.name
  }
}

taxonomy$longest.taxonomy[18]
taxonomy$qiime.formatted.longest.taxonomy[18]


#Optional:  write the taxonomy file, here.  Or, proceed to modify the taxonomy so it will work in Qiime2
#write.csv(taxonomy, file = 'taxonomy.with.longest.taxonomy.csv', row.names = TRUE)


###Modify taxonomy so it works in QIIME!


#view deficient taxon.
taxonomy$qiime.formatted.longest.taxonomy[20]

#add phylum
for(i in 1: length(taxonomy$qiime.formatted.longest.taxonomy)){

  if(grepl("p__", taxonomy$qiime.formatted.longest.taxonomy[i]) == 'FALSE'){
    
    taxonomy$qiime.formatted.longest.taxonomy[i] <- paste(taxonomy$qiime.formatted.longest.taxonomy[i], ";p__unidentified", sep = "")

    }
}

#check work
taxonomy$qiime.formatted.longest.taxonomy[20]

#add class
for(i in 1: length(taxonomy$qiime.formatted.longest.taxonomy)){
  
  if(grepl("c__", taxonomy$qiime.formatted.longest.taxonomy[i]) == 'FALSE'){
    
    taxonomy$qiime.formatted.longest.taxonomy[i] <- paste(taxonomy$qiime.formatted.longest.taxonomy[i], ";c__unidentified", sep = "")
    
  }
}

#check work
taxonomy$qiime.formatted.longest.taxonomy[20]

#add order
for(i in 1: length(taxonomy$qiime.formatted.longest.taxonomy)){
  
  if(grepl("o__", taxonomy$qiime.formatted.longest.taxonomy[i]) == 'FALSE'){
    
    taxonomy$qiime.formatted.longest.taxonomy[i] <- paste(taxonomy$qiime.formatted.longest.taxonomy[i], ";o__unidentified", sep = "")
    
  }
}

#check work
taxonomy$qiime.formatted.longest.taxonomy[20]

#add family
for(i in 1: length(taxonomy$qiime.formatted.longest.taxonomy)){
  
  if(grepl("f__", taxonomy$qiime.formatted.longest.taxonomy[i]) == 'FALSE'){
    
    taxonomy$qiime.formatted.longest.taxonomy[i] <- paste(taxonomy$qiime.formatted.longest.taxonomy[i], ";f__unidentified", sep = "")
    
  }
}

#check work
taxonomy$qiime.formatted.longest.taxonomy[20]


#add genus
for(i in 1: length(taxonomy$qiime.formatted.longest.taxonomy)){
  
  if(grepl("g__", taxonomy$qiime.formatted.longest.taxonomy[i]) == 'FALSE'){
    
    taxonomy$qiime.formatted.longest.taxonomy[i] <- paste(taxonomy$qiime.formatted.longest.taxonomy[i], ";g__unidentified", sep = "")
    
  }
}

#check work
taxonomy$qiime.formatted.longest.taxonomy[20]


#add species.  Note:  This one is unique!  It adds "sp" to the lowest level of taxonomy.


taxonomy$qiime.formatted.longest.taxonomy[980]
taxonomy$qiime.formatted.longest.taxonomy[992]
taxonomy$qiime.formatted.longest.taxonomy[11]
taxonomy$qiime.formatted.longest.taxonomy[20]

#If genus is present and species is absent, use this.
for(i in 1: length(taxonomy$qiime.formatted.longest.taxonomy)){
  if(strsplit(taxonomy$qiime.formatted.longest.taxonomy[i], split = ";")[[1]][6] != "g__unidentified" & length(strsplit(taxonomy$qiime.formatted.longest.taxonomy[i], split = ";")[[1]]) == 6){
    genus <- strsplit(taxonomy$qiime.formatted.longest.taxonomy[i], split = ";")[[1]][6]
    genus <- sub(".*__", "", genus)
    taxonomy$qiime.formatted.longest.taxonomy[i] <- paste(taxonomy$qiime.formatted.longest.taxonomy[i], ";s__", genus, "_sp", sep = "")
  }
}

#check work.
taxonomy$qiime.formatted.longest.taxonomy[980]
taxonomy$qiime.formatted.longest.taxonomy[992]
taxonomy$qiime.formatted.longest.taxonomy[11]
taxonomy$qiime.formatted.longest.taxonomy[20]

taxonomy$qiime.formatted.longest.taxonomy[963]

#If family is present, use this.
for(i in 1: length(taxonomy$qiime.formatted.longest.taxonomy)){
  if(strsplit(taxonomy$qiime.formatted.longest.taxonomy[i], split = ";")[[1]][5] != "f__unidentified" & length(strsplit(taxonomy$qiime.formatted.longest.taxonomy[i], split = ";")[[1]]) == 6){
    family <- strsplit(taxonomy$qiime.formatted.longest.taxonomy[i], split = ";")[[1]][5]
    family <- sub(".*__", "", family)
    taxonomy$qiime.formatted.longest.taxonomy[i] <- paste(taxonomy$qiime.formatted.longest.taxonomy[i], ";s__", family, "_sp", sep = "")
  }
}

#check work.
taxonomy$qiime.formatted.longest.taxonomy[963]
taxonomy$qiime.formatted.longest.taxonomy[992]


#If order is present, use this.

taxonomy$qiime.formatted.longest.taxonomy[994]

for(i in 1: length(taxonomy$qiime.formatted.longest.taxonomy)){
  if(strsplit(taxonomy$qiime.formatted.longest.taxonomy[i], split = ";")[[1]][4] != "o__unidentified" & length(strsplit(taxonomy$qiime.formatted.longest.taxonomy[i], split = ";")[[1]]) == 6){
    order <- strsplit(taxonomy$qiime.formatted.longest.taxonomy[i], split = ";")[[1]][4]
    order <- sub(".*__", "", order)
    taxonomy$qiime.formatted.longest.taxonomy[i] <- paste(taxonomy$qiime.formatted.longest.taxonomy[i], ";s__", order, "_sp", sep = "")
  }
}

taxonomy$qiime.formatted.longest.taxonomy[994]
taxonomy$qiime.formatted.longest.taxonomy[992]

#If class is present, use this.

taxonomy$qiime.formatted.longest.taxonomy[995] #view a deficient taxon

for(i in 1: length(taxonomy$qiime.formatted.longest.taxonomy)){
  if(strsplit(taxonomy$qiime.formatted.longest.taxonomy[i], split = ";")[[1]][3] != "c__unidentified" & length(strsplit(taxonomy$qiime.formatted.longest.taxonomy[i], split = ";")[[1]]) == 6){
    class <- strsplit(taxonomy$qiime.formatted.longest.taxonomy[i], split = ";")[[1]][3]
    class <- sub(".*__", "", class)
    taxonomy$qiime.formatted.longest.taxonomy[i] <- paste(taxonomy$qiime.formatted.longest.taxonomy[i], ";s__", class, "_sp", sep = "")
  }
}

taxonomy$qiime.formatted.longest.taxonomy[995]


#If phylum is present, use this.

taxonomy$qiime.formatted.longest.taxonomy[993]

for(i in 1: length(taxonomy$qiime.formatted.longest.taxonomy)){
  if(strsplit(taxonomy$qiime.formatted.longest.taxonomy[i], split = ";")[[1]][2] != "p__unidentified" & length(strsplit(taxonomy$qiime.formatted.longest.taxonomy[i], split = ";")[[1]]) == 6){
    phylum <- strsplit(taxonomy$qiime.formatted.longest.taxonomy[i], split = ";")[[1]][2]
    phylum <- sub(".*__", "", phylum)
    taxonomy$qiime.formatted.longest.taxonomy[i] <- paste(taxonomy$qiime.formatted.longest.taxonomy[i], ";s__", phylum, "_sp", sep = "")
  }
}

taxonomy$qiime.formatted.longest.taxonomy[993]

#If kingdom is present.

taxonomy$qiime.formatted.longest.taxonomy[1000]

for(i in 1: length(taxonomy$qiime.formatted.longest.taxonomy)){
  if(strsplit(taxonomy$qiime.formatted.longest.taxonomy[i], split = ";")[[1]][1] != "k__unidentified" & length(strsplit(taxonomy$qiime.formatted.longest.taxonomy[i], split = ";")[[1]]) == 6){
    kingdom <- strsplit(taxonomy$qiime.formatted.longest.taxonomy[i], split = ";")[[1]][1]
    kingdom <- sub(".*__", "", kingdom)
    taxonomy$qiime.formatted.longest.taxonomy[i] <- paste(taxonomy$qiime.formatted.longest.taxonomy[i], ";s__", kingdom, "_sp", sep = "")
  }
}

taxonomy$qiime.formatted.longest.taxonomy[1000]

#Check entire taxonomy column for all taxonomy levels.  (All should add to zero.)
sum(grepl("k__", taxonomy$qiime.formatted.longest.taxonomy) != 'TRUE')
sum(grepl("p__", taxonomy$qiime.formatted.longest.taxonomy) != 'TRUE')
sum(grepl("c__", taxonomy$qiime.formatted.longest.taxonomy) != 'TRUE')
sum(grepl("o__", taxonomy$qiime.formatted.longest.taxonomy) != 'TRUE')
sum(grepl("f__", taxonomy$qiime.formatted.longest.taxonomy) != 'TRUE')
sum(grepl("g__", taxonomy$qiime.formatted.longest.taxonomy) != 'TRUE')
sum(grepl("s__", taxonomy$qiime.formatted.longest.taxonomy) != 'TRUE')

#Write to Excel
write.csv(taxonomy, file = '~/Dropbox/Noah_USDA_FungiSCN/MetagenomicSequencing/Bulk.Soil.2015.2016/AMPTK/OTU_Tables/3.30.18/taxonomy.with.longest.taxonomy.Aug.16.2018.csv', row.names = TRUE)
```
