## Supplemental Code for "Fungal Communities in Soybean Cyst Nematode-Infested Soil under Long Term Corn and Soybean Monoculture and Crop Rotation": README.md

# OTU-taxonomy-assignment
This repository contains a script that compares taxonomies assigned to OTUs by different methods (i.e. AMPtk, Consensus BLAST in QIIME2, etc.), addresses conflicts between different methods, and assigns the deepest (i.e species-level when possible) reliable level of taxonomy to OTUs.
