## Supplemental Table 1 for "Fungal Communities in Soybean Cyst Nematode-Infested Soil under Long Term Corn and Soybean Monoculture and Crop Rotation"

**Table S1**. Corn and soybean crop sequences in a long-term crop rotation study site established in 1982 in Waseca, MN. Crop sequences include 5 consecutive years of corn after the 5th year of soybean (C1-C5); 5 consecutive years of soybean after the 5th year of corn (S1-S5); soybean in annual rotation with corn (Sa); corn in annual rotation with soybean (Ca); long-term *Bt* corn (Cc) and SCN-susceptible soybean (Ss) monocultures; non-*Bt* Corn (Cn) and SCN-resistant soybean (Sr) monocultures. Prior to 2010 Cn and Sr plots were planted with *Bt* corn and SCN-susceptible soybean, respectively, since 1982.

| | **Crop sequence** | |
|---|---|---|
| **Treatments** | **2015** | **2016** |
| 1 | C4 | C5 |
| 2 | C3 | C4 |
| 3 | C2 | C3 |
| 4 | C1 | C2 |
| 5 | S5 | C1 |
| 6 | S4 | S5 |
| 7 | S3 | S4 |
| 8 | S2 | S3 |
| 9 | S1 | S2 |
| 10 | C4 | S1 |
| 11 | Cc | Cc |
| 12 | Ss | Ss |
| 13 | Sa | Ca |
| 14 | Ca | Sa |
| 15 | Cn | Cn |
| 16 | Sr | Sr |
