## Supplemental Table 2 for "Fungal Communities in Soybean Cyst Nematode-Infested Soil under Long Term Corn and Soybean Monoculture and Crop Rotation"

**Table S2**. Mock community 5 composition.

| Isolate | DNA added (ng) |
|---|---|
| *Alternaria alternata* | 100 |
| *Arthrobotrys oligospora* | 100 |
| *Aspergillus flavus* | 100 |
| *Aspergillus niger* | 10 |
| *Beauveria bassiana* | 100 |
| *Bipolaris sorokiniana* | 100 |
| *Biscogniauxia nummularia* | 100 |
| *Botrytis cinerea* | 100 |
| *Ceratobasidium sp* | 100 |
| *Cladosporium sp* | 10 |
| *Clavulinopsis fusiformis* | 100 |
| *Clonostachys rosea* | 100 |
| *Colletotrichum graminicola* | 100 |
| *Dactylellina ellipsospora* | 100 |
| *Dactylonectria macrodidyma* | 100 |
| *Exophiala crusticola* | 100 |
| *Fusarium graminearum* | 100 |
| *Fusarium oxysporum* | 100 |
| *Geoglossum difforme* | 100 |
| *Hirsutella minnesotensis* | 100 |
| *Hirsutella rhossiliensis* | 100 |
| *Hirsutella thompsonii* | 100 |
| *Ischnoderma resinosum* | 100 |
| *Lactarius atroviridis* | 100 |
| *Metacordyceps chlamydosporia* | 100 |
| *Mucor circinelloides* | 100 |
| *Orbilia auricolor* | 100 |
| *Phlyctochytrium lagenaria* | 100 |
| *Saccharomyces cerevisiae* | 100 |
| *Scleroderma citrinum* | 100 |
| *Spizellomyces punctatus* | 100 |
| *Suillus granulatus* | 100 |
| *Suillus luteus* | 100 |
| *Tolypocladium inflatum* | 100 |
| *Trichaptum biforme* | 100 |
| *Trichoderma harzianum* | 100 |
