## Supplemental Table 4 for "Fungal Communities in Soybean Cyst Nematode-Infested Soil under Long Term Corn and Soybean Monoculture and Crop Rotation"

**Table S4**. Mock community taxa identified by bioinformatics pipeline. "mk5-S3," "mk5-S117," and "mk5" are mock community samples. "mk5-S117" and "mk5" are subsamples of the same PCR product but were sequenced in separate runs. "mk5-S3" was amplified separately and sequenced in its own run. Taxa in bold are split OTUs.

| | **OTU read counts in mock community samples** | | | | |
|---|---|---|---|---|---|
| **OTU ID** | **mk5-S3** | **mk5-S117** | **mk5** | **Taxonomy assigned by bioinformatics pipeline** | **Equivalent mock community member** |
| OTU156 | 4687 | 0 | 0 | *Spizellomyces_punctatus* | *Spizellomyces punctatus* |
| OTU64 | 4248 | 8878 | 32521 | *Tolypocladium_inflatum* | *Tolypocladium inflatum* |
| OTU81 | 4176 | 0 | 0 | *Phlyctochytrium_lagenaria* | *Phlyctochytrium lagenaria* |
| OTU189 | 3577 | 0 | 107 | *Suillus_luteus* | *Suillus luteus* |
| OTU4 | 3390 | 9658 | 34158 | ***Fusarium_oxysporum*** | ***Fusarium oxysporum*** |
| OTU100 | 3156 | 4405 | 16802 | *Ischnoderma_resinosum* | *Ischnoderma resinosum* |
| OTU393 | 2999 | 0 | 0 | *Saccharomyces_cerevisiae* | *Saccharomyces cerevisiae* |
| OTU62 | 2438 | 152 | 690 | ***Cladosporium_sp*** | ***Cladosporium sp*** |
| OTU50 | 2399 | 5485 | 20009 | *Fusarium_sp* | *Fusarium graminearum* |
| OTU55 | 2397 | 11652 | 39398 | ***Beauveria_bassiana*** | ***Beauveria bassiana*** |
| OTU112 | 1800 | 1521 | 6085 | *Metacordyceps_chlamydosporia* | *Metacordyceps chlamydosporia* |
| OTU9 | 1703 | 3858 | 14033 | ***Alternaria_alternata*** | ***Alternaria alternata*** |
| OTU116 | 1683 | 6685 | 23092 | *Aspergillus_flavus* | *Aspergillus flavus* |
| OTU182 | 1504 | 58 | 246 | *Bipolaris_sorokiniana* | *Bipolaris sorokiniana* |
| OTU269 | 1194 | 2820 | 10274 | ***Exophiala_crusticola*** | ***Exophiala crusticola*** |
| OTU446 | 1171 | 73 | 214 | *Trichaptum_biforme* | *Trichaptum biforme* |
| OTU218 | 941 | 88 | 319 | *Trichoderma_harzianum* | *Trichoderma harzianum* |
| OTU19 | 884 | 182 | 601 | *Clonostachys_sp* | *Clonostachys rosea* |
| OTU54 | 850 | 3069 | 11506 | *Dactylonectria_macrodidyma* | *Dactylonectria macrodidyma* |
| OTU258 | 763 | 0 | 0 | *Arthrobotrys_oligospora* | *Arthrobotrys oligospora* |
| OTU80 | 695 | 1328 | 4839 | *Dactylellina_ellipsospora* | *Dactylellina ellipsospora* |
| OTU272 | 680 | 3001 | 9822 | *Aspergillus_niger* | *Aspergillus niger* |
| OTU214 | 658 | 1768 | 5834 | *Colletotrichum_graminicola* | *Colletotrichum graminicola* |
| OTU207 | 657 | 1538 | 5940 | *Ceratobasidium_sp* | *Ceratobasidium sp* |
| OTU196 | 544 | 201 | 784 | *Botrytis_cinerea* | *Botrytis cinerea* |
| OTU126 | 542 | 2287 | 8668 | *Scleroderma_citrinum* | *Scleroderma citrinum* |
| OTU414 | 413 | 2003 | 6353 | *Hirsutella_minnesotensis* | *Hirsutella minnesotensis* |
| OTU967 | 360 | 3 | 17 | *Suillus_granulatus* | *Suillus granulatus* |
| OTU107 | 340 | 1728 | 5693 | *Hirsutella_thompsonii* | *Hirsutella thompsonii* |
| OTU365 | 303 | 608 | 2177 | *Biscogniauxia_nummularia* | *Biscogniauxia nummularia* |
| OTU443 | 276 | 329 | 1202 | *Lactarius_atroviridis* | *Lactarius atroviridis* |
| OTU620 | 84 | 524 | 1929 | *Geoglossum_sp* | *Geoglossum difforme* |
| OTU1182 | 79 | 117 | 398 | ***Beauveria_bassiana*** | ***Beauveria bassiana*** |
| OTU197 | 68 | 267 | 837 | *Hirsutella_rhossiliensis* | *Hirsutella rhossiliensis* |
| OTU1054 | 34 | 123 | 482 | *Clavulinopsis_sp* | *Clavulinopsis fusiformis* |
| OTU1072 | 17 | 0 | 0 | *Orbilia_auricolor* | *Orbilia auricolor* |
| OTU1554 | 13 | 0 | 0 | *Mucor_sp* | *Mucor circinelloides* |
| OTU2689 | 10 | 4 | 11 | *Phaeocalicium_polyporaeum* | No match |
| OTU5347 | 5 | 2 | 2 | *Trichoderma_velutinum* | No match |
| OTU12482 | 4 | 0 | 0 | *Fungi_sp* | No match |
| OTU974 | 2 | 0 | 0 | *Penicillium_chrysogenum* | No match |
| OTU3076 | 1 | 0 | 0 | *Lecanicillium_fungicola* | No match |
| OTU7925 | 1 | 0 | 0 | ***Cladosporium_sp*** | ***Cladosporium sp*** |
| OTU559 | 0 | 1 | 0 | *Hygrophoraceae_sp* | No match |
| OTU650 | 0 | 0 | 7 | *Plantae_sp* | No match |
| OTU1732 | 0 | 0 | 5 | *Acremonium_sclerotigenum* | No match |
| OTU1821 | 0 | 2 | 4 | *Podospora_sp* | No match |
| OTU3478 | 0 | 0 | 2 | ***Capnodiales_sp*** | ***Cladosporium sp*** |
| OTU3544 | 0 | 15 | 39 | ***Chaetothyriales_sp*** | ***Exophiala crusticola*** |
| OTU4082 | 0 | 0 | 3 | ***Fusarium_oxysporum*** | ***Fusarium oxysporum*** |
| OTU5856 | 0 | 8 | 7 | ***Cantharellales_sp*** | ***Ceratobasidium sp*** |
| OTU6727 | 0 | 0 | 2 | ***Alternaria_alternata*** | ***Alternaria alternata*** |
| OTU7618 | 0 | 0 | 2 | *Devriesia_pseudoamericana* | No match |
| OTU9435 | 0 | 0 | 3 | *Penicillium_sp* | No match |
| Percent taxa detected | 100% | 81% | 83% | | |
