## Supplemental Table 5 for "Fungal Communities in Soybean Cyst Nematode-Infested Soil under Long Term Corn and Soybean Monoculture and Crop Rotation"

**Table S5**. OTUs detected in synthetic mock community sample.

| OTU ID | Number of Reads | Taxonomy |
|---|---|---|
| OTU381 | 13605 | SYN00012 |
| OTU259 | 12169 | SYN00007 |
| OTU347 | 11156 | SYN00011 |
| OTU293 | 9913 | SYN00008 |
| OTU422 | 7149 | SYN00005 |
| OTU457 | 6518 | SYN00006 |
| OTU456 | 4738 | SYN00001 |
| OTU544 | 4706 | SYN00010 |
| OTU523 | 4579 | SYN00009 |
| OTU534 | 4099 | SYN00002 |
| OTU6301 | 13 | *Amanita*_sp |
