## Supplemental Table 6 for "Fungal Communities in Soybean Cyst Nematode-Infested Soil under Long Term Corn and Soybean Monoculture and Crop Rotation"

**Table S6**. Nematophagous guilds.

| Genus | Species | Synonyms | Nematophagous guild |
|---|---|---|---|
| *Clonostachys* | *rosea* | *Gliocladium roseum* | egg_parasite |
| *Cylindrocarpon* | *destructans* | *Ilyonectria destructans* | egg_parasite |
| *Exophiala* | *pisciphila* | | egg_parasite |
| *Fusarium* | *oxysporum* | | egg_parasite |
| *Fusarium* | *solani* | | egg_parasite |
| *Fusarium* | *neocosmosporiellum* | *Neocosmospora vasinfecta* | egg_parasite |
| *Lecanicillium* | *lecanii* | *Verticillium lecanii* | egg_parasite |
| *Metacordyceps* | *chlamydosporia* | | egg_parasite |
| *Phoma* | *macrostoma* | | egg_parasite |
| *Phoma* | *chrysanthemicola* | | egg_parasite |
| *Purpureocillium* | *lilacinum* | *Paecilomyces lilacinus* | egg_parasite |
| *Setophoma* | *terrestris* | *Pyrenochaeta terrestris* | egg_parasite |
| *Stagonospora* | *heteroderae* | | egg_parasite |
| *Catenaria* | *anguillulae* | | endoparasite |
| *Catenaria* | *auxiliaris* | | endoparasite |
| *Drechmeria* | *coniospora* | | endoparasite |
| *Haptocillium* | *balanoides* | | endoparasite |
| *Harposporium* | | | endoparasite |
| *Hirsutella* | *minnesotensis* | | endoparasite |
| *Hirsutella* | *rhossiliensis* | | endoparasite |
| *Hyphochytridium* | *catenoides* | | endoparasite |
| *Verticillium* | *balanoides* | | endoparasite |
| *Arthrobotrys* | | | nematode_trapping |
| *Brachyphoris* | *oviparasitica* | | nematode_trapping |
| *Cystopage* | | | nematode_trapping |
| *Dactylaria* | | | nematode_trapping |
| *Dactylella* | | | nematode_trapping |
| *Dactylellina* | | | nematode_trapping |
| *Drechslerella* | | | nematode_trapping |
| *Gamsylella* | | | nematode_trapping |
| *Hohenbuehelia* | | *Nematoctonus* | nematode_trapping |
| *Monacrosporium* | | | nematode_trapping |
| *Orbilia* | | | nematode_trapping |
| *Stylopage* | | | nematode_trapping |
| *Dactylella* | *oxyspora* | | not_nematode_trapping |
| *Dactylella* | *heptameres* | | not_nematode_trapping |
