## Supplemental Table 7 for "Fungal Communities in Soybean Cyst Nematode-Infested Soil under Long Term Corn and Soybean Monoculture and Crop Rotation"

**Table S7**. Mean values for soil properties within crop sequences. Standard error of the means (SEM) are indicated in parentheses. Variables that were significantly related to crop sequence are indicated with an asterisk, and *P*-values are reported for the Crop Sequence term in the ANOVA on the model, Soil Property ~ Block + Crop Sequence. $R^2$ and *P*-values are reported for univariate adonis using the formula, Community Distance ~ Soil Property. All reported *P*-values have been corrected using the false discovery rate (FDR) procedure. Significant *P*-values ($P < 0.05$) and adonis $R^2$ values greater than 0.1 that are associated with significant *P*-values are in bold. Crop sequences include 5 consecutive years of corn after the 5th year of soybean (C1-C5); 5 consecutive years of soybean after the 5th year of corn (S1-S5); soybean in annual rotation with corn (Sa); corn in annual rotation with soybean (Ca); long-term *Bt* corn (Cc) and SCN-susceptible soybean (Ss) monocultures; non-*Bt* Corn (Cn) and SCN-resistant soybean (Sr) monocultures.

| | **2015** | | | | | | | | | | | |
|---|---|---|---|---|---|---|---|---|---|---|---|---|
| | Crop Sequence | P, ppm | K, ppm | %OM | pH | Fe, mg kg-1 | Mn, mg kg-1 | Zn, mg kg-1 | Cu, mg kg-1 | Total N, %N | TOC, %C | C:N |
| | C1 | 24 (3.4) | 147 (17) | 4.8 (0.5) | 6.4 (0.4) | 63 (18) | 49 (9) | 2.3 (0.16) | 1.3 (0.11) | 0.17 (0.016) | 2.3 (0.21) | 13.4 (0.42) |
| | C2 | 21 (2.7) | 145 (8) | 5.3 (0.1) | 6.7 (0.2) | 36 (14) | 41 (7) | 1.9 (0.14) | 1.2 (0.07) | 0.18 (0.009) | 2.4 (0.13) | 13.5 (0.14) |
| | C3 | 19 (1.8) | 150 (5) | 5.1 (0.2) | 6.5 (0.3) | 54 (20) | 48 (9) | 2.1 (0.17) | 1.3 (0.09) | 0.18 (0.004) | 2.3 (0.1) | 12.8 (0.58) |
| | C4 | 22 (2.4) | 147 (8) | 5.6 (0.3) | 6.2 (0.1) | 70 (12) | 58 (7) | 2.0 (0.07) | 1.3 (0.16) | 0.20 (0.01) | 2.7 (0.14) | 13.2 (0.08) |
| | C5 | 17 (3.3) | 141 (8) | 5.1 (0.3) | 6.4 (0.3) | 57 (21) | 51 (8) | 2.3 (0.27) | 1.4 (0.14) | 0.17 (0.009) | 2.3 (0.16) | 13.4 (0.29) |
| | Cc | 16 (1.8) | 140 (3) | 5.4 (0.3) | 5.8 (0.3) | 99 (23) | 73 (8) | 2.3 (0.18) | 1.5 (0.09) | 0.19 (0.004) | 2.5 (0.09) | 13.1 (0.24) |
| | Cn | 18 (1.8) | 143 (8) | 5.7 (0.4) | 5.8 (0.2) | 95 (21) | 67 (6) | 2.3 (0.32) | 1.5 (0.08) | 0.21 (0.012) | 2.7 (0.15) | 13.3 (0.06) |
| | S1 | 23 (1.7) | 151 (6) | 5.4 (0.1) | 6.0 (0.1) | 78 (9) | 64 (4) | 2.2 (0.3) | 1.4 (0.11) | 0.20 (0.007) | 2.6 (0.1) | 13.2 (0.08) |
| | S2 | 19 (3.3) | 147 (6) | 5.4 (0.5) | 6.7 (0.3) | 37 (10) | 46 (10) | 1.9 (0.21) | 1.3 (0.13) | 0.18 (0.016) | 2.4 (0.21) | 13.6 (0.13) |
| | S3 | 26 (2.8) | 152 (9) | 5.6 (0.4) | 6.1 (0.1) | 65 (12) | 56 (3) | 2.7 (0.48) | 1.3 (0.05) | 0.19 (0.013) | 2.6 (0.2) | 13.4 (0.22) |
| | S4 | 22 (3.1) | 160 (22) | 5.3 (0.4) | 6.2 (0.2) | 66 (15) | 57 (3) | 2.3 (0.12) | 1.5 (0.07) | 0.17 (0.006) | 2.3 (0.07) | 13.0 (0.04) |
| | S5 | 24 (3.5) | 145 (13) | 5.3 (0.2) | 6.1 (0.2) | 78 (20) | 55 (2) | 2.6 (0.21) | 1.4 (0.12) | 0.19 (0.007) | 2.5 (0.09) | 13.2 (0.05) |
| | Ss | 32 (3.9) | 142 (4) | 4.9 (0.2) | 6.7 (0.1) | 32 (6) | 37 (3) | 2.5 (0.3) | 1.2 (0.04) | 0.17 (0.011) | 2.3 (0.14) | 13.5 (0.35) |
| | Sr | 39 (4.1) | 132 (8) | 5.2 (0.2) | 6.5 (0.1) | 45 (11) | 43 (6) | 1.8 (0.09) | 1.1 (0.07) | 0.18 (0.004) | 2.4 (0.04) | 13.4 (0.14) |
| | Ca | 20 (0.9) | 152 (2) | 5.7 (0.2) | 6.3 (0.2) | 70 (21) | 54 (8) | 2.5 (0.18) | 1.4 (0.04) | 0.20 (0.007) | 2.7 (0.12) | 13.4 (0.13) |
| | Sa | 20 (3.9) | 129 (5) | 5.5 (0.4) | 6.5 (0.1) | 46 (12) | 51 (3) | 2.0 (0.04) | 1.3 (0.08) | 0.19 (0.012) | 2.6 (0.19) | 13.6 (0.16) |
| ANOVA | *P*-value | **0.002** | 0.73 | 0.43 | 0.17 | 0.15 | **0.038** | 0.32 | **0.038** | 0.1 | 0.1 | 0.73 |
| Adonis | $R^2$ | 0.049 | 0.023 | 0.029 | **0.12** | **0.12** | **0.12** | 0.017 | 0.078 | 0.037 | 0.03 | 0.031 |
| Adonis | *P*-value | **0.002** | 0.058 | **0.019** | **0.002** | **0.002** | **0.002** | 0.26 | **0.002** | **0.0034** | **0.012** | **0.013** |
| | **2016** | | | | | | | | | | | |
| | Crop Sequence | P, ppm | K, ppm | %OM | pH | Fe, mg kg-1 | Mn, mg kg-1 | Zn, mg kg-1 | Cu, mg kg-1 | Total N, %N | TOC, %C | C:N |
| | C1 | 20 (3.2) | 135 (14) | 5.4 (0.2) | 6.4 (0.2) | 94 (22) | 47 (2) | 5.6 (0.66) | 1.5 (0.1) | 0.19 (0.008) | 2.5 (0.13) | 13.2 (0.15) |
| | C2 | 16 (3.4) | 148 (15) | 5 (0.5) | 6.7 (0.4) | 86 (26) | 40 (8) | 7.2 (1.3) | 1.4 (0.16) | 0.17 (0.016) | 2.3 (0.22) | 13.6 (0.35) |
| | C3 | 12 (2.3) | 132 (3) | 5.5 (0.09) | 6.8 (0.2) | 54 (18) | 41 (5) | 5.4 (0.8) | 1.4 (0.07) | 0.18 (0.004) | 2.4 (0.07) | 13.4 (0.09) |
| | C4 | 14 (2.1) | 144 (9) | 5.4 (0.2) | 6.6 (0.3) | 76 (28) | 43 (7) | 6.1 (1.7) | 1.5 (0.12) | 0.18 (0.006) | 2.5 (0.11) | 13.3 (0.2) |
| | C5 | 16 (2.4) | 132 (10) | 5.6 (0.1) | 6.3 (0.1) | 96 (12) | 46 (2) | 7.5 (1.1) | 1.4 (0.09) | 0.19 (0.005) | 2.6 (0.08) | 13.5 (0.13) |
| | Cc | 8 (0.9) | 140 (10) | 5.8 (0.4) | 6.1 (0.3) | 115 (30) | 60 (4) | 6.7 (2.2) | 1.7 (0.1) | 0.20 (0.01) | 2.7 (0.2) | 13.3 (0.29) |
| | Cn | 11 (1.7) | 132 (8) | 5.8 (0.4) | 6.0 (0.2) | 119 (23) | 60 (3) | 6.6 (1.1) | 1.7 (0.11) | 0.20 (0.013) | 2.7 (0.2) | 13.4 (0.18) |
| | S1 | 11 (2.8) | 126 (14) | 5.3 (0.3) | 6.6 (0.3) | 73 (24) | 42 (4) | 7.2 (1.8) | 1.5 (0.13) | 0.18 (0.008) | 2.4 (0.14) | 13.2 (0.22) |
| | S2 | 16 (2.1) | 134 (7) | 5.6 (0.03) | 6.2 (0.1) | 97 (9) | 48 (2) | 5 (0.57) | 1.5 (0.1) | 0.19 (0.003) | 2.6 (0.05) | 13.5 (0.15) |
| | S3 | 13 (2.9) | 131 (10) | 5.5 (0.5) | 6.9 (0.3) | 47 (13) | 37 (7) | 4.3 (0.54) | 1.3 (0.11) | 0.18 (0.015) | 2.5 (0.21) | 13.7 (0.07) |
| | S4 | 16 (1.2) | 121 (7) | 5.6 (0.3) | 6.5 (0.1) | 81 (10) | 47 (2) | 6 (0.7) | 1.4 (0.07) | 0.19 (0.009) | 2.5 (0.17) | 13.4 (0.28) |
| | S5 | 15 (2.5) | 139 (15) | 5.4 (0.4) | 6.5 (0.2) | 80 (18) | 45 (4) | 5.3 (0.29) | 1.5 (0.11) | 0.18 (0.014) | 2.4 (0.2) | 13.4 (0.2) |
| | Ss | 28 (4) | 126 (2) | 5.1 (0.2) | 6.9 (0.1) | 38 (6) | 30 (1) | 4.9 (0.5) | 1.2 (0.05) | 0.17 (0.009) | 2.2 (0.12) | 13.3 (0.19) |
| | Sr | 35 (4.7) | 125 (2) | 5.4 (0.2) | 6.8 (0.1) | 58 (11) | 37 (3) | 7 (1.1) | 1.2 (0.05) | 0.18 (0.004) | 2.5 (0.07) | 13.7 (0.12) |
| | Ca | 16 (4.7) | 124 (3) | 5.7 (0.4) | 6.6 (0.1) | 61 (14) | 45 (4) | 5 (0.25) | 1.4 (0.07) | 0.19 (0.013) | 2.6 (0.2) | 13.6 (0.15) |
| | Sa | 14 (1.2) | 140 (7) | 5.8 (0.2) | 6.5 (0.3) | 86 (24) | 44 (4) | 6.2 (1.2) | 1.5 (0.03) | 0.20 (0.006) | 2.8 (0.17) | 14.0 (0.51) |
| ANOVA | *P*-value | **5.00E-05** | 0.5 | 0.64 | 0.3 | 0.3 | **0.0052** | 0.64 | **0.0039** | 0.34 | 0.4 | 0.59 |
| Adonis | $R^2$ | 0.05 | 0.03 | 0.023 | **0.13** | **0.14** | **0.11** | 0.019 | 0.079 | 0.027 | 0.023 | 0.036 |
| Adonis | *P*-value | **0.002** | **0.012** | **0.071** | **0.002** | **0.002** | **0.002** | 0.15 | **0.002** | **0.037** | 0.071 | **0.002** |
