## Supplemental Table 8 for "Fungal Communities in Soybean Cyst Nematode-Infested Soil under Long Term Corn and Soybean Monoculture and Crop Rotation"

**Table S8.** Correlations between soil properties and corn and soybean yields. Bold type indicates significant FDR-corrected *P*-values ($P < 0.05$) and associated $R^2$ values and linear equations. Significant *P*-values are reported at $*P < 0.05$, $**P < 0.01$, and $***P < 0.001$.

| **2015** | | | | | | |
|---|---|---|---|---|---|---|
| **Corn** | | | | **Soybean** | | |
| **Soil Property** | **Equation** | $R^2$ | ***P*-value** | **Equation** | $R^2$ | ***P*-value** |
| P | Y = 9.9 + 0.05X | 0.02 | 0.3 | Y = 4.1 - 0.0015X | 0.0013 | 0.7 |
| K | Y = 12 - 0.0098X | 0.0065 | 0.6 | Y = 3.8 + 0.001X | 0.01 | 0.4 |
| OM | Y = 11 - 0.057X | 0.00032 | 0.9 | **Y = 2.8 + 0.2X** | **0.24** | **0.00003*** ** |
| pH | **Y = 5 + 0.9X** | **0.079** | **0.02*** | **Y = 5.8 - 0.27X** | **0.14** | **0.001**** |
| Fe | **Y = 12 - 0.012X** | **0.062** | **0.04*** | Y = 3.9 + 0.002X | 0.036 | 0.1 |
| Mn | **Y = 13 - 0.033X** | **0.09** | **0.02*** | **Y = 3.5 + 0.01X** | **0.15** | **0.0007*** ** |
| Zn | **Y = 7.1 + 2X** | **0.11** | **0.01*** | **Y = 3.7 + 0.2X** | **0.087** | **0.01*** |
| Cu | **Y = 15 - 2.7X** | **0.088** | **0.02*** | **Y = 3.4 + 0.5X** | **0.1** | **0.006**** |
| Total N | Y = 12 - 7.4X | 0.006 | 0.6 | **Y = 2.9 + 6X** | **0.2** | **0.00009*** ** |
| TOC | Y = 10 + 0.3X | 0.0026 | 0.7 | **Y = 3 + 0.4X** | **0.19** | **0.0002*** ** |
| C:N | **Y = -6.9 + 1X** | **0.18** | **0.0005*** ** | Y = 5 - 0.074X | 0.0083 | 0.5 |
| **2016** | | | | | | |
| **Corn** | | | | **Soybean** | | |
| **Soil Property** | **Equation** | $R^2$ | ***P*-value** | **Equation** | $R^2$ | ***P*-value** |
| P | Y = 11 + 0.04X | 0.035 | 0.2 | **Y = 4 - 0.011X** | **0.072** | **0.02*** |
| K | **Y = 15 - 0.025X** | **0.11** | **0.009**** | **Y = 3.2 + 0.004X** | **0.071** | **0.02*** |
| OM | **Y = 6.7 + 0.9X** | **0.14** | **0.006**** | **Y = 2.7 + 0.2X** | **0.16** | **0.0005*** ** |
| pH | Y = 11 + 0.1X | 0.0013 | 0.8 | **Y = 5.1 - 0.21X** | **0.091** | **0.01*** |
| Fe | Y = 12 - 0.0052X | 0.026 | 0.2 | Y = 3.7 + 0.001X | 0.021 | 0.2 |
| Mn | Y = 11 + 0.008X | 0.0037 | 0.7 | **Y = 3.3 + 0.01X** | **0.16** | **0.0005*** ** |
| Zn | Y = 12 + 0.006X | 0.000095 | 0.9 | Y = 3.8 + 0.001X | 0.000097 | 0.9 |
| Cu | Y = 12 - 0.49X | 0.0053 | 0.7 | **Y = 2.8 + 0.7X** | **0.23** | **0.00003*** ** |
| Total N | **Y = 7 + 25X** | **0.11** | **0.009**** | **Y = 2.5 + 7X** | **0.23** | **0.00003*** ** |
| TOC | **Y = 7.8 + 2X** | **0.1** | **0.01*** | **Y = 2.9 + 0.3X** | **0.15** | **0.0005*** ** |
| C:N | Y = 5.9 + 0.4X | 0.014 | 0.4 | Y = 3.9 - 0.0074X | 0.00019 | 0.9 |
