## Supplemental Table 9 for "Fungal Communities in Soybean Cyst Nematode-Infested Soil under Long Term Corn and Soybean Monoculture and Crop Rotation"

**Table S9.** Alpha diversity metrics based on the model, Diversity ~ Block + Season + Host + Season:Host. Chao1 is total alpha diversity of bulk soil fungal communities. Diversity metrics are also reported for individual nematophagous guilds and for trophic modes. Only OTUs assigned a trophic mode with a confidence ranking of "Probable" or "Highly Probable" were used for this analysis. Significant *P*-values associated with ANOVA are reported at $*P < 0.05$, $**P < 0.01$, and $***P < 0.001$. Different lowercase letters indicate significant differences in means according to Tukey's honestly significant difference (HSD) test.

| | | | | | | | | |
|---|---|---|---|---|---|---|---|---|
| **2015** | | | | | | | | |
| **ANOVA** | | **Chao1** | **Saprotrophs** | **Pathotrophs** | **Symbiotrophs** | **Trapping fungi** | **Egg parasites** | **Endoparasites** |
| | Season | <0.001*** | <0.001*** | <0.001*** | <0.001*** | <0.001*** | 0.02* | 0.66 |
| | Host | 0.012* | 0.03* | 0.36 | <0.001*** | 0.3 | <0.001*** | 0.46 |
| | Season:Host | 0.58 | 0.79 | 0.8 | <0.001*** | 0.09 | 0.38 | 0.62 |
| **Host** | | | | | | | | |
| | Corn | 797a | 91a | 24a | 55a | 3.18a | 5.32b | 0.53a |
| | Soy | 725b | 85b | 24a | 34b | 3.44a | 6.11a | 0.46a |
| **2016** | | | | | | | | |
| **ANOVA** | | **Chao1** | **Saprotrophs** | **Pathotrophs** | **Symbiotrophs** | **Trapping fungi** | **Egg parasites** | **Endoparasites** |
| | Season | <0.001*** | <0.001*** | <0.001*** | <0.001*** | 0.08. | 0.40 | 0.26 |
| | Host | 0.0063** | 0.03* | 0.68 | <0.001*** | 0.0027** | 0.02* | 0.15 |
| | Season:Host | 0.14 | 0.3 | 0.19 | 0.06 | 0.91 | 0.85 | 0.56 |
| **Host** | | | | | | | | |
| | Corn | 1088a | 111a | 29a | 74a | 3.41b | 5.46b | 0.52a |
| | Soy | 1011b | 106b | 30a | 52b | 4.27a | 5.93a | 0.4a |
