## Supplemental Table 10 for "Fungal Communities in Soybean Cyst Nematode-Infested Soil under Long Term Corn and Soybean Monoculture and Crop Rotation"

**Table S10**. Rho values associated with Spearman correlation tests between total alpha diversity of bulk soil fungal communities (Chao1), alpha diversity of trophic modes, and alpha diversity of nematophagous guilds, and SCN density, monoculture year, and yield of corn and soybeans. Significant correlations are reported at false-discovery rate (FDR)-corrected *P*-values of *$P < 0.05$, **$P < 0.01$, and ***$P < 0.001$

| Test | SeasonYear | Chao1 | Trapping fungi | Egg Parasites | Endoparasites | Saprotroph | Pathotroph | Symbiotroph |
|---|---|---|---|---|---|---|---|---|
| **SCN** | Fall 2015 | 0.0028 | 0.058 | 0.2 | -0.026 | 0.046 | 0.03 | -0.02 |
| **SCN** | Mid 2015 | -0.19 | 0.25 | 0.21 | 0.1 | -0.27 | -0.1 | -0.32 |
| **SCN** | Spring 2015 | -0.13 | 0.3* | 0.3* | 0.038 | -0.12 | -0.037 | -0.3 |
| **SCN** | Fall 2016 | -0.33* | 0.025 | -0.015 | -0.14 | -0.16 | -0.092 | -0.4** |
| **SCN** | Mid 2016 | -0.25 | 0.15 | 0.2 | -0.091 | -0.26 | 0.00068 | -0.23 |
| **SCN** | Spring 2016 | -0.093 | 0.3* | 0.33* | -0.35* | -0.054 | -0.037 | -0.25 |
| **Soy Year** | Fall 2015 | 0.12 | 0.2 | 0.018 | 0.63** | 0.13 | 0.28 | 0.2 |
| **Soy Year** | Mid 2015 | -0.33 | 0.14 | 0.089 | -0.058 | -0.4 | -0.11 | -0.49* |
| **Soy Year** | Spring 2015 | -0.28 | 0.38 | 0.43 | 0.3 | -0.2 | -0.051 | -0.4 |
| **Soy Year** | Fall 2016 | -0.42 | 0.45 | 0.27 | -0.0064 | -0.31 | -0.18 | -0.39 |
| **Soy Year** | Mid 2016 | -0.15 | 0.47 | 0.26 | -0.092 | -0.33 | 0.11 | -0.36 |
| **Soy Year** | Spring 2016 | -0.19 | 0.33 | 0.072 | 0.085 | -0.36 | 0.16 | -0.7*** |
| **Corn Year** | Fall 2015 | 0.068 | -0.089 | -0.14 | 0.1 | 0.079 | 0.1 | 0.18 |
| **Corn Year** | Mid 2015 | 0.18 | -0.54* | -0.33 | -0.11 | 0.26 | 0.044 | 0.34 |
| **Corn Year** | Spring 2015 | 0.12 | -0.31 | -0.55* | -0.34 | 0.27 | -0.22 | 0.52 |
| **Corn Year** | Fall 2016 | 0.031 | -0.3 | -0.4 | -0.13 | -0.044 | -0.4 | 0.0028 |
| **Corn Year** | Mid 2016 | 0.0089 | -0.65** | -0.59** | 0.24 | 0.086 | -0.27 | -0.012 |
| **Corn Year** | Spring 2016 | 0.28 | -0.46* | -0.59** | 0.51* | 0.15 | 0.1 | 0.5* |
| **Soy Yield** | Fall 2015 | 0.24 | 0.22 | 0.018 | -0.1 | 0.12 | 0.12 | 0.25 |
| **Soy Yield** | Mid 2015 | 0.16 | -0.12 | 0.092 | 0.025 | 0.076 | 0.2 | 0.039 |
| **Soy Yield** | Spring 2015 | 0.087 | -0.0063 | 0.25 | 0.051 | 0.099 | 0.062 | -0.093 |
| **Soy Yield** | Fall 2016 | 0.27 | 0.15 | -0.11 | 0.35 | 0.2 | 0.32 | 0.21 |
| **Soy Yield** | Mid 2016 | -0.043 | -0.062 | -0.038 | 0.28 | -0.16 | 0.16 | 0.02 |
| **Soy Yield** | Spring 2016 | -0.03 | -0.18 | -0.21 | 0.18 | 0.049 | -0.04 | -0.0013 |
| **Corn Yield** | Fall 2015 | 0.068 | -0.13 | 0.28 | 0.0016 | -0.024 | 0.15 | 0.018 |
| **Corn Yield** | Mid 2015 | -0.24 | 0.31 | -0.28 | 0.091 | -0.45 | -0.095 | -0.27 |
| **Corn Yield** | Spring 2015 | 0.042 | 0.54** | 0.34 | -0.15 | 0.13 | -0.019 | -0.043 |
| **Corn Yield** | Fall 2016 | -0.087 | 0.12 | 0.25 | 0.079 | 0.18 | 0.18 | -0.071 |
| **Corn Yield** | Mid 2016 | -0.086 | 0.37 | 0.38 | 0.031 | -0.045 | 0.12 | -0.19 |
| **Corn Yield** | Spring 2016 | -0.53* | 0.072 | 0.39 | -0.07 | -0.34 | -0.095 | -0.55** |
